# A single-nucleus transcriptomic atlas of the adult *Aedes aegypti* mosquito

**DOI:** 10.1101/2025.02.25.639765

**Authors:** Olivia V. Goldman, Alexandra E. DeFoe, Yanyan Qi, Yaoyu Jiao, Shih-Che Weng, Brittney Wick, Leah Houri-Zeevi, Priyanka Lakhiani, Takeshi Morita, Jacopo Razzauti, Adriana Rosas-Villegas, Yael N. Tsitohay, Madison M. Walker, Ben R. Hopkins, Aedes aegypti Mosquito Cell Atlas Consortium^, Maximilian Haeussler, Omar S. Akbari, Laura B. Duvall, Helen White-Cooper, Trevor R. Sorrells, Roshan Sharma, Hongjie Li, Leslie B. Vosshall, Nadav Shai

## Abstract

The female *Aedes aegypti* mosquito’s remarkable ability to hunt humans and transmit pathogens relies on her unique biology. Here, we present the *Aedes aegypti* Mosquito Cell Atlas, a comprehensive single-nucleus RNA sequencing dataset of more than 367,000 nuclei from 19 dissected tissues of adult female and male *Aedes aegypti*, providing cellular-level resolution of mosquito biology. We identify novel cell types and expand our understanding of sensory neuron organization of chemoreceptors to all sensory tissues. Our analysis uncovers male-specific cells and sexually dimorphic gene expression in the antenna and brain. In female mosquitoes, we find that glial cells in the brain, rather than neurons, undergo the most extensive transcriptional changes following blood feeding. Our findings provide insights into the cellular basis of mosquito behavior and sexual dimorphism. The *Aedes aegypti* Mosquito Cell Atlas resource enables systematic investigation of cell type-specific expression across all mosquito tissues.

## Introduction

Mosquito-borne diseases affect hundreds of millions of people worldwide, with rising infection rates each year^1,2^. By 2050, climate change-driven habitat expansion is predicted to put nearly half of the world’s population at risk for viral infection from *Aedes* mosquitoes^3^. *Aedes aegypti* is the primary vector for mosquito-borne viruses, including dengue, Zika, yellow fever, and chikungunya^4,5^. Management of mosquito vector populations, the most effective strategy for controlling mosquito-borne disease, has historically relied on insecticides although gene drive strategies are being developed^6^. Deeper insight into mosquito biology is needed to develop additional control methods.

The unique sexual dimorphism of *Aedes aegypti* mosquitoes is fundamental to the threat they pose to public health. Mosquitoes are attracted to human cues, including exhaled carbon dioxide (CO_2_), body heat, and skin odor^7–10^. Only females feed on blood, which contains proteins and other nutrients required for egg production. Humans are the preferred host for female *Aedes aegypti*, contributing to their effectiveness as a disease vector^11,12^. After a blood meal, females undergo physiological and behavioral changes, including suppressed host seeking and generally reduced activity for 48-72 hours while they develop their eggs and find a suitable oviposition site, guided by sensory attraction to freshwater^13–17^. While female mosquitoes have evolved specialized behavioral and reproductive mechanisms for host seeking, blood feeding, finding freshwater for egg laying, and egg development, males have a simpler behavioral repertoire focused on nectar feeding and mating.

Single-cell RNA sequencing (scRNA-seq) and atlasing have been instrumental in defining the molecular identity of known cell types and discovering new cell types. Cell atlases have been constructed for *Drosophila melanogaster*^18^, *Caenorhabditis elegans*^19,20^, *Schmidtea mediterranea*^21^, *Mus musculus*^22,23^, *Microcebus murinus*^24^, and others. These have been key resources for understanding cell type diversity and gene expression patterns.

Prior studies have used bulk RNA sequencing (RNA-seq) to profile diverse *Aedes aegypti* tissues^25–31^. Recently, scRNA-seq and single-nucleus RNA sequencing (snRNA-seq) have been used to profile mosquito tissues such as the *Aedes aegypti* gut^32–35^, olfactory organs^36,37^, brain^38^, fat body^35^, and larval ventral nerve cord^39^, the *Anopheles gambiae* testes^40,41^ and immune system^42,43^, and the *Culex tarsalis* gut^44^. Additional immune system studies have compared hemocytes across both *Anopheles gambiae* and *Aedes aegypti*^42^. The major mosquito vector genera for human disease (*Aedes, Anopheles*, and *Culex*) diverged approximately 110-180 million years ago^45^, representing substantial evolutionary distance important for interpreting comparisons of cellular and molecular findings across species. While previous studies have provided valuable insights into mosquito biology, most single-cell studies focused on specific tissues or cell types, primarily in females. A global gene expression map spanning multiple tissues and both sexes within a single species is needed to enable deeper investigation and uncover unique insights into *Aedes aegypti* biology^18,46^.

We sought to gain system-level insights into the molecular and cellular differences underlying the extraordinary sexual dimorphism of this species. To achieve this, we developed the *Aedes aegypti* Mosquito Cell Atlas, a large-scale snRNA-seq project characterizing every major tissue from the adult female and male *Aedes aegypti* mosquitoes. We profiled 367,096 nuclei from 19 tissues, providing cellular resolution of the entire mosquito transcriptome. For the female brain, we include multiple timepoints before and after blood feeding to investigate transcriptional changes correlated with behavioral shifts linked to reproductive state. We found specialized gene expression patterns and identified antimicrobial peptide-expressing cells in female salivary glands. In the antennae, we discovered male-specific *ppk317*-expressing cells and sexually dimorphic olfactory sensory neurons. We observe that mosquito legs and proboscis house polymodal sensory neurons that co-express receptors for different sensory modalities and across gene families, as shown previously in the antenna ^36,37^. In the brain, we identified sexually dimorphic gene expression in Kenyon cells and extensive transcriptional changes in glial cells following blood feeding.

This atlas represents a valuable resource for the vector biology community, bridging the gap between model organism studies and mosquito-specific biology. We hope the *Aedes aegypti* Mosquito Cell Atlas will encourage comparative studies to further understand the mosquito’s unique biology. While a century of *Drosophila melanogaster* research has provided foundational knowledge of insects, creating the tools and datasets directly related to mosquitoes allows us to move away from homology-based research that seeks to align mosquito and fly biology. More broadly, these data offer new avenues for studying the molecular biology underlying the specific adaptations and specializations that make mosquitoes such effective and deadly pathogen vectors.

## Results

### snRNA-seq atlasing of the adult female and male *Aedes aegypti* mosquito

Tissues for the *Aedes aegypti* Mosquito Cell Atlas were selected based on biological importance and physical dissection feasibility, aiming to map all female and male cell types from aged-matched, sugar-fed animals. We also collected female brains at various times after blood feeding to capture gene expression changes across the reproductive cycle. We collected major body segments (head, thorax, and abdomen) to allow for verification of gene expression signatures and collection of cells and tissues that were not separately dissected. Nineteen tissues were selected across five biological themes: (I) major body segments, (II) sensation and host seeking, (III) viral infection, (IV) reproduction, and (V) central nervous system (Figure 1A). Given the difficulty in isolating intact cells from cuticular tissues such as antennae and maxillary palps^18,36^, we used snRNA-seq rather than scRNA-seq for its demonstrated consistency when applied across tissues^47^ and in representing *in vivo* compositions of insect cell types^48^.

**Figure 1.**
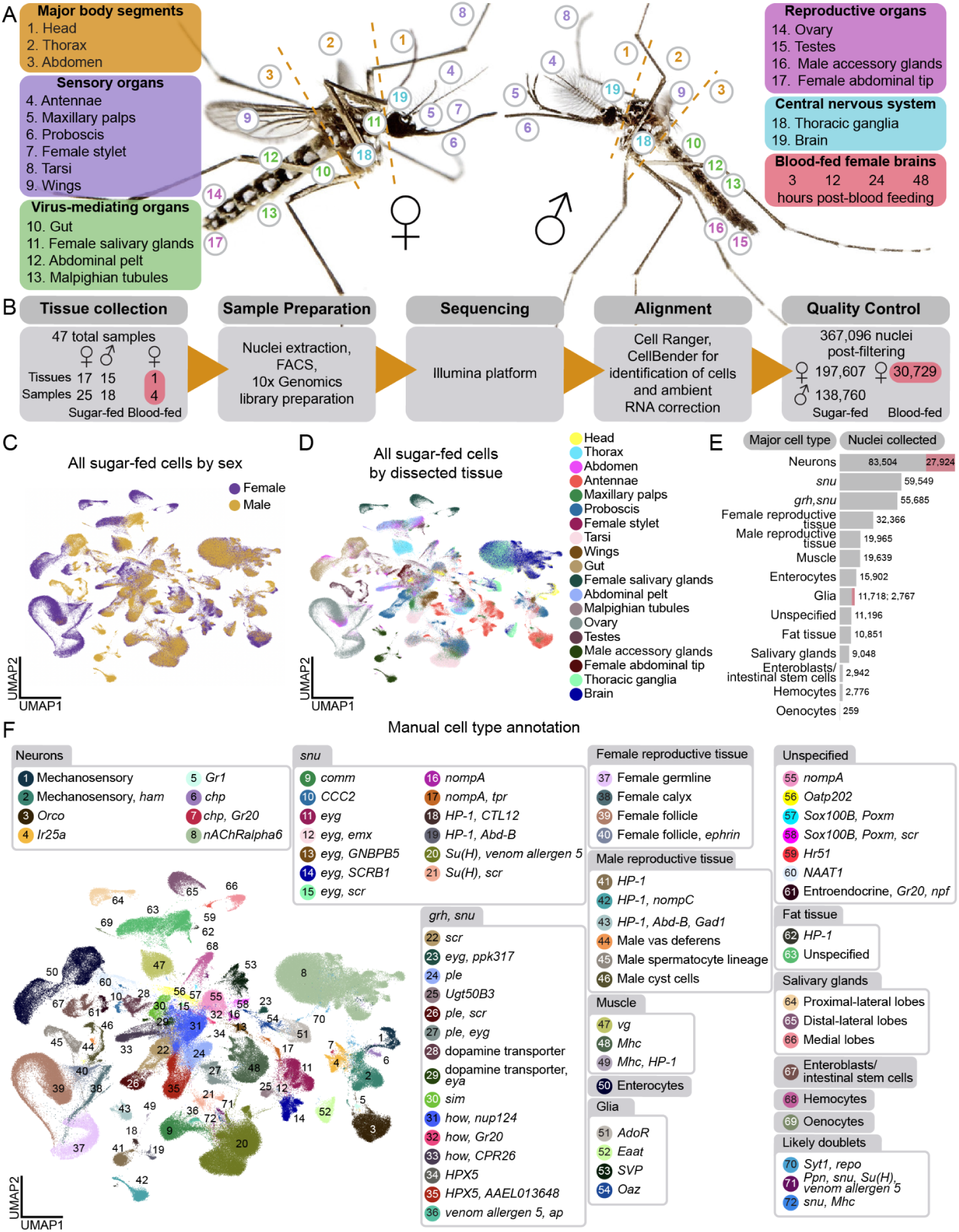
*Aedes aegypti* Mosquito Cell Atlas tissues and data. (A) Photos of *Aedes aegypti* female (left) and male (right). Numbers indicate location of collected tissues (listed in legend boxes). Photos by Alex Wild. (B) Schematic of *Aedes aegypti* Mosquito Cell Atlas workflow. A sample represents an individual library prepared with 10x Genomics commercial kits. Sample counts shown for each sex. Tissues underwent nuclei extraction followed by fluorescence-activated cell sorting (FACS), single-nucleus RNA libraries preparation with 10x Genomics commercial kits, then sequenced using the Illumina platform unless stated otherwise. Raw sequencing data was aligned with Cell Ranger^63^. Cells were identified and ambient RNA removed using CellBender^64^. Samples were individually processed for cell quality control filtering, yielding 367,096 high-quality nuclei (nuclei counts shown for each sex and for blood-feeding conditions). **(C-D)** Uniform manifold approximation and projection (UMAP) for dimension reduction of 330,364 nuclei from mated, sugar-fed samples colored by sex (C) and dissected tissue (D). Blood-fed samples excluded. For batch processing consistency, testes sample underwent ambient RNA removal and cell identification with CellBender (spermatids removed). **(E)** Nuclei counts from mated, sugar-fed (grey) and mated, blood-fed (red) samples for each major cell type, sorted by abundance. **(F)** UMAP of nuclei from all mated, sugar-fed samples, colored and numbered by manual annotation of 66 distinct cell types (“Level 2” annotations, legend) and major cell type categories (“Level 1”, grey headers) (Figure S2B). For more information on annotation, see Table S1.

Female mosquitoes require a blood meal for egg development and suppress host-seeking and biting behavior for several days after a blood meal until the eggs are laid^13–17^. Bulk RNA-seq studies identified hundreds of gene expression changes associated with blood feeding in many tissues, including the brain^26,28,49–52^. To resolve these changes at single-cell resolution, we sequenced female brains at 3, 12, 24, and 48 hours after blood feeding (Figure 1A), spanning key stages of egg maturation and suppressed host attraction.

We dissected 44 samples: 17 sugar-fed female tissues, 15 sugar-fed male tissues, and 4 samples of brains from blood-fed females at the defined timepoints. Male and female animals were co-housed and females were presumably mated prior to dissection. Extracted nuclei were collected using fluorescence-activated cell sorting (FACS), and single-nucleus transcriptomes were generated using 10x Genomics technology and Illumina sequencing unless otherwise stated (Figure 1B). Because data collection methods were identical, we also re-analyzed female antenna and maxillary palp data from Herre, Goldman et al.^36^. All samples were aligned to the *Aedes aegypti* L5 genome^53^ and quality-filtered (Figure S1 and Data S1), yielding 367,096 nuclei: 197,607 from sugar-fed females, 138,760 from sugar-fed males, and 30,729 from blood-fed females. Nuclei were collected from 9,651 mosquitoes across 47 samples (10x Genomics libraries) (44 new, 3 from our previous study^36^ (Figure 1B and Table S1).

Median UMI/nucleus was 3,424, and median genes/nucleus was 1,296, with low average mitochondrial gene content (0.13%) (Figure S1). We analyzed male and female data of the same tissue to compare cell composition and gene expression between the sexes (Data S2 and Zenodo Supplemental Data). We then combined the data from all male and female sugar-fed tissues to create a complete *Aedes aegypti* mosquito cell atlas (Figures 1C-1D).

The hallmark of a cell atlas is the ability to annotate distinct cell types, which is especially challenging in non-model organisms lacking well-defined markers. We developed two complementary strategies to address these challenges. First, we relied on experts in mosquito biology and entomology to annotate data using known *Aedes aegypti* gene markers wherever possible. Second, we computationally identified gene markers using standard scRNA-seq differential gene expression tools^54,55^. When marker genes were uncharacterized, we referenced *Drosophila melanogaster* orthologs via Ensembl Metazoa BioMart^56^, BLAST^57^, or VectorBase^58^. Many of our annotations use marker genes that may imply function based on *Drosophila melanogaster* literature (see Table S1 for gene identifiers and ortholog names). *Aedes aegypti* and *Drosophila melanogaster* are separated by 260 million years of evolution^59–61^, with distinct behaviors, life cycles, and physiology. Orthologous gene function is often unvalidated, and homology-based annotations should be interpreted with caution. To avoid mischaracterizing a cell type, we sought to use multiple orthologous genes and genes predicted to encode a protein directly related to the function of the cells. We often used gene names for annotation to avoid the pitfall of presuming *Drosophila melanogaster* cell type orthology from gene orthology.

We first annotated tissues individually, which offered a higher cell type resolution (Figures 1, 2, 6, Data S2, Table S1 and Zenodo Supplemental Data). To understand the broader cell type relationships, we integrated data across sexes and dissected tissues (Figures 1C and 1D). Cells from tissues across samples merged as expected (e.g., follicular cells identified from female abdomen and female ovary samples merged) (Figures S1D-S1F). We discerned 66 distinct cell types (“Level 2” annotations) (Figures 1F and S2C), grouped into 14 major cell type categories (“Level 1” annotations) (Figures 1E, S2B, and S2C) using marker genes such as *Syt1* (*AAEL000704*) in neurons, *repo* (*AAEL027131*) in glia, *FAS1* (*AAEL001194*) in fat tissue, *FAS2* (*AAEL008160*) in oenocytes, *titin* (*AAEL002565*) for muscle, *Ppn* (*AAEL019468*) for hemocytes, *nub* (*AAEL017445*) for enterocytes, and *Delta* (*AAEL025606*) for enteroblasts/intestinal stem cells. *grh (AAEL001168)* and *snu (AAEL018334)* were used as non-exclusive markers for epithelial-like cells, though lack functional validation in *Aedes aegypti*. Reproductive tissues and salivary glands cell types were distinct and primarily composed of cells of those specific tissue dissections, with expected contributions from abdomen and thorax samples (Figures 1D). All processed data and annotations are available via the UCSC Cell Browser^62^ (http://mosquito.cells.ucsc.edu).

### Annotation of male testes and identification of spermatids

To validate the quality of our snRNA-seq data and our annotation approach, we focused first on male testes. Mosquito testes, a potential target for mosquito control, have well-characterized cell types and marker genes. We dissected testes from 212 male mosquitoes and acquired 12,074 nuclei after quality-control filtering (Figure 2A). We identified 14 distinct cell types and confirmed these annotations with RNA fluorescence *in situ* hybridization (Figure 2A-2G).

**Figure 2.**
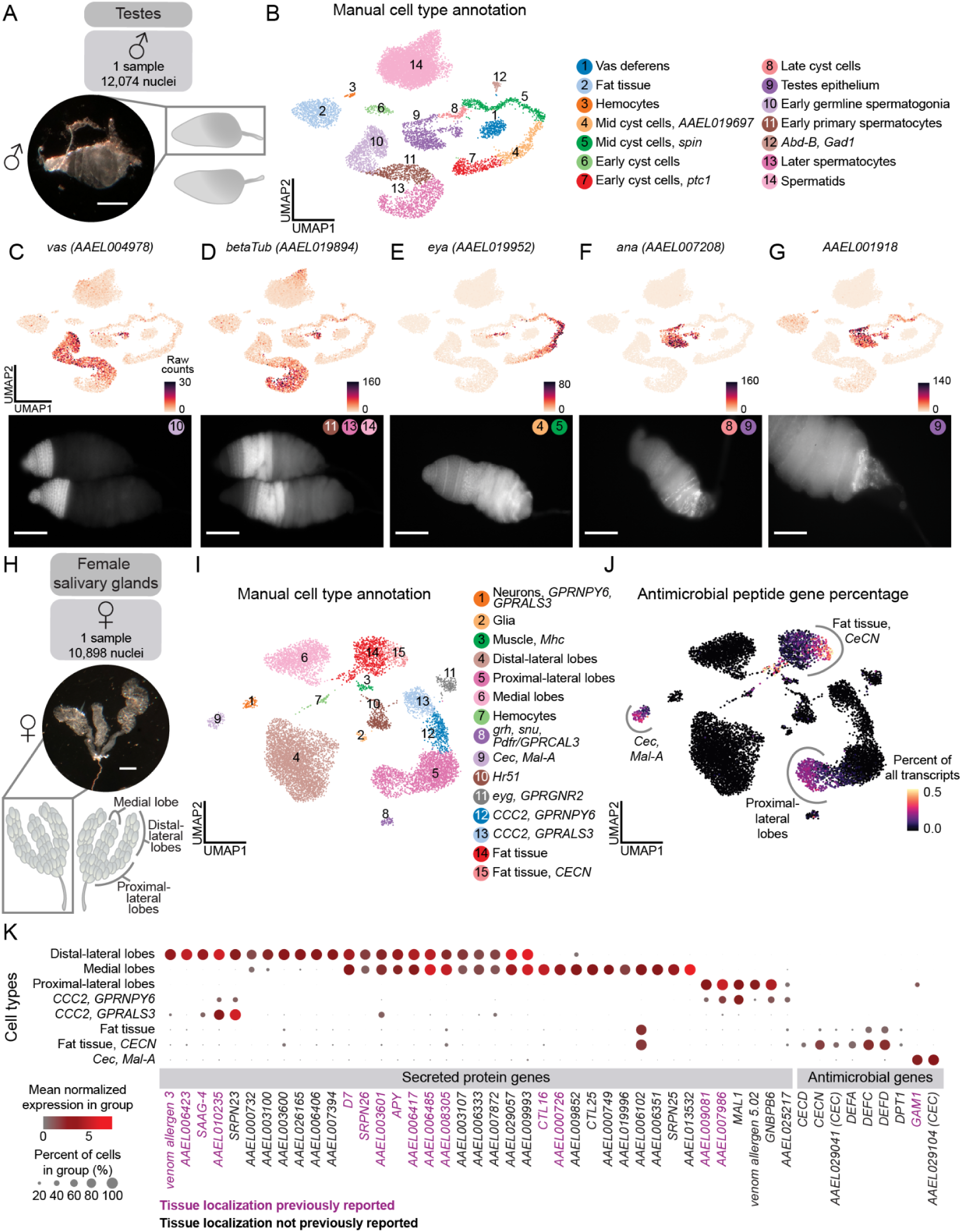
Localization and validation of male testes and female salivary gland RNA transcripts. **(A)** Dissected male testis with anatomical diagram of testes pair. One sample (10x Genomics library) yielded 12,074 nuclei from 212 animals (male, mated, sugar-fed). Scale bar: 500 μm. **(B)** UMAP of nuclei, colored and numbered by manual cell type annotation (legend). **(C-G)** UMAP illustrating raw counts (unique molecular identifiers) of a selection of genes used to annotate testes data. Corresponding validation using RNA *in situ* hybridization (below) are labeled with indicated cell types (number and color from B). Genes: *vas* (*AAEL004978*) (C), *betaTub* (*AAEL019894*) (D), *eya (AAEL019952*) (E), *ana (AAEL007208)* (F), and *AAEL001918* (G). Raw counts shown to correlate expression patterns to *in situ* images (normalized gene expression shown in Data S1). Scale bar: 100 μm for C-F; 50 μm for G. **(H)** Dissected female *Aedes aegypti* salivary gland with anatomical diagram of salivary gland pair. One sample yielded 10,898 nuclei from 495 animals (female, mated, sugar-fed). Scale bar: 500 μm. **(I)** UMAP of nuclei, colored and numbered by manual cell type annotation (legend). **(J)** Fraction of total transcripts per cell of antimicrobial peptides gene set: *CECD* (*AAEL029046*), *CECN* (*AAEL029047*), putative cecropins (*AAEL029041* and *AAEL029104*), *DEFA* (*AAEL003841*), *DEFC* (*AAEL003832*), *DEFD* (*AAEL003857*), *DPT1* (*AAEL004833*), *GAM1* (*AAEL004522*). Ends of color bar trimmed 0.1% for visibility. **(K)** Dot plot illustrating mean normalized expression of secreted protein and antimicrobial genes by cell type (Table S1). Localization of genes colored in purple has been validated by previous work^91^. Normalized expression is *ln([(raw count/total cell counts)*median total counts across cells]+1)*.

We identified the germline lineage using the expression of the widely conserved marker *vas* (*AAEL004978*)^65^, and spermatocyte-specific *betaTub* (*AAEL019894*)^66,67^ (Figure 2B-2D). Initially, spermatids were absent from our analysis, due to characteristically low transcriptional activity^18,68,69^ (Data S1), but modifying our filtering criteria revealed a distinct cluster expressing *S-Lap* (*AAEL000108*), *DBF4* (*AAEL008779*)^70^, and *Orco* (*AAEL005776*)^71^ (Data S1). We observed clusters representative of the stages of cyst cell development, with the mid-stages expressing *eya* (*AAEL019952*)^72^ (Figure 2E). *ana* (*AAEL007208*) was detected in the testes epithelium, particularly towards the posterior of the testis, and in late cyst cells (Figure 2F). *AAEL001918* was also detected in the terminal epithelium and was enriched in the most posterior region (Figure 2G). These findings suggest the existence of a transcriptomically-distinct terminal subpopulation of the testes epithelium.

### Enhanced spatial mapping of infection-related genes

Female mosquitoes inject saliva, produced by the salivary glands, beneath the skin during blood feeding. Salivary components influence the host immune response and reduce pain, allowing the mosquito to feed undetected^73–76^, and facilitate pathogen transmission^76–88^. The paired salivary glands consist of three lobes (the proximal-lateral, distal-lateral, and medial lobes) (Figure 2H), each surrounded by a basal lamina containing a single layer of saliva-secretory cells which is arranged around a central duct with an apical cavity for saliva storage^89–91^. We dissected salivary glands from 495 female mosquitoes and obtained 10,898 nuclei after quality-control filtering (Figure 2H). Using known marker genes from recent studies^77,92–99^, we annotated all expected lobes and cell types (Figure 2I and Table S1). The majority of saliva protein genes localized to the three lobes, as confirmed by published RNA *in situ* hybridization and immunofluorescence data^91,100–102^(Figure 2K). We also identified cell type-specific expression of 24 secreted proteins previously identified by mass spectrometry^92,103,104^, but whose secretory cell types were unknown (Figure 2K and Table S1).

Antimicrobial peptide genes are important for mosquito innate immunity against pathogens they transmit to humans^87^. In *Drosophila melanogaster*, fat body cells synthesize antimicrobial peptides for secretion into hemolymph^105^. We found antimicrobial peptide genes, including cecropins and defensins, in the fat tissue, as well as in enterocytes and intestinal stem cells (Figures 2J and Data S1). Transcriptomic access to cells involved in secreted salivary proteins and mosquito immunity may stimulate new avenues of investigation into viral transmission and vector effectiveness.

### *ppk317* labels a previously unknown male-specific cell type in the antenna

Female mosquitoes rely on their antennae to detect human body odor during host-seeking^7,106–110^. While male mosquitoes seek out humans to mate with females^111^, it is not known if they are attracted to the same cues as females^112^. The female antenna has been extensively investigated^36,37,113–118^, however, the male antenna is largely unexplored. To understand sex-specific cellular composition, we performed snRNA-seq on one male and two female antenna samples, integrating these with previously published female data^36^, for a total of 24,046 female nuclei and 8,016 male nuclei (Figures 3A-3B). This revealed shared and sex-specific subpopulations (Figures 3B, S3A, and S3D). We avoided batch correction to preserve biological differences^119^ and instead used marker genes to identify divergent cell types (Figures S3C-S3D).

**Figure 3.**
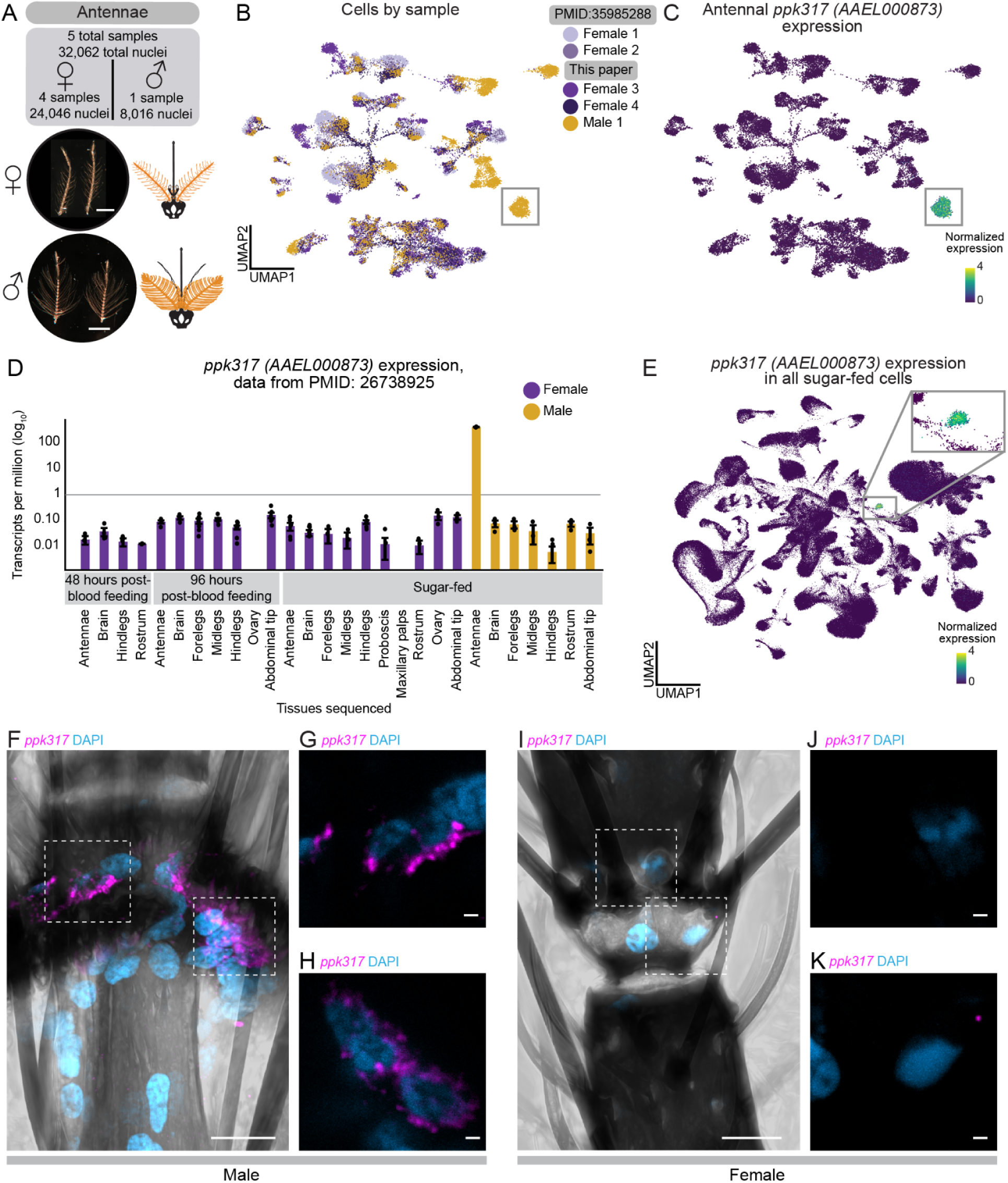
Male-specific *ppk317* cell type in the *Aedes aegypti* antenna. **(A)** Dissected antennae from female (top) and male (bottom) *Aedes aegypti* with anatomical diagrams (orange). Five samples yielded 32,062 nuclei from ∼2400 animals (mated, sugar-fed). Scale bar: 500 μm. **(B)** UMAP of antennal nuclei, colored by sample (female = 4, male = 1). Putative male-specific cluster highlighted (grey box). **(C)** *ppk317* (*AAEL000873*) normalized expression. Cluster with high expression highlighted (grey box). Normalized expression is *ln([(raw count/total cell counts)*median total counts across cells]+1)*. **(D)** *ppk317* expression [transcripts per million (log_10_)] in previously published bulk RNA-seq data of indicated tissues^26^. Female tissues (purple) are from animals that were sugar-fed, post-blood feeding (48 or 96 hours). Male tissues are sugar-fed (yellow). Both sexes were mated; females were not provided an egg-laying substrate before tissue collection. **(E)** *ppk317* normalized expression in all mated, sugar-fed nuclei. Cluster with high expression highlighted (grey box, enlarged in inset). **(F)** Maximum-intensity projection of *ppk317* RNA *in situ* hybridization (magenta) with DAPI nuclear staining (blue) from whole-mount male antenna (mated, sugar-fed). Scale bar: 10 μm. **(G-H)** Single Z plane (0.24 μm) corresponding to highlighted boxes from (F). Left box (G), right box (H). Scale bar: 1 μm. **(I)** Maximum-intensity projection of *ppk317* RNA *in situ* hybridization (magenta) with DAPI nuclear staining (blue) from whole-mount female antenna (mated, sugar-fed). Scale bar: 10 μm. **(J-K)** Maximum-intensity projection corresponding to highlighted boxes from (I). Left box (J), right box (K). Scale bar: 1 μm.

We focused on a male-specific cluster marked by *ppk317* (*AAEL000873*), a gene in the *pickpocket* (PPK) ion channel (DEG/ENaC) family^120–122^, expressed exclusively in the male antenna^26^ (Figures 3B-3E, S3D, and S3H-S3I). Male-specific *ppk317* cells are likely epithelial-related, based on their expression of *grh*, a *Drosophila melanogaster* epithelial marker^18,123^. *ppk317* cells are relatively homogenous and highly distinct relative to other antenna cell types, based on diffusion component analysis, gene-expression correlations, and partition-based graph abstraction, suggesting they represent a unique cell population (Figures S3E-S3G and S3J). There was no female counterpart to the male *ppk317* cells, which may reflect its absence in females or relationship to a more distant homologous cell type that does not express *ppk317*. RNA *in situ* hybridization confirmed selective expression in male antennal joints, with no expression in female antenna (Figures 3F-3K and Data S3). While the function of these male-specific *ppk317* cells is unknown, they may support sexually dimorphic physiology or behavior involving the male antenna.

### A precise sexual dimorphism in a single antennal chemosensory cell type

Understanding the mosquito olfactory system is crucial to deciphering how mosquitoes excel at locating human hosts. Insects detect chemosensory cues with heteromultimeric ligand-gated ion channels encoded by three large multigene families: odorant receptors (ORs), ionotropic receptors (IRs), and gustatory receptors (GRs). These receptors assemble into complexes composed of broadly expressed co-receptors and more selectively expressed ligand-specific subunits. Recent work using snRNA-seq and other methods showed that female *Aedes aegypti* olfactory sensory neurons co-express both co-receptors and ligand specific receptors within and between major receptor families^36,37^. We investigated whether these co-expression patterns also occur in the male antenna.

From the male and female antennal nuclei, we isolated 7,950 neurons (7,003 females, 947 males) (Figures 3A-3B, 4A and Data S2). We excluded neurons expressing *nompC* (*AAEL019818*), a putative mechanosensory receptor (9% of neurons), in addition to other filtering parameters (Figures S4A-S4I). We manually annotated 54 olfactory sensory neuron cell types based on unique chemoreceptor and putative transcription factor gene patterns (Figure 4B and Data S2). In at least 6 cases, chemoreceptor genes co-expressed within a cluster but not within the same cells, suggesting distinct cell types that share phenotype space but that are computationally indistinguishable without targeted analysis (e.g., *Ir41b* and *Ir41e* in Figures 4C-4D, asterisk-marked cell types in Figure S5B). These findings align with a recent study identifying ∼60 olfactory sensory neuron cell types in the female antenna^37^. We confirmed that *Aedes aegypti* olfactory sensory neurons can express multiple ligand-specific chemoreceptors, sometimes across receptor families^36,37^. Our new female samples replicated co-expression of *Orco and Ir25a* as well as the previously reported *Ir41l* neuron profile, which includes co-receptors *Orco, Ir25a, Ir76b*, and ligand-specific receptors *Ir41l, Ir41m*, *Or80, Or81,* and *Or82*^36^ (Figures 4C-4D and S4L). We also identified *ppk205* expression in *Ir41l* neurons (Figures 4C-4D).

**Figure 4.**
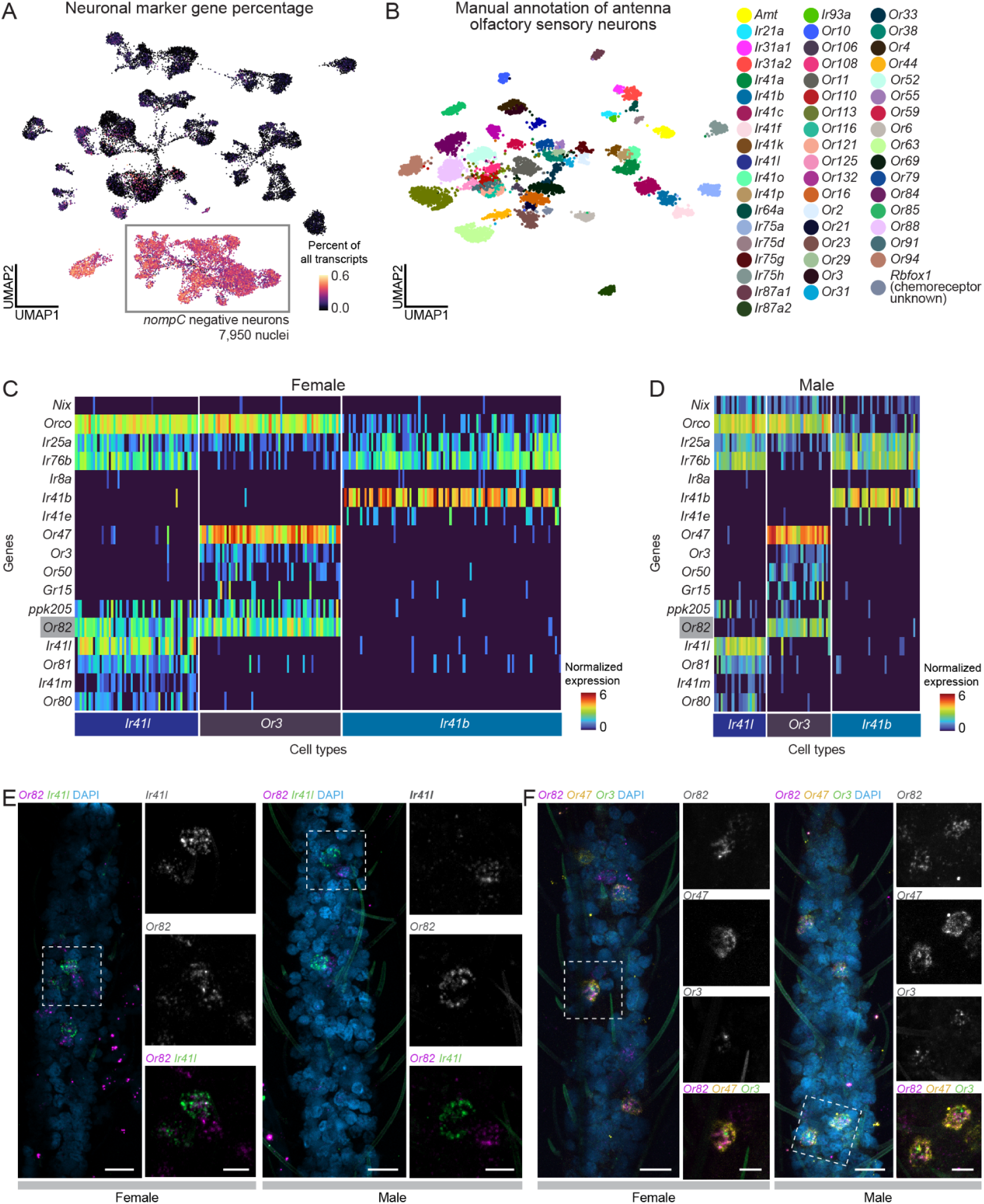
Precise sexually-dimorphic expression of *Or82* in a single antennal chemosensory cell type. **(A)** Fraction of total transcripts per cell of neuronal genes set: *Syt1* (*AAEL000704*), *brp* (*AAEL018153*), *nSyb* (*AAEL024921*), *CadN* (*AAEL000597*). *nompC* (*AAEL019818*)-negative cells highlighted (grey box). For *nompC* gene percentage, see Figure S4B. **(B)** UMAP of antenna *nompC*-negative (olfactory sensory) neurons, colored by manual cell type annotation (legend). **(C-D)** Heatmap of *Ir41l, Or3*, and *Ir41b* cells female (C) and male (D) samples. Selected genes are indicated in rows, cells in columns, with cell type annotations below. Heatmap colors represent normalized expression. Normalized expression is *ln([(raw count/total cell counts)*median total counts across cells]+1)*. **(E)** Maximum-intensity projection of *Or82* (magenta) and *Ir41l* (green) RNA *in situ* hybridization with DAPI nuclear staining (blue) from whole-mount female and male antenna (mated, sugar-fed). Scale bar: 10 μm. Highlighted white boxes enlarged to the right, with indicated probes. Scale bar: 5 μm. **(F)** Maximum-intensity projection of *Or82* (magenta), *Or47* (yellow) and *Or3* (green) RNA *in situ* hybridization with DAPI nuclear staining (blue) from whole-mount female and male antenna (mated, sugar-fed). Scale bar: 10 μm. Highlighted white boxes enlarged to the right, with indicated probes. Scale bar: 5 μm.

We investigated the sex differences in mosquito olfactory sensory neurons. Despite well-known sexually dimorphic olfactory behaviors^12,112,124,125^, transcriptional differences between male and female olfactory sensory neurons were limited. All annotated cell types contained both male and female cells, although in varying proportions (Figures S4I-S4J). Differential gene expression^55^ identified frequent sex-specific expression of the ADP/ATP carrier protein *SLC25A5* (*AAEL004855*), a putative Mg^2+^/Na^+^ transporter (*AAEL009150),* the male-determining factor *Nix* (*AAEL022912*), a putative serine/threonine kinase (*AAEL004217*), and the odorant binding protein *OBP35* (*AAEL002606*) (Figure S4M and Table S2). Of these, only *Nix* has a known role in sex determination. Out of 403 putative sensory genes, including ORs, IRs, GRs, PPKs, transient receptor potential (TRP) ion channels, opsins, and mechanosensory receptors (Table S1), only four (*Or82*, *Ir25a*, *Ir76b*, and *Or2*) showed significant sex-specific expression differences (Figures S4N-S4Q and Table S2). To rule out artifacts, we examined raw counts (unique molecular identifiers) and confirmed that transcript abundance of sensory genes was consistent across samples and sexes (Figure S5C).

We found an exception to the high degree of similarity between male and female chemoreceptor expression in the *Ir41l* neurons: only females express *Or82*, despite otherwise identical chemoreceptor profiles (Figures 4C-4D and S4N). RNA *in situ* hybridization confirmed *Or82* co-localization with *Ir41l* in females but not males, while in both sexes *Or82* is present in *Or3* and *Or47* neurons (Figures 4C-4F). The ligand profile of *Or82* is unknown and it is unclear if or how its absence in male *Ir41l* neurons contributes to sex-specific behaviors. These findings highlight the broad similarity of chemoreceptor expression between male and female olfactory sensory neurons, with one notable exception.

### Molecular signature of polymodal sensory detection in leg sensory neurons

*Aedes aegypti* legs contribute to host seeking^10^, blood feeding^126^, mating^127–129^, and oviposition^122,130^ through detection of a wide array of stimuli including temperature^10^, osmolality^122^, bitter compounds^126,131,132^, sugars^133^, pheromones, and amino acids^134^. Mosquitoes have three pairs of legs (forelegs, midlegs, and hindlegs) with neuronal cell bodies concentrated in the most distal segment, the tarsi^10,135,136^. From 332 female and 298 and male mosquitoes, we obtained 29,323 tarsal nuclei, including a population of 1,060 sensory neurons (*nompC*-negative) (Figures 5A-B and Data S2). Clustering revealed distinct chemosensory receptor profiles (Figures 5C-5D and Zenodo Supplemental Data).

**Figure 5.**
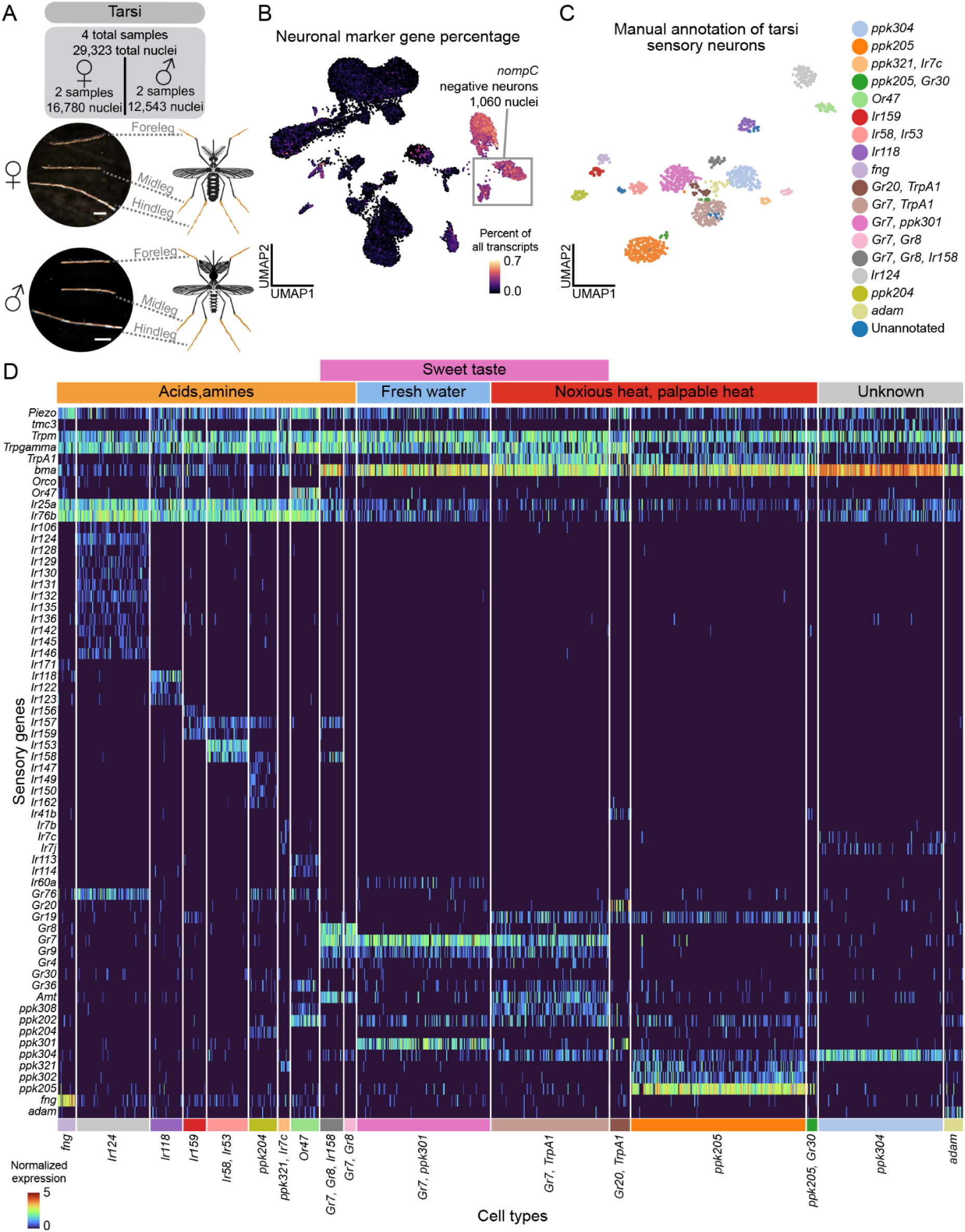
Tarsi sensory neurons are polymodal. **(A)** Dissected tarsi from female (top) and male (bottom) *Aedes aegypti* with anatomical diagram (orange). Four samples yielded 29,323 nuclei from 630 animals (mated, sugar-fed). Scale bar: 500 μm **(B)** Fraction of total transcripts per cell of neuronal genes set: *Syt1*, *brp*, *nSyb*, *CadN*. *nompC*-negative cells highlighted (grey box). For *nompC* gene percentage, see Data S4. **(C)** UMAP of tarsi chemosensory (*nompC*-negative) neurons after filtering, colored by manual cell type annotation (legend). **(D)** Heatmap of chemoreceptor gene expression in all annotated clusters. Selected genes indicated in rows, cells in columns, with annotations for cell type (below) and respective sensory function (above). Heatmap colors represent normalized expression. Normalized expression is *ln([(raw count/total cell counts)*median total counts across cells]+1)*.

Some tarsal sensory neurons co-express multiple receptor families (e.g., IRs, GRs, and PPKs). Co-expression of IRs and PPKs has also been observed in *Drosophila melanogaster* tarsi^137^. For example, *ppk204* neurons co-express IR co-receptors along with ligand-specific receptors and GRs (Figures 5C-5D). *Or47* neurons co-express PPKs in the antenna and tarsi, and with IRs in the proboscis and the tarsi (Figures 4C-4D, 5C-5D, S5, S6, Data S2 and Zenodo Supplemental Data). Intriguingly, only tarsal *Or47* neurons lack the obligate OR co-receptor *Orco*, raising questions about receptor function in this context. These data show that mosquito subsets of neurons in the tarsi, proboscis, and antenna co-express chemosensory receptors from multiple gene families.

Subpopulations of tarsal neurons also co-express receptors known to operate in distinct sensory modalities including taste, heat, and osmolality, suggesting that these neurons are polymodal. For example, tarsal *ppk301* neurons, critical for freshwater detection during oviposition^122^, co-express sweet taste receptors *Gr7* and *Gr9* (Figure 5D). In the proboscis, these genes are expressed in separate cell types (Figure S6C), suggesting appendage-specific receptor combinations for different sensory coding functions.

Additionally, a subset of *Gr7* neurons co-express with *TrpA1* (*AAEL001268*), which functions in noxious heat detection^138^, indicating these neurons may detect both sweet taste and heat. Co-expressed *ppk304* and *ppk102* are putative orthologues of *Drosophila melanogaster ppk29* and *ppk23*, subunits of a pheromone-sensing channel^139^ (Figure 5D). We detected sparser chemoreceptor expression in other tissues, however note expression of *Gr39* in the wings and *Gr20*/*Gr60* in the abdominal tip (Zenodo Supplemental Data). These polymodal sensory neurons may enable complex behavioral responses, though many of these receptors still require functional validation.

We investigated sensory neurons from the tarsi, proboscis, and maxillary palp for sexual dimorphism (Figures S7A-S7B and Data S4). Proboscis *Ir7e* neurons were female-specific and tarsal *ppk205/Gr30* neurons were male-specific, though this was based on small cell numbers (19 and 12 cells, respectively; Data S4). Given the complexity of chemosensory neuron populations, data from more nuclei will likely improve resolution of rarer cell types. Other cell types varied in abundance between sexes but were not sex specific. No sensory-related genes were differentially expressed across clusters (Data S4 and Table 2), underscoring the strong molecular similarity between male and female sensory neurons.

### Sensory neurons express a cell type-specific neuropeptide receptor code

Neuropeptides modulate mosquito behavior and physiology. In *Aedes aegypti*, over 100 predicted neuropeptides regulate diverse processes, including host seeking, blood feeding, and reproduction^17,140,141^. To examine their role in sensory neurons, we analyzed the expression of 122 neuropeptide genes. Some receptors were broadly expressed (e.g., *SIFaR1*, *InR*, *GPRNPY7*, *NPYLR3*), while other receptors showed cell type-enriched patterns aligned with chemosensory receptor profiles (Figure S7 and Data S4). This receptor code suggests that neuropeptides may modulate sensory neurons in a cell type-specific manner.

### Sexually dimorphic Kenyon cells and glia in the brain

The central nervous system coordinates how sex^142,143^ and blood-feeding states^13–15,17,52,144^ modulate mosquito behavior^142,143^. We collected 68,898 brain nuclei (21,820 and 16,349 nuclei from mated, sugar-fed female and males, and 30,729 from mated, blood-fed females) (Figures 6A-6B, and S8A) and 17,610 thoracic ganglia of the ventral nerve cord (9,306 female and 8,304 male nuclei) (Data S2).

**Figure 6.**
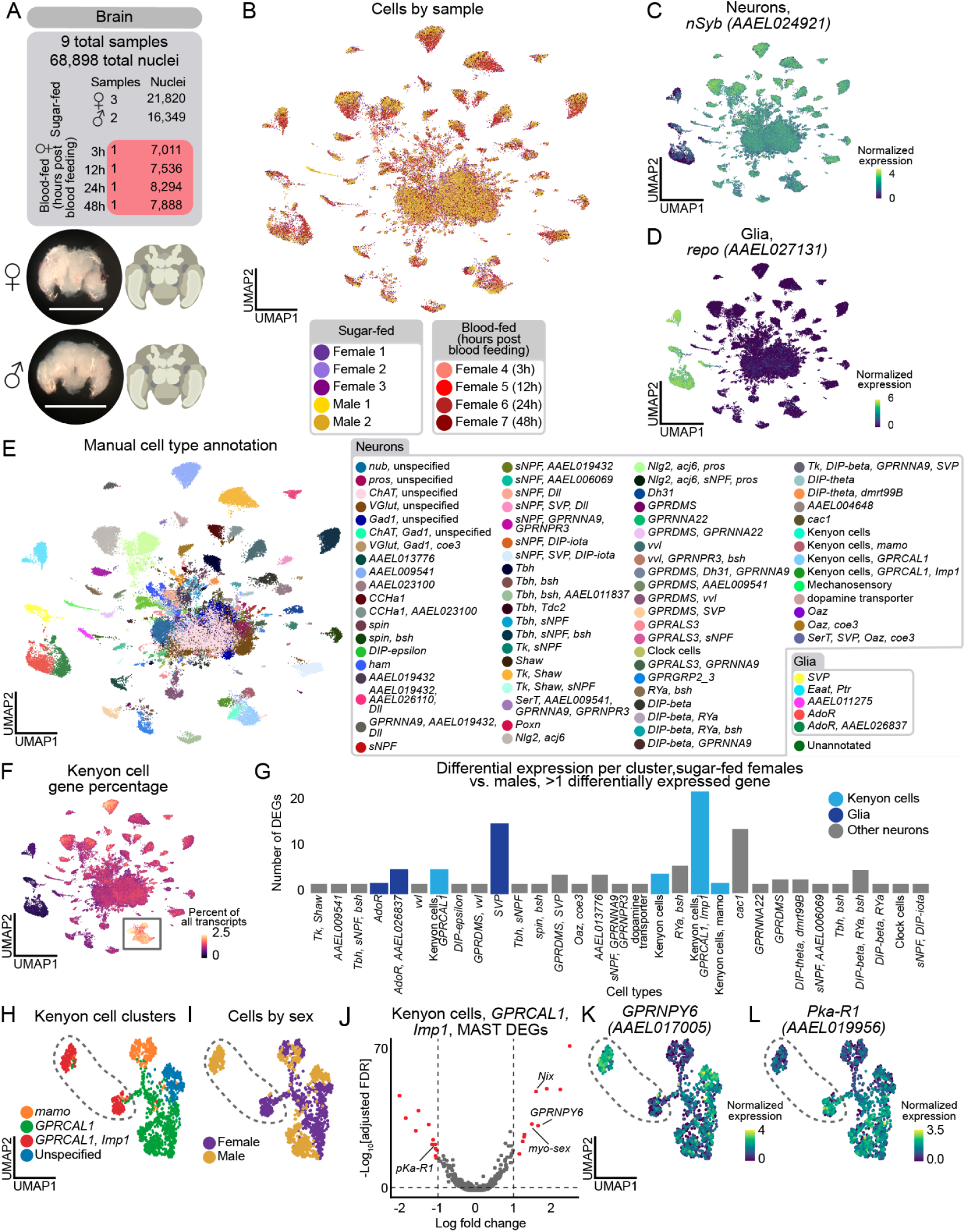
Brain annotation identifies sexually dimorphic Kenyon cells and glia. **(A)** Dissected brain from female (top) and male (bottom) *Aedes aegypti* with anatomical diagram. Data was collected from sugar-fed females and males, and blood-fed females 3, 12, 24, and 48 hours after blood feeding. Five samples yielded 68,898 nuclei from 182 animals. Scale bar: 500 μm **(B)** UMAP of brain nuclei, colored by sample (female = 7, male = 2). **(C-D)** Normalized expression of neuronal marker *nSyb* (C) and glial marker *repo (AAEL027131)* (D). Normalized expression is *ln([(raw count/total cell counts)*median total counts across cells]+1)*. **(E)** UMAP of nuclei from all samples, colored and numbered by manual annotation using marker genes (legend). Major cell type annotations represented in shaded gray headers. **(F)** Fraction of total transcripts per cell of 30 putative Kenyon cell gene markers (Table S1). Annotated Kenyon cells highlighted (grey box). **(G)** Bar plot showing cell types with at least 2 differentially expressed genes (DEGs) between sugar-fed male and female cells. Clusters colored by cell identity: Kenyon cells (light blue), glia (dark blue), other neurons (grey) (Table S3). Significant genes had |log fold change| >1, false discovery rate <0.05, determined by MAST on normalized expression. **(H-I)** UMAP of Kenyon cell nuclei from all sugar-fed brains, colored by manual cell type annotation (H), and by sex (I). *GPRCAL1, Imp1* cells *(AAEL010043, AAEL006876)* highlighted (dotted area). **(J)** Volcano plot of DEGs in *GPRCAL1, Imp1* Kenyon cells. Significant genes indicated in red. Male biased genes on right, female biased genes on left. **(K-L)** *GPRNPY6 (AAEL017005)* (K) and *pKa-R1 (AAEL019956)* (L) normalized expression in Kenyon cell nuclei from all sugar-fed brains.

In our brain data, 92% of nuclei were neurons and 8% were glia (Figures 6C-6D), consistent with estimates that the *Aedes aegypti* brain contains ∼220,000 neurons (out of ∼250,000 total cells)^145^. To assess our sampling depth of neuronal cell types, we looked for the central clock cells, a group of fewer than 15 cells in the adult mosquito brain^146,147^. We identified a small cluster marked exclusively by *Pdf* (*AAEL001754*) and other circadian genes (Figures S9C-S9D), demonstrating our ability to identify rare cell types. We manually annotated cell types based on marker gene expression (Figures 6E, S8B, and S9A), in most cases relying on ortholog information from *Drosophila melanogaster* for neuron and glia subtypes that will require further validation. Alignment of our mosquito brain and *Drosophila melanogaster* head data using SAMap^148^ also supported our annotations (Figure S9 and Table S3). This included the identification of Kenyon cells in the mushroom body, a conserved invertebrate brain structure involved in learning and memory^149^, based on high alignment scores and of orthologs of *Drosophila melanogaster* Kenyon cell gene markers (Figures 6F, S9J, S9L-S9P, and Table S3)^18,150^. Due to considerable evolutionary distance and similarities between neuron types within species, we cautiously used mapping scores and orthologous genes to infer cell type identity.

Using our annotated brain data, we investigated sex-specific gene expression differences within cell types^55^ (Figure 6G and Table S3). We tested cell types with >10 cells in the male and non-blood-fed female conditions. 28 of 72 neuronal and 3 of 5 glial cell types had at least 2 differentially expressed genes. Among the frequently differentially expressed genes were four involved in sex determination and sex-specific neuronal function: *Nix* and *myo-sex (AAEL021838)* were upregulated in males, while *fru* (*AAEL024283*) and *dsx* (*AAEL009114*) were upregulated in females (Table S3). *nompC* mechanosensory neurons were exclusive to male samples (Figures 6E, S8A, and S9B). Otherwise, cell type abundance was similar across sexes (Figure S8A).

Kenyon cells expressing *GPRCAL1* (*AAEL010043*) and *Imp1* (*AAEL006876*) (Figures 6H-6I) showed the strongest sex-specific gene expression (Figure 6G). Neuropeptide Y receptor *GPRNPY6* (*AAEL017005*) was upregulated in males while signaling receptor *Pka-R1* (*AAEL019956*) was upregulated in females (Figures 6J-6L). *SVP* (*AAEL002765*) glial cells also had pronounced sex-specific gene expression (Figure 6G), aligned with recent work from *Drosophila* species suggesting glia may be sites for brain adaptation^151^.

### Glial cells display dramatic transcriptional changes in the female brain after blood feeding

Blood feeding induces a sequence of dramatic physiological and behavioral changes in the female mosquito. To understand how brain cell types may contribute to these, we collected snRNA-seq data from the brain at 3, 12, 24 and 48 hours after blood feeding (Figure 7A). For each cell type, we compared gene expression at each timepoint to the corresponding sugar-fed cells^55^. (Figure 7B and Table S2). Cell type abundance remained stable across timepoints (Figure S8A).

**Figure 7.**
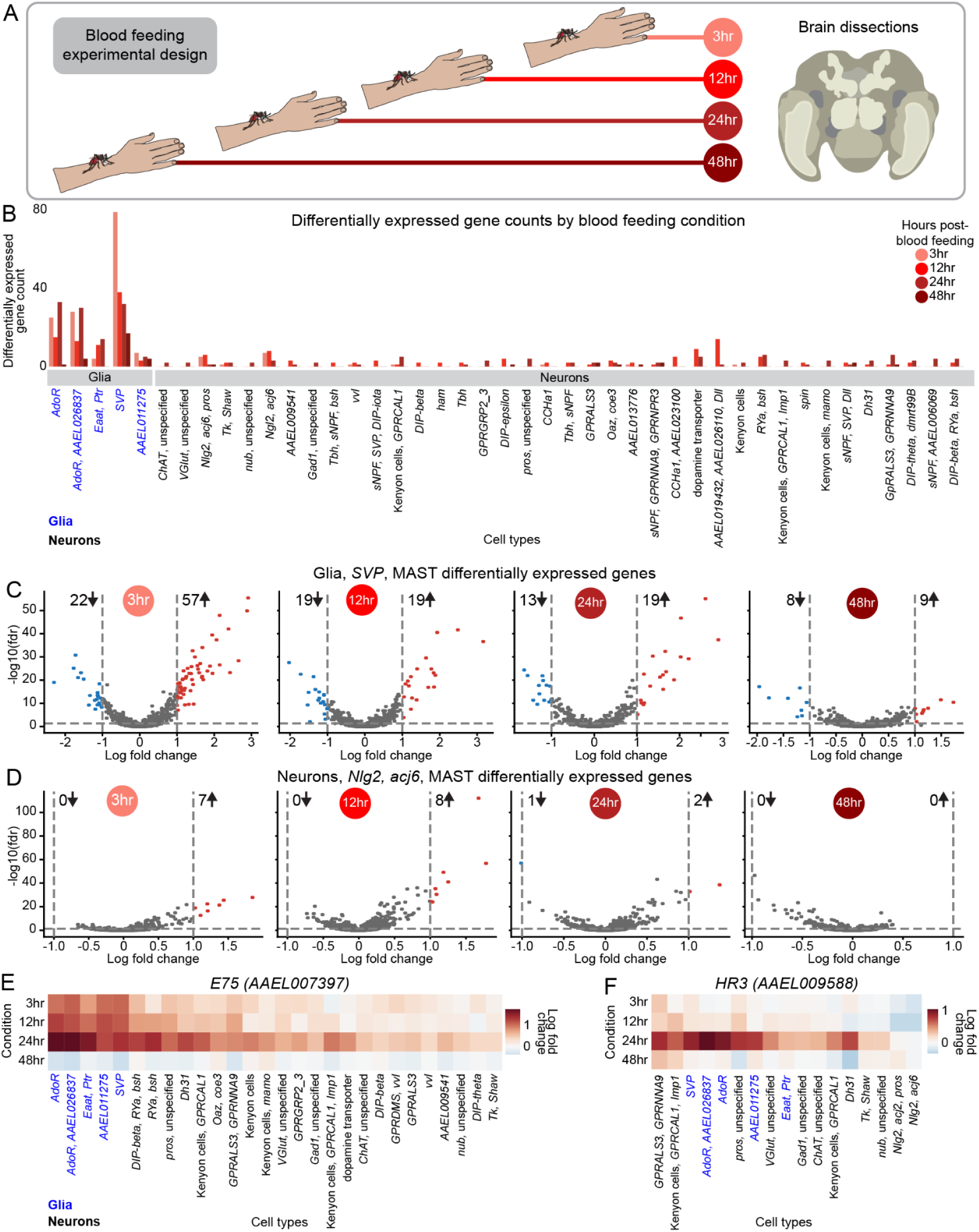
Glia show extensive transcriptional changes after blood feeding. **(A)** Blood feeding experimental design. **(B)** Bar plot showing cell types with differentially expressed genes (DEGs) between sugar-fed and blood-fed female cells. Bars colored by blood-feeding condition. Glia and neuron cell types labeled below. Significant genes had |log fold change| >1, false discovery rate <0.05, determined by MAST on normalized expression (Table S2). Normalized expression is *ln([(raw count/total cell counts)*median total counts across cells]+1)*. **(C-D)** Volcano plots of DEGs for *SVP* glia *(AAEL002765*) (C) and *Nlg2, acj6* neurons *(AAEL014303, AAEL005507)* (D) for sugar-fed compared to indicated blood-feeding timepoint Number of downregulated (blue) and upregulated (red) genes indicated alongside arrows. **(E-F)** Heatmaps of log fold change of *E75 (AAEL007397)* (E) and *HR3 (AAEL009588)* (F) for glia (blue) and neuron (black) cell types. Cell types are sorted by the total log fold change across all timepoints. Cell types shown have >10 cells in each timepoint, and at least one timepoint where change from sugar-fed condition had a false discovery rate <0.05.

Contrary to expectations, glial cells showed more dramatic transcriptomic changes than neurons after blood feeding (Figure 7B). *SVP* glia demonstrated the strongest shift, peaking 3 hours post-blood feeding with 79 differentially genes, followed by 38 at 12 hours, 32 at 24 hours, and 17 at 48 hours. Expression was largely upregulated at 3 hours (72% of differentially expressed genes) but later time points exhibited a more balanced mix of up- and down-regulated genes (Figure 7C and Table S2). Neuronal responses to blood feeding were more modest, though 38 of 47 neuronal cell types (>10 cells per timepoint) expressed at least two differentially expressed genes. Neuron cell types marked by “*Nlg2*, *acj6*, *pros*”, “*Ngl2*, *acj6*” (Figure 7D), “*AAEL019432, AAEL026110, Dll”*, dopamine transporter *(AAEL024732)*, “*RYa, bsh”* showed the greatest changes (Figure 7B and Table S2).

Next, we investigated the expression dynamics of individual genes. *E75* (*AAEL007397*), *EcR* (*AAEL019431*), and *HR3* (*AAEL009588*) are nuclear steroid hormone receptors involved in ecdysone signaling, which regulates multiple processes in insects, including blood-feeding-induced changes in female mosquito reproduction^152^. *E75, EcR,* and *HR3* are widely expressed in both glia and neurons in our sugar-fed brain data (Figures S10A-S10D). 3 to 24 hours post-blood feeding, *E75* and *EcR* were strongly upregulated in all glial and several neuronal cell types, peaking at 24 hours and declining at 48 hours (Figures 7E and S10E). Their greater upregulation in glia suggests that glia may be the primary mediators of their role in the blood-feeding response. In contrast, *HR3* showed relatively little change from sugar-fed levels at early timepoints but increased sharply at 24 hours in a subset of both glial and neuron cell types, indicating a distinct temporal expression pattern (Figure 7F). Other genes demonstrated different dynamics. For instance, insulin-like peptide *IA-2* (*AAEL005692*) was upregulated in a small subset of neurons at 12 hours and 24 hours post-blood feeding and then downregulated in a broader range of glial and neuron cell types at 48 hours (Figure S10F). The clock genes *ITP* (*AAEL019725*) and *PER* (*AAEL008141*) were significantly upregulated in glia but not neurons, although with unique temporal and cell type expression patterns (Figures S10I-S10J).

Sexual dimorphism-related transcription factors *fru* and *dsx* also showed changes after blood feeding. *dsx* was downregulated at 3, 12, and 24 hour post-blood feeding, almost exclusively in glia, before returning to near-baseline by 48 hours (Figure S10G). Conversely, *fru* showed modest changes, although was notably downregulated in *SVP* glia at 48 hours (Figure S10H). Cell type-specific *fru* regulation has also been observed in *Drosophila melanogaster*, where the male isoform masculinizes brain circuitry through uniquely regulating effector genes in different neuronal cell types^153–156^. Whether *fru* could play a similar role in the behavioral states of the female mosquito is unknown. These findings highlight potential regulators of gene expression changes both globally and in specific cell types across blood-feeding timepoints in the brain.

Our data confirm that gene expression changes in the female mosquito brain are correlated with blood-feeding state^26^. While some changes occur in neurons, more pronounced transcriptomic changes occur in glia. The functional implications of these glial response patterns for mosquito metabolism, physiology, and behavior remain to be explored.

## Discussion

### A cell atlas of the adult male and female *Aedes aegypti* mosquito

We present the first comprehensive snRNA-seq atlas of adult male and female *Aedes aegypti* profiling 367,096 nuclei from 19 tissues. This atlas provides cell type markers, selected through computational analysis and gene orthology, to annotate individual tissues sampled from the entire mosquito. This enabled insights into mosquito cellular diversity and sexually-dimorphic gene expression. All data and annotations are available through the UCSC Cell Browser (http://mosquito.cells.ucsc.edu)^62^.

The *Aedes aegypti* Mosquito Cell Atlas will aid the identification of specialized cell types and their molecular signatures as potential targets for vector control. Cell type markers throughout spermatogenesis may provide more effective targets for mosquito population control that leverage male sterility or gene drive approaches^157,158^. Increased spatial-transcriptomic mapping in the salivary gland could inform transgenic expression of antiviral effector molecules. Identification of specialized, antimicrobial peptide-expressing fat tissue cells provides an opportunity to influence mosquito immunity and vector competence. Beyond translational potential, this atlas may facilitate the development of molecular tools, including cell type-specific drivers.

### Sexual dimorphic organization of receptors in the antenna

Our analysis revealed unexpected examples of sexual dimorphism in the *Aedes aegypti* antenna, including previously unknown male-specific *ppk317* epithelial cells in antennal joints. Of the *Aedes aegypti* PPK gene family, only *ppk301* has been functionally characterized^122^. However, studies in *Drosophila melanogaster* show that PPKs have diverse functions, such as larval liquid clearance (*ppk4* and *ppk11*)^120,159^, and male courtship and pheromone detection (*ppk23*, *ppk25*, and *ppk29*)^139,160–162^. Notably, *ppk1, rpk*, and *ppk26*^122^ are closest *Drosophila melanogaster* orthologues to *Aedes aegypti ppk317*, which are involved in mechanical nociception in multi-dendritic neurons^121,163^. Future work is needed to understand whether this male-specific cell type plays a specific role in the antenna.

In antennal chemosensory neurons, overall sexual dimorphism was limited. However, we identified a striking exception in *Ir41l* cells: *Or82* is expressed in female cells but absent in males. Both male and female *Ir41l* cells express *Or3*, highlighting the precision of *Or82* transcriptional regulation. This suggests active mechanisms governing OR gene expression and sexually dimorphic sensory processing, potentially supporting an efficient evolutionary strategy for the precise tuning of sensory responses across sexes while maintaining shared olfactory functions. Investigating *Or82* regulation may uncover broader mechanisms underlying sexual dimorphism in sensory systems. Although *Or82*’s ligand profile is unknown, its female-specific expression in *Ir41l* neurons raises the possibility of a role in female sensory behaviors.

### Widespread receptor co-expression in *Aedes aegypti* sensory appendages

Mosquito sensory neurons challenge canonical principles of chemosensory organization through extensive receptor co-expression. Our data extend recent findings of co-receptor and ligand-specific receptor co-expression in antennal and maxillary palp neurons^36,37^ to other sensory appendages, including the proboscis and tarsi, suggesting a fundamental organizational principle across mosquito sensory systems. We observed two patterns: (1) co-expression of multiple ligand-specific receptors from the same family, (2) co-expression of receptors across families (ORs, IRs, GRs, PPKs, TRPs). Notably, *Or82/Ir41l* and *Or82/Or47/Or3* co-expression, which we validated with RNA *in situ* hybridization, illustrate these patterns. These data suggest coordinated receptor co-expression across gene families.

This complex organization may represent an evolutionary adaptation for efficient processing of environmental cues. While co-expression in antennae and maxillary palps has been hypothesized to enhance host detection^36^, its presence in proboscis and tarsi suggests a broader strategy. By co-expressing different receptor families, mosquito sensory neurons can detect diverse chemical cues simultaneously, enabling either specificity of behavioral responses in different contexts, or redundancy and increased signal reliability. Polymodal sensory neurons may be especially advantageous for *Aedes aegypti* as a human specialist, supporting robust host detection despite variable human odor profiles and shifting environments. Critical questions remain: Do co-expressed ligand-specific or cross-family receptors assemble into functional complexes? How do they interact? How is this information integrated by higher-order neurons? Elucidating these mechanisms could uncover fundamental principles in sensory perception through the integration of receptor multiplexing.

Beyond chemoreceptor distribution, we discovered coordinated, cell type-specific expression of neuropeptide receptors across sensory neurons. While some receptors are broadly expressed, others display restricted patterns aligned with the chemoreceptor expression profiles. This organization may allow sensory modulation based on internal state, possibly related to host seeking, post-blood-feeding behavior, or oviposition. Future work should explore how neuropeptide signaling shapes sensory neuron function and whether specific receptor combinations enable flexible tuning of sensory processing based on physiological states.

### Sexual dimorphism and glial plasticity in the mosquito brain

Our analysis identifies new cell types for the study of sexual dimorphism in the mosquitoes. Kenyon cells, associated with learning and memory^149^, show pronounced sex-specific gene expression, in particular *GPRCAL1, Imp1* cells. These include male-enriched neuropeptide Y receptor *GPRNPY6* and female-enriched protein kinase A receptor *Pka-R1* expression, suggesting sex-specific circuit modulation. Neuroanatomical evidence supports this sexual dimorphism, with some male Kenyon cells larger in size despite overall smaller male brains^164^. Given the mushroom body’s role in innate behaviors and internal states^165^, Kenyon cells may contribute to sex-specific mosquito behaviors such as host seeking or male courtship.

Glial cells emerge as key cell types in both sexual dimorphism and blood-feeding response. Glia show more pronounced transcriptomic divergence than neurons between the male and female *Aedes aegypti* brain, echoing recent cross-species comparisons in drosophilids^151^. This suggests glia transcriptional plasticity may provide a permissive substrate for insect evolutionary and sexually dimorphic plasticity, enabling novel properties without disrupting the more conserved functions of neuronal circuits.

We also find that glia, more than neurons, undergo extensive transcriptional changes following blood feeding. This highlights a broader role for glia than previously recognized^166,167^, as potential master regulators of sexual dimorphism and behavior state responsiveness. Several factors may explain glia’s extensive response.

Glia regulate blood-brain barrier permeability and are ideally positioned to detect blood- or food-derived signals and trigger immune responses^168^. Perineurial glia of the blood-brain barrier demonstrate transcriptomic divergence between *Drosophila sechellia* and *Drosophila melanogaster*, possibly reflecting dietary differences in carbohydrate intake^151^. Glia also serve a critical role in neuronal metabolic support^169^ and the extensive metabolic demands of blood meal processing^28,170,171^ and may be involved in changing sugar uptake in the brain^151^. Glial transcriptional changes could relate to the divergent molecular or metabolic responses required as a female mosquito’s diet transitions from sugar to blood.

Glia can also release neuroactive molecules and regulate the extracellular environment to broadly influence neural circuit function^172,173^. The temporal dynamics of glial gene expression, particularly in nuclear steroid hormone receptors like *HR3* and *E75*, suggest a transcriptional cascade that could maintain prolonged suppression of host-seeking behavior after blood feeding.

Understanding how specific glial populations influence neuronal function and behavior through these pathways could reveal novel aspects of glia-neuron interactions and their role in regulating mosquito behavior.

### Limitations of the Study

While our cell atlas provides insights into mosquito cellular diversity, several limitations should be considered. Although snRNA-seq enables unified profiling of tissues, nuclear transcriptomes may not fully reflect cytoplasmic mRNA levels and provide no insight on protein expression^174^. This is particularly relevant for chemoreceptor co-expression studies, where post-transcriptional regulation could affect final receptor composition^175^. Some detected transcripts could be untranslated, as seen for ORs in the clonal raider ant *Ooceraea biroi*^176^.

All cell dissociation protocols may introduce cell type bias. While snRNA-seq does not efficiently capture immune cell types in mammals^177,178^, it is less biased for attached cell types compared to single-cell methods^47^. Our nuclei extraction protocol has previously been demonstrated to accurately reflect histological cell compositions in *Drosophila melanogaster*^48^, but sampling bias may still be present.

Annotation of the *Aedes aegypti* genome is imperfect. Overlapping gene annotations can cause multimapping of transcripts during data alignment leading to transcripts being discarded, as we observed with *Or111* and *AAEL019786*. As a result, absence or low expression of genes should be interpreted cautiously.

Our cell type annotations rely heavily on *Drosophila melanogaster* orthology despite 260 million years of evolutionary separation^59,60^, potentially causing us to miss mosquito-specific adaptations. While we profiled 367,096 nuclei across the mosquito, rare cell types may remain undetected. We characterized 19 tissues, however most were not discussed here in detail, leaving opportunities for future exploration.

While we observed extensive receptor co-expression in sensory appendages, validation was limited to a few specific cell types. Whether these receptors form functional complexes or contribute to behavior requires electrophysiological, genetic, proteomic and behavioral studies. Similarly, causally linking sexually dimorphic transcript expression patterns in the antenna and brain to behavioral dimorphism will require further study.

## Resource availability

### Lead contact

Requests for resources and reagents should be directed to the lead contact, Nadav Shai.

### Materials availability

This study did not generate new unique reagents.

### Data and code availability

Supplementary Figures S1-S10, Data S1-S5 and Tables S1-S3 accompany the paper. Processed data are available for user-friendly visualization and download through UCSC Cell Browser (https://mosquito.cells.ucsc.edu). Raw snRNA-seq data are available from NCBI (BioProject: PRJNA1223381). Raw snRNA-seq data from female antenna and maxillary palp samples previously published^36^ and re-analyzed in this study are also on NCBI (BioProject: PRJNA794050). Additional processed data, plots, analysis and custom scripts are available at Zenodo Supplemental Data (https://doi.org/10.5281/zenodo.14890012).

Any additional information required to reanalyze the data reported in this paper is available from the lead contact upon request.

### Consortia

The members of the *Aedes aegypti* Mosquito Cell Atlas Consortium are (in alphabetical order): Joshua X. D. Ang, Igor Antoshechkin, Yu Cai, Fangying Chen, Yen-Chung Chen, Julien Devilliers, Linhan Dong, Roberto Feuda, Paolo Gabrieli, Artyom Kopp, Hyeogsun Kwon, Hsing-Han Li, Tzu-Chiao Lu, Thalita Lucio, João T. Marques, Marcus F. Oliveira, Roenick P. Olmo, Umberto Palatini, Zeaan M. Pithawala, Julien Pompon, Yan Reis, João Rodrigues, and Ryan C. Smith. See Data S5 for consortium affiliations and funding support.

## Acknowledgments

We acknowledge the teams, institutions and grants that support open-access databases, including VectorBase^58^ and FlyBase^180^, which were essential to this work. We thank members of the Vosshall Lab for comments on the manuscript; Libby Mejia and Melissa Dallesandro for expert mosquito rearing; Colin Berry for providing mosquito larvae for the testes RNA *in situ* hybridization experiment; Connie Zhao, Helen Duan, and Bin Zhang at the Rockefeller Genomics Core; Alison North, Priyam Banerjee, Maria Belen Harreguy Alfonso, Christina Pyrgaki, and Tao Tong at the Rockefeller Bio-Imaging Resource Center (RRID: SCR_017791) for their support and assistance with imaging; Jason Banfelder, Balakanagaram Jayaraman, and Rebecca Bennett at the Rockefeller University High Performance Computing for their expert technical support; Bushra Bibi and Vivian Niewiadonski at Novogene for operational help with sequencing; Begüm Aydin, Pyonghwa Kim, Andras Sziraki, and Andrea Terceros for helpful technical discussions; Cori Bargmann, Dana Pe’er, Vanessa Ruta, and Sarah Teichmann for insightful guidance and feedback.

This work was supported by National Science Foundation Graduate Research Fellowship (No. 1946429), Kavli Neural Systems Institute Graduate Fellowship, and Schmidt Science Fellowship (O.V.G.), B.W. and M.H. are supported by NHGRI U24HG002371 and NIMH RF1MH132662, L.H.-Z. is a Junior Fellow of the Simons Society of Fellows and was supported by EMBO Long-Term Fellowship (ALTF 1103-2019). J.R was supported by the Boehringer Ingelheim Fonds PhD Fellowship; B.R.H. by the Human Frontier Science Program Organization LT000123/2020-L. O.S.A. is supported by NIH awards R01AI151004, RO1AI148300, RO1AI175152, and EPA STAR award RD84020401. This publication was developed under Assistance Agreement No. RD84020401 awarded by the U.S. Environmental Protection Agency to O.S.A., It has not been formally reviewed by EPA. The views expressed in this document are solely those of the authors and do not necessarily reflect those of the Agency. EPA does not endorse any products or commercial services mentioned in this publication. L.B.D. is supported by R35 GM137888 NIH-NIGMS, Beckman Young Investigator Award, Pew Biomedical Scholar Award, and Klingenstein-Simons Fellowship Award in Neuroscience. H.W.-C. is supported by BBSRC BB/L004445/1. T.R.S. is supported by DP2AI177891 and is a Freeman Hrabowski Scholar of the Howard Hughes Medical Institute. R.S. is employed by SAIL, MSKCC and is supported in part by The Alan and Sandra Gerry Metastasis and Tumor Ecosystems Center (GMTEC). H.L. is a CPRIT Scholar in Cancer Research (RR200063) and supported by NIH (U01AG086143), the Longevity Impetus Grant, the Welch Foundation, the Ted Nash Long Life Foundation and the Hevolution/AFAR Foundation. This research was supported by the Stavros Niarchos Foundation (SNF) as part of its grant to the SNF Institute for Global Infectious Disease Research at The Rockefeller University. L.B.V. is supported by the Howard Hughes Medical Institute. N.S. was supported by an EMBO Long-Term Fellowship (ALTF 286-2019).

## Author contributions

O.V.G., H.L., L.B.V. and N.S. together conceived the study.

O.V.G., A.E.D., L.H.-Z., P.L., T.M., J.R., A.R.-V., T.R.S., Y.N.T., M.M.W., and N.S. dissected all the tissues.

Y.Q. performed the nuclei extraction, FACS, and 10x library preparation for all the snRNA-seq experiments with the supervision of H.L.

U.P. assembled the annotation file.

O.V.G. aligned data to the genome and performed quality control filtering with the supervision of R.S.

O.V.G. analyzed all the data with the supervision of R.S.

Y.J. carried out the differential expression analysis in Data S2, and Data S4. with the supervision of T.R.S.

H.W.-C. carried out the experiment in Figure 2C-G.

S.-C.W. performed the analysis for Figure 2K with the supervision of O.S.A.

L.D. and L.B.D. provided gene lists for circadian- and neuropeptide-related genes.

A.E.D. carried out experiments in Figures 3F-K, 4E-F, and Data S3 with the supervision of N.S.

O.V.G., A.E.D. and N.S. annotated the data with expert input from B.R.H., L.B.D., H.W-C., T.R.S. and all members of the Mosquito Cell Atlas Consortium.

Z.M.P. and T.R.S. performed the SAMap analysis.

B.W. imported the data onto the UCSC Cell Browser with the supervision of M.H.

H.L. and L.B.V. provided funding resources for this project.

O.V.G., A.E.D., L.B.V. and N.S. together designed the figures and wrote the paper with input from all authors.

## Open access statement

This article is subject to HHMI’s Open Access to Publications policy. HHMI lab heads have previously granted a nonexclusive CC BY 4.0 license to the public and a sublicensable license to HHMI in their research articles. Pursuant to those licenses, the author-accepted manuscript of this article can be made freely available under a CC BY 4.0 license immediately upon publication.

## STAR**★**Methods

### Key resources table

| REAGENT or RESOURCE | SOURCE | IDENTIFIER |
| --- | --- | --- |
| **Chemicals, peptides, and recombinant proteins** |  |  |
| Schneider's Medium | Gibco | Cat#21720024 |
| Hoechst-33342 | Invitrogen | Cat#H3570 |
| Hoechst-33342 | ThermoFisher | Cat#H3570 |
| Chitinase | Sigma-Aldrich | Cat#C6137 |
| Chymotrypsin | Sigma-Aldrich | Cat#CHY5S |
| Slowfade Diamond | Thermo Fisher | Cat#S36972 |
| Triton X-100, 10% | Sigma-Aldrich | Cat#93443 |
| Triton X-100, 10% | VWR | Cat#97063-996 |
| HEPES, 1M | Sigma-Aldrich | Cat#H0887 |
| DAPI | Sigma-Aldrich | Cat#D9542 |
| USB Dithiothreitol (DTT), 100 mM | Thermo Fisher Scientific | Cat#707265ML |
| MgCl<sub>2</sub> | MilliporeSigma | Cat#63069 |
| Low TE Buffer (10 mM Tris-HCl pH 8.0, 0.1 mM EDTA) | Thermo Fisher Scientific | Cat#12090-015 |
| Buffer EB | QIAGEN | Cat#19086 |
| 50x Protease Inhibitor Cocktail | Promega | Cat#G6521 |
| RNasin Plus Ribonuclease Inhibitor | Promega | Cat#N2615 |
| Falcon 5 mL Round Bottom Polystyrene Test Tube, with Cell Strainer Snap Cap | Corning | Cat#352235 |
| FLOWMI 40 mM Cell Strainers | Bel-Art | Cat#H13680-0040 |
| Kimble Pellet Pestle | Grainger | Cat#6HAY5 |
| Kimble Pellet Pestle Motor | Grainger | Cat#6HAZ6 |
| Wheaton Dounce Tissue Grinder (1 mL) | DWK Life Sciences | Cat#357538 |
| Beckman Coulter SPRiselect | Beckman Coulter | Cat#B23318 |
| C-Chip Disposable Hemacytometers by SKC, Inc | Fisher Scientific | Cat#22-600-100 |
| PBS, pH 7.4, 10X | Fisher Scientific | Cat#70011044 |
| Water Molecular Biology Grade, 500ML | Fisher Scientific | Cat#AAJ71786AP |
| UltraPure Bovine Serum Albumin (BSA, 50 mg/ml) | Thermo Fisher Scientific | Cat#AM2616 |
| Tween 20 | Bio-Rad | Cat#1662404 |
| KCl, 1M, 100ml | VWR | Cat#AAJ63739-AE |
| Tris Buffer, 1M, pH8.0, 100ML | MilliporeSigma | Cat#648314-100ML |
| **Critical commercial assays** |  |  |
| Chromium Next GEM Single Cell 3' GEM, Library & Gel Bead Kit v3.1 | 10x Genomics | Cat#1000269 |
| Chromium Next GEM Chip G Single Cell Kit | 10x Genomics | Cat#1000127 |
| Single Index Kit TT Set A, 96 rxn | 10x Genomics | Cat#1000215 |
| High Sensitivity D5000 Reagents | Agilent Technologies | Cat#5067–5593 |
| HCR RNA *in situ* hybridization probes, amplifiers, buffers | Molecular Instruments | see Table S11 |
| **Deposited data** |  |  |
| *Aedes aegypti* mosquito cell atlas snRNA-sequencing data | This study | BioProject: PRJNA1223381 |
| Female *Aedes aegypti* antenna and maxillary palp snRNA-sequencing data | Herre et al.<sup>36</sup> | BioProject: PRJNA794050 |
| *Drosophila melanogaster* head snRNA-sequencing data, processed | Hongjie Li et al.<sup>18</sup> | EMBL-EBI Biostudies: E-MTAB-10519 |
| Processed data | This study; Uploaded on UCSC Cell Browser | https://mosquito.cells.ucsc.edu |
| Additional raw and processed data, plots, analysis and custom scripts | This paper; Zenodo Supplemental Data | https://doi.org/10.5281/zenodo.14890012 |
| *Aedes aegypti* LVP_AGWG AaegL5.3 genome (including mitochondrial chromosome) | Matthews et al.<sup>53</sup>, Sayers et al.<sup>179</sup> | NCBI RefSeq assembly: GCF_002204515.2 |
| VectorBase, LVP_AGWG AaegL5.3 Genome Annotations (Release 58) | Giraldo-Calderón et al.<sup>58</sup> |  |
| Manual chemoreceptor annotation | Matthews et al.<sup>53</sup> | Supplementary Data 18 (Chemoreceptor Annotation.gff3) |
| the “all translation” file from the FB2023_02 version of the FlyBase genome | Öztürk-Çolak et al.<sup>180</sup> | https://ftp.flybase.net/releases/FB2023_02/dmel_r6.51/fasta/ |
| **Experimental models: Organisms/strains** |  |  |
| *Aedes aegypti* wild-type strain: Liverpool | Leslie Vosshall lab, Laboratory of Neurogenetics and Behavior, The Rockefeller University, New York, NY, USA | N/A |
| *Aedes aegypti* wild-type strain: Liverpool (for testes RNA *in situ* hybridization) | Colin Berry Lab, School of Biosciences, Cardiff University, Cardiff, UK. | N/A |
| **Oligonucleotides** |  |  |
| RNA *in situ* hybridization probes | This paper | Table S11 |
| **Software and algorithms** |  |  |
| AGAT: Another Gff Analysis Toolkit (Version v1.4.1) | Dainat et al.<sup>181</sup> | https://zenodo.org/records/13799920 |
| GFFutils (Version 0.13) | Dale<sup>182</sup> | https://daler.github.io/gffutils/ |
| GffRead (Version 0.12.7) | Pertea and Pertea.<sup>183</sup> | https://github.com/gpertea/gffread |
| Cell Ranger (version 7.1.0) | Zheng et al.<sup>63</sup> | https://support.10xgenomics.com/single-cell-gene-expression/software/pipelines/latest/what-is-cell-ranger |
| CellBender | Fleming et al.<sup>64</sup> | https://github.com/broadinstitute/CellBender |
| SCANPY (Version 1.9.6) | Wolf et al.<sup>54</sup> | https://github.com/theislab/scanpy |
| Python (Version 3.8) | Python Software Foundation<sup>184</sup> | https://www.python.org/ |
| Jupyter Notebooks | Kluyver et al.<sup>185</sup> | https://jupyter.org/ |
| NumPy (Version 1.24.4) | Harris et al.<sup>186</sup> | https://numpy.org/ |
| pandas (Version 1.3.1) | The pandas development team<sup>187</sup> | https://pandas.pydata.org/ |
| Matplotlib (Version 1.3.1) | Hunter<sup>188</sup> | https://matplotlib.org/ |
| Scrublet (Version 0.2.3) | Wolock et al.<sup>189</sup> | https://github.com/AllonKleinLab/scrublet |
| PhenoGraph (Version 1.5.7) | Levine et al.<sup>190</sup> | https://github.com/jacoblevine/PhenoGraph |
| SciPy (Version 1.10.1) | Virtanen et al.<sup>191</sup> | https://scipy.org/ |
| seaborn (Version 0.12.2) | Waskom et al.<sup>192</sup> | https://seaborn.pydata.org/ |
| Ensembl Metazoa BioMart, release 56 | Cunningham et al.<sup>56</sup> | https://feb2023-metazoa.ensembl.org/index.html |
| BLAST | Altschul et al.<sup>57</sup> | https://blast.ncbi.nlm.nih.gov/Blast.cgi |
| VectorBase (orthology tool) | Giraldo-Calderón et al.<sup>58</sup> | https://vectorbase.org/vectorbase/app |
| MAST (Version 1.26.0) | Final et al.<sup>55</sup> | https://github.com/RGLab/MAST |
| anndata (Version 0.9.2) | Virshup et al.<sup>193</sup> | https://github.com/scverse/anndata |
| batchelor (Version 1.16.0) | Haghverdi et al.<sup>194</sup> | https://www.bioconductor.org/packages/release/bioc/html/batchelor.html |
| SAMap (Version 1.0.15) | Tarashansky et al.<sup>148</sup> | https://github.com/atarashansky/SAMap |
| UCSC Cell Browser | Speir et al.<sup>62</sup> | https://www.cells.ucsc.edu/ |
| Fiji | Schindelin et al.<sup>195</sup> | https://imagej.net/software/fiji/ |
| Zen Blue (Version 3.5) | Zeiss | N/A |
| Olympus cellSens | Olympus | N/A |

### Experimental model and study participant details

#### Human and animal ethics statement

Blood-feeding procedures and behavioral experiments with live hosts were approved and monitored by The Rockefeller University Institutional Animal Care and Use Committee (IACUC protocol 23040 (PRV 20068)) and Institutional Review Board (IRB protocol LV-0652), respectively. Human participants gave their written informed consent to participate in this study.

### Mosquito rearing and maintenance

*Aedes aegypti* wild-type (Liverpool) mosquitoes were reared in an environmental chamber maintained at 26°C ± 2°C with 70-80% humidity with a photoperiod of 14 h light: 10 h dark as previously described^125^. Embryos were hatched in 1 L hatching broth: one tablet of powdered Tetramin (TetraMin Tropical Tablets 16110M) in 1 L of deionized water, then autoclaved. Larvae were reared in deionized water (3 L total) and fed 3 crushed Tetramin tablets on the first day post hatching and 2 tablets daily thereafter. To maintain low rearing density, approximately 400 larvae were kept in 3 L deionized water from L3-L4 stage. Adult mosquitoes were supplied with unlimited access to 10% sucrose solution (w/v in deionized water), delivered in a glass bottle (Fisher Scientific FB02911944) with a cotton dental wick (Richmond Dental 201205), and were kept in 30 cm^3^ BugDorm-1 Insect Rearing Cages (BugDorm DP1000). Animals were dissected on day 7 of adulthood (14 days post-hatching). All animals were co-housed to allow mating prior to dissection.

## Method details

### Photographs of mosquito tissues

7-14 day-old mosquitoes were cold-anesthetized and kept on ice. The indicated tissues were freshly dissected using using Dumont #5 Forceps (Fine Science Tools 11295-10/11295-20 or Roboz Surgical RS-4955) ice in 1 X PBS (Thermo Fisher Scientific AM9625). Only brains were pre-fixed in 4% paraformaldehyde (Electron Microscopy Sciences 15710-S) in 1X PBS, 0.25% Triton X-100 prior to dissection for 3 h at 4°C. Tissues were placed on a stage micrometer (Fine Science Tools 29025-01) and photographed using an iPhone X (Apple) through the iDu Optics LabCam adapter (iDu Optics) attached to the eyepiece of a Nikon SMZ1500 stereo zoom microscope (Nikon). A scale bar of 500 µM was added to the images using the stage micrometer’s scale (Fine Science Tools 29025-01).

### Tissue collection

Adult wild-type (Liverpool) mosquitoes aged 7 days were aspirated using an oral aspirator (John W. Hock Company 612) into a 16 ounce container (Webstaurant KH16A-J8000) and were sealed using double 0.8 mm polyester mosquito netting (ahh.biz F03A-PONO-MOSQ-M008-WT) then anesthetized on ice for 10 minutes. Mosquitoes were then placed in a 40 μm cell strainer (Falcon 352340) in a 100 mm Petri dish (Corning 430293) and soaked in ice-cold molecular-grade 100% ethanol (Sigma-Aldrich E7023 or Fisher Scientific BP2818500) for 5-10 seconds. The animals were rinsed in ice-cold Schneider’s Medium (Gibco 21720024) and placed in a clean Petri dish with approximately 20 mL ice-cold Schneider’s Medium on a reusable ice pack (GenTap, Cooler Shock. Amazon.com 854850006121). Tissues of interest were dissected using Dumont #5 Forceps (Fine Science Tools 11295-10/11295-20 or Roboz Surgical RS-4955) on a 100 mm Petri dish (Corning 430293) lined with or without SYLGARD 184 silicone (World Precision Instruments SYLG184). Tissues were placed directly into a DNA LoBind 1.5 mL tube (Eppendorf 022431021) pre-wet with 100 µL Schneider’s Medium on wet ice or in a 70 μm cell strainer (pluriSelect 43–10070-70) and DNA LoBind 1.5 mL tube (Eppendorf 022431021) pre-wetted with 100 µL Schneider’s Medium on ice by inverting the cell strainer over the Eppendorf tube using Dumont #5 Forceps (Fine Science Tools 11295-10/11295-20 or Roboz Surgical RS-4955) and pipetting 300 µL ice-cold Schneider’s Medium onto the strainer to expel the tissue. Each sample was collected in 90 minutes or less. The Eppendorf tube was wrapped in parafilm (Bemis Company Inc. PM996), flash-frozen in liquid nitrogen and stored at −70°C. All tissues were dissected in the Vosshall Laboratory at Rockefeller University. With the exception of two antenna samples, all samples were shipped to Baylor College of Medicine on dry ice for nuclei extraction. For individual sample information, including number of mosquitoes, dissection tube IDs, see Table S1. Further information on individual dissection tube IDs that were pooled together for samples, including start time of 90 minute dissection session is available in Table S1.

Our approach used physical dissection to obtain tissue for snRNA-seq, rather than generating genetically-labeled strains and isolating cells by fluorescent marker expression. This somewhat limited our ability to finely subdivide the mosquito into the largest number of individual tissues and organs. For example, we did not obtain separate data from tissues such as the heart, male salivary gland, or hemocytes. This is a common limitation in dissection-based whole animal single-cell atlases.

### Human arm feeding for blood-fed brain samples

Approximately 30 4-7 day old female mated adults were aspirated into a 30 cm^3^ BugDorm-1 Insect Rearing cage (BugDorm DP1000) and allowed to feed on a human arm for 20-30 minutes. One human subject was used for all blood feeding. Fed females were placed in an environmental chamber maintained at 26°C ± 2°C with 70-80% humidity with unlimited access to 10% sucrose solution until they reached 7 days of adulthood and were dissected. Brain dissections and collections were performed as described above.

### Single-nucleus suspension

Single-nucleus suspensions were prepared as described previously^196^. Thawed samples were spun down using the bench-top centrifuge, removing the Schneider’s medium as much as possible. Samples of like tissues were combined into one tube using a pipette with wide-bore tips and then centrifuged. Specific large tissues (abdomen, thorax and abdominal pelt) were ground using a pestle motor (Kimble 6HAZ6) for 30 seconds on ice after thawing (Table S1).

Samples were resuspended in 900 µL of fresh homogenization buffer (250 mM sucrose, 10 mM Tris PH 8.0, 25 mM KCl, 5 mM MgCl_2_, 0.1% Triton-x 100, 0.5% RNasin Plus, protease inhibitor, 0.1 mM DTT in 10 mL nuclease-free water) and transferred into a 1 mL Dounce (Wheaton 357538). Sample tubes were rinsed in 100 µL of homogenization buffer and transferred into the same dounce. Dounce sets were autoclaved at 200°C for more than 2 hours before each use.

Nuclei were released by 20 strokes of loose dounce pestle and 40 strokes of tight dounce pestle. 1000 µL of the samples were filtered into a 5 mL tube through 35 µM cell strainer cap (Corning 352235) and then filtered using Flowmi (40 µM; BelArt H136800040) into a 1.5 mL Eppendorf tube. After 10 minutes of centrifuging at 1000g at 4°C the pellet was resuspended using 500 µL of 1xPBS/0.5% BSA with RNase inhibitor (9.5 mL 1x PBS, 0.5 mL 10% BSA, 50 µl RNasin Plus). Mechanical nucleus extraction is typically estimated to recover approximately 10% of the total nuclei from tissue samples, as previously documented^197^.

### Fluorescence-activated cell sorting (FACS)

Samples that underwent FACS were filtered using a 40 µm Flowmi into a new 5 mL FACS tube (Corning 352052) and kept on ice. 10 µL of the sample was moved into a new 5 mL FACS tube with 190 µL PBS as unstained control for FACS. Nuclei were stained with Hoechst-33342 (Invitrogen H3570) on wet ice (1:1000; >5 min).

Hoechst-positive nuclei were collected using the BD FACSAria III Cell Sorter (BD Biosciences). 80k–150k individual nuclei were collected into one 1.5 mL RNAse-free Eppendorf tube with 300-500 µL 1x PBS with 0.5% BSA as the receiving buffer (with RNase inhibitor). Next, nuclei were centrifuged for 10 min at 1000 g at 4°C, and resuspended using 30 µL of 1x PBS with 0.5% BSA (with RNase inhibitor). FACS files, including gating strategy, are available in Zenodo Supplemental Data (file names for each sample listed in Table S1).

### Nuclei counting

2 µL of the nucleus suspension was used to calculate the concentration on a hemocytometer (Fisher Scientific 22-600-100). Because we have found capture efficiency to be typically around 50-65% (Table S1), 20,000 nuclei per sample were loaded on the Chromium 10x Controller (10x Genomics) to recover approximately 10,000 cells after sequencing. Excess nuclei were discarded. The nuclei capture protocol used (10x Genomics) has a limit of approximately 10,000 nuclei per sample. We note that the number of nuclei recovered is not a reflection of the nuclei yield from the original tissue amount. For most samples, we collected more tissue and extracted more nuclei than necessary to ensure availability at this step was not a limitation towards the final number of nuclei we recovered.

For specific tissues with limited nuclei (male and female malpighian tubules, male wings and female stylet), we maximized nuclei yield by using all Hoechst-positive nuclei from single-nucleus suspensions were collected with FACS, and the counting step was skipped. Nuclei suspension concentration for loading was estimated via FACS (Table S1).

### Library preparation and sequencing

10x Genomics sequencing libraries were prepared following the standard protocol from Chromium Next GEM Single Cell 3’ GEM, Library & Gel Bead Kit v3.1 (10x Genomics 1000269) with the following settings. All PCR reactions were performed using C1000 Touch Thermal cycler with 96-Deep Well Reaction Module (BioRad 1851197). Cycle numbers were used as recommended in 10x protocol for cDNA amplification and sample index PCR. As per 10x protocol, 1:10 dilutions of amplified cDNA were evaluated using a Qubit fluorometer (Thermo Fisher). Final libraries were evaluated using TapeStation (Agilent). The final libraries were sent to Novogene Corporation Inc. (Sacramento, California, USA) for Illumina NovaSeq PE150 S4 lane sequencing with the dual index configuration Read 1 28 cycles, Index 1 (i7) 10 cycles, Index 2 (i5) 10 cycles and Read 2 91 cycles. Given our target of 10,000 cells per sample, we aimed for a sequencing depth of approximately 80,000 reads per nucleus (Table S1).

### Antenna samples prepared without FACS

All samples were prepared with the above protocol, with the exception of two female antenna samples (see Table S1 and Figure S3C). These samples were not FACS-sorted prior to library creation. Methods of data collection for these samples is published in Herre, Goldman et al. 2022 (“Rockefeller” sample)^36^. The concentration of nuclei was determined by counting cells on a Luna FX7 automated cell counter (Logos Biosystems L70001) prior to library creation. Libraries were created using the standard protocol from Chromium Next GEM Single Cell 3’ GEM, Library & Gel Bead Kit v3.1 (10x Genomics 1000269). Images from automated cell counting are available in Zenodo Supplemental Data.

### Testes RNA in situ hybridization and imaging

Hybridization chain reaction RNA fluorescence *in situ* hybridization (RNA *in situ* hybridization) was conducted in whole male testes to detect RNA, using an adaptation of published protocols^198,199^. 1-3 days old adult male *Aedes aegypti* wild-type (Liverpool) were anesthetized at 4°C for 10 minutes. Testes were dissected from male mosquitoes in ice-cold PBST (1X PBS, 0.1% Tween-20) with 0.5% formaldehyde using Dumont biology tweezers (Agar Scientific T5291). The terminal abdomen was removed by grasping the upper abdomen and genitalia with separate pairs of forceps. Testes and male genital tract were cleaned of excess fat tissue. Dissected testes were fixed in 4% paraformaldehyde (made from 40% stock: 0.368 g paraformaldehyde, 1 mL RNase-free water, 7 µL 2N KOH, heated until dissolved and filtered through 0.3 µm filter) in PBST for 30 minutes at room temperature. Samples were washed twice in PBST for 5-10 minutes each, then dehydrated in 100% methanol and stored at −20°C in 100% methanol for up to 2 weeks. Prior to hybridization, samples were rehydrated by rinsing once in 70% ethanol and stored overnight at 4°C in 70% ethanol. The next day, samples were transferred to 0.2 mL PCR tubes (Azenta Life Sciences PCR1174) and rinsed twice with PBST. Samples were then pre-hybridized in 30% probe hybridization buffer (30% formamide, 5X SSC, 0.1% Tween 20, 50 μg/mL heparin, 5X Denhardt’s solution, and 10% dextran sulfate) at 37°C for 30 minutes. Probe solution was prepared by adding 0.4 µL of 100 µM probe stock to 100 µL hybridization buffer (Full list of probe sequences can be found in Data S3). Both samples and probe solution were heated to 80°C for 5 minutes before combining. Hybridization was performed overnight at 37°C in dry bath. Following hybridization, samples were washed four times for 20 minutes each in pre-warmed probe wash buffer (30% formamide, 5X SSC, 0.1% Tween 20, and 50 μg/mL heparin) at 37°C. Hairpin amplification was performed by heating 2 µL of each hairpin to 95°C for 90 seconds, cooling to room temperature for 30 minutes, then adding to 50 µL amplification buffer (5X SSC, 0.1% Tween 20, and 10% dextran sulfate). Samples were incubated in amplification buffer for 30 minutes at room temperature before overnight incubation with hairpin solution at room temperature in the dark. Samples were washed 5 times with 5X SSCT (5X SSC and 0.1% Tween 20) for 5 minutes each, followed by three 5-minute washes in 1X PBS. Tissues were then mounted in mounting medium on a cover slip and imaged. Images were acquired using an Olympus BX63 microscope (Olympus) equipped with a Cool LED pE-300 light source and Hamamatsu ORCA Spark camera (Hamamatsu Photonics C11440-36U), using 20x/0.80 UPlan XApo objective (Figures 2C-2F) or Olympus Uplan Fl 40x/0.75 objective (Figure 2G). Images were acquired as a 1920×1200 size image. Image acquisition was performed using Olympus cellSens software.

### Antennal RNA *in situ* hybridization

RNA *in situ* hybridization was conducted in whole mount female and male antenna to detect RNA using adaptations of published protocols^36,198,200^. Products including HCR custom probes, amplifiers, probe hybridization buffer, probe wash buffer, and amplification buffer were purchased from Molecular Instruments Inc. (https://www.moleclarinstruments.com). All staining steps were done in a modified cell strainer snap cap (Fisher Scientific, Falcon 352235) in a well of a 24-well plate (Fisher Scientific, Falcon 353047).14-day-old adult Liverpool mosquitoes were anesthetized on wet ice. Antennae were dissected in a bubble of ice-cold 1X PBS (Thermo Fisher Scientific AM9625) in a 100 mm Petri dish (Corning 430293) lined with SYLGARD 184 silicone (World Precision Instruments SYLG184) on a reusable ice pack (GenTap, Cooler Shock. Amazon.com 854850006121) using Dumont #5 Forceps (Fine Science Tools 11295-10/11295-20 or Roboz Surgical RS-4955). Samples were digested in a chitinase-chymotrypsin solution [119 mM NaCl, 48 mM KCl, 2 mM CaCl_2_, 2 mM MgCl_2_, 25 mM HEPES, 5 U/mL chitinase (Sigma-Aldrich C6137-50UN), 100 U/mL alpha-chymotrypsin (Sigma-Aldrich CHY5S-10VL), 1% DMSO] rotating at 37°C for 1.5 hours. Antennae were washed in 1% PBS, 0.1% Tween-20 (PBST) for 10 minutes three times at room temperature. Samples were then fixed in 4% paraformaldehyde (Electron Microscopy Sciences 15710-S) in 1X PBS, 0.025% Triton X-100 for two hours at room temperature, following six five-minute washes at room temperature in PBST. Antennae were then dehydrated at 4°C in a stepwise sequence of 25% methanol/PBST, 50% methanol/PBST, 75% methanol/PBST, then 100% methanol twice, for 10 minutes at each step. Samples were kept in 100% methanol overnight at −20°C. The following day tissues were rehydrated at 4°C in a stepwise sequence of 75% methanol/PBST, 50% methanol/PBST, 25% methanol/PBST for 10 minutes each. At room temperature, samples were washed in PBST four times for ten minutes, fixed in 4% paraformaldehyde in PBS with 0.1% Tween for 20 minutes, and then washed again in PBST three times for 15 minutes. Antennae were transferred to preheated probe hybridization buffer at 37°C for 30 minutes. 8 µL of 1 µM stock of each probe was added to 800 µL of preheated probe hybridization buffer at 37°C, samples were transferred to this probe solution for two nights and kept at 37°C (Full list of probes can be found in Data S3). They were then washed four times for 15 minutes at 37°C in probe wash buffer, followed by four 15-minute washes in 5X SSC (Invitrogen 15557044) in nuclease-free water, 0.1% Tween 20 solution (SSCT) at room temperature. Antennae were then incubated in amplification buffer for 30 minutes at room temperature. Hairpin amplifiers were combined and activated per the manufacturer’s instructions. 8 µL of 3 μM stock hairpins were added to 800 µL of amplification buffer at room temperature overnight in the dark. At room temperature, samples were washed in SSCT twice for 15 minutes, incubated in 1:500 DAPI (Sigma-Aldrich D9542-5MG) in SSCT for one hour, then washed again in SSCT five times for 15 minutes. Tissues were then mounted on slides in SlowFade Diamond (Thermo Fisher S36972), topped with a coverslip, sealed with clear nail polish, and stored at 4°C until imaged.

### Antennal imaging and image processing

Confocal images of antennae were acquired on a Zeiss Axio Observer 7 Inverted LSM 980 scanning confocal microscope (Zeiss) with a 63x/1.40 PlanApochromat Oil DIC M27 objective. The sample was scanned bidirectionally without averaging (Figures 4E-4F and Data S3) or with 4x averaging (Figures 3F-3K). The images were acquired as a standard 1024×1024 size image, which, depending on the zoom used, resulted in a voxel size of 0.0658 μm x 0.0658 μm x 0.24 μm (for Figures 3F-3K) or 0.1315 μm x 0.1315 μm x 0.26 μm (for Figures 4E-4F and Data S3). Zen Blue v3.5 software was used for image acquisition.

For all comparative experiments, image acquisition parameters were kept consistent. We note that all confocal imaging was conducted in a manner that would maximize our ability to visualize the presence or absence of each fluorophore and was not intended as a quantitative measure of fluorescence intensity. Confocal images were processed in ImageJ (NIH). Brightness/contrast was adjusted to maximize visualization, and for all comparative experiments, adjusted parameters were kept consistent.

## Quantification and statistical analysis

### Gene Annotation File

Gene annotations were prepared from VectorBase (www.vectorbase.org, Release 58, as of June 2022) using the *Aedes aegypti* LVP_AGWG AaegL5.3 Genome^53,58^. These were merged with the manual chemoreceptor annotation from^53^, then double checked and corrected for errors manually as well as using AGAT^181^ and then processed using gffread^183^. For quick identification in downstream analyses, the prefixes “MT-”, “RP-” and “RR-” were appended to all AAEL gene IDs for mitochondrial, ribosomal protein, and rRNA genes, respectively. Final annotation file was assembled using Cell Ranger package (version 7.1.0) function *mkgtf*^63^ using the *Aedes aegypti* genome, including the mitochondrial chromosome, downloaded from NCBI (NCBI RefSeq assembly: GCF_002204515.2)^53,201^. Gene annotation file (including prefixes identifying MT, PR, and RR genes) is available in Zenodo Supplemental Data.

### Alignment and ambient RNA removal

FASTQ files were aligned using 10x Genomics Cell Ranger 7.1.0 (include-introns set to “true”)^63^. While the Cell Ranger performs alignment, PCR duplication correction and identification of empty droplets, the cells are susceptible to ambient RNA noise. A droplet containing a nucleus may also contain remnant floating RNA, which can occlude the nucleus’ expression. We therefore used the CellBender package^64^ for ambient RNA correction (*epochs=200, fpr=0.01*). We used the Cell Ranger cell count estimate as the number of expected cells and set the number of total droplets to the recommended default value (generally 30,000 droplets for typical samples). We selected the learning rate based on the smoothness of ELBO value along the epochs, as suggested by the developers. For most cases, we used the default learning rate and in cases where the ELBO value was “wobbly” we chose x0.1, x0.5 or x0.01 as suggested in the CellBender package^64^. A list of parameter values is provided in Table S1 and scripts used for Cell Ranger and CellBender are available in Zenodo Supplemental Data.

Prior to ambient RNA removal, on average for each sample, we sequenced 115,918 reads/nucleus, a median of 1,417 genes/nucleus and 3,809 UMI counts/nucleus, with a sequence saturation of 82% (for metrics on individual samples, see Table S1). On average across an entire sample, we detected 16,285 out of 19,920 annotated genes (Table S1).

### Quality control and cell filtering

For all downstream analysis, we used the Scanpy package (referred to as sc from here on^54^, in Python^184,202^ in addition to standard Python libraries such as numpy, pandas, matplotlib, csv, os, datetime^186–188^. Most analysis was carried out in Jupyter notebooks^185^, and all scripts and additional data are available on Zenodo Supplemental Data.

#### Quality control metrics

We began by evaluating basic quality control metrics using *calculate_qc_metrics* function in Scanpy in each sample. We evaluated the distribution of each metric such as the total counts in a cell, total number of genes expressed in a cell and the number of cells each gene is expressed in to filter for high quality cells and genes. We also evaluated Mitochondrial (MT), rRNA (RR), and ribosomal protein (RP) fractional expression distribution across cells. These metrics are associated with apoptotic cells or are typically uninformative^203^, hence understanding their contribution to the expression of each cell is important. To err on the conservative side, we began by removing only a few cells that were clearly noisy or outliers. Specific parameters and scripts for each sample are in Table S1 and Zenodo Supplemental Data.

We also performed basic filtering in the gene space. First, as a standard practice in the analysis of scRNA-seq data, we removed RP genes from downstream computation, as they are typically uninformative and are often confounders in biological signals^203^. Additionally, to reduce noise in the data, genes that were expressed in fewer than 12 cells were also removed, unless they were registered as possibly biologically meaningful after discussion with Aedes aegypti Mosquito Cell Atlas co-authors. For this, we compiled a list of around 2,464 genes that were of interest based on the current literature (see Zenodo Supplemental Data).

#### Data Normalization

After basic clean-up, each sample was median library size normalized followed by log-transformation, which was recently shown to perform just as well, if not better, than more sophisticated transformations^204^. We used *sc.pp.normalize_total* function in Scanpy and took the natural logarithm of the data with a pseudocount of 1 to preserve zeroes. We then computed the top 4000 highly variable genes (*sc.pp.highly_variable_genes*), followed by a principal component analysis (PCA, 30 components). We then computed k-nearest neighbors using *sc.pp.neighbors(n_neighbors=30, use_rep=’X_pca’, metric=’euclidean’)* function in Scanpy. UMAP, tSNE, Force Directed Layout (FDL) visualizations were used for visualization of data in 2D.

#### Doublet detection

For doublet detection, we used the scrublet package^189^. Scrublet expects an estimate of doublets as an input, for which we used the formula *y=0.000759x+0.052721* from the expected multiplet table provided by 10x Genomics, where *x* is the total number of cells in the dataframe. The predicted doublets were then analyzed together with other quality metrics for data clean-up as described below.

#### Cell type-informed data filtering

Combining all the metrics discussed above, cell filtering was performed through identification of low quality clusters. A typical strategy to filter individual cells relies on individual metrics such as library size or doublet score, which can be manual and less generalizable. We instead sought to utilize the entire transcriptome to first group cells and filter out clusters of cells that cumulatively have low quality scores for the above described set of metrics: doublet score, mitochondrial gene fraction, ribosomal protein fraction, total counts, gene counts and cell type-specific gene expression. We removed clusters of cells that demonstrated low quality features (Table S1). To do this systematically, we first identified obvious outlier clusters, using which we defined a threshold that was uniformly applied to all clusters in each sample. For clustering we used the PhenoGraph^190^ package with the Leiden algorithm (*resolution_parameter=5* or 10, see Table S1) as implemented in the *sc.external* module. We chose such a high value of resolution_parameter, which results in a large number of clusters, to ensure that only highly specific noisy clusters were removed from downstream analysis. At minimum, clusters from all samples were removed that had a mitochondrial gene fraction of 5 or higher, and a doublet score of 0.3 or higher (Table S1). In many cases, these thresholds were adjusted based on their distribution to retain only high-quality cells for downstream analysis (see individual sample scripts in Zenodo Supplemental Data), because low-quality cells confound the characterization of real biological features in the data. Thus, we prioritized our analysis on high quality cells to enhance our understanding of these uncharacterized cell types with minimal exceptions (see testes data below), Because there is limited prior knowledge on basic quality metrics for single-nuclei data from mosquitoes, we sought to biologically guide and complement our cell-filtering strategy using whatever limited information we have about cell type markers in mosquitoes. For the purposes of a preliminary annotation to inform cell filtering strategy, we queried genes that appear often in most samples and utilized those to represent broad cell type categories. In particular, we used *AAEL024921* (*nSyb*) for neurons, *AAEL027131* (*repo*) for glia, *AAEL019468* (*Ppn*) for hemocytes, *AAEL001168* (*grh*) for epithelial-like cells, *AAEL002417* (*troponin T*) for muscle, *AAEL001194* (*FASN1*) for fat cells. We also included *AAEL019457* (*Lim1*), a commonly expressed transcription factor, that typically labels a discrete subset of cells. Cells that expressed these genes were typically not removed in filtering and used as reference for setting thresholds (described above) and identifying outlier clusters (Table S1).

We note that *grh* is an orthologue of a *Drosophila melanogaster* transcription factor that is not exclusive to epithelial cells, but serves as an epithelial cell marker^18,123,205^. Similarly, *Drosophila melanogaster snu* encodes an ABC transporter involved in cuticle development^206^ and also serves as a non-exclusive epithelial cell marker^207^. In the absence of functional validation that *snu (AAEL018334)*and *grh* are epithelial markers in *Aedes aegypti*, we took a cautious approach and retained the gene names for annotations rather than assigning definitive epithelial cell type labels.

To validate male and female samples, we also queried *Nix (AAEL022912)*, which showed markedly differential expression in male and female samples, as expected^26,208^.

Samples were then each reprocessed, which included renormalizing the data, re-computing highly variable genes, PCA, and nearest neighbors. Same sets of parameters were used. For clustering, we used the Jaccard + Louvain algorithm implementation of PhenoGraph at resolution 1 for downstream annotation and analysis, unless indicated otherwise^190,191^. Preliminary annotation was performed on each sample as described in the next section. Only one round of filtering (or cluster removal) was performed for each sample. Preliminary annotation was performed on each sample individually.

#### Testes sample

We processed the testes sample both with and without CellBender. We observed that CellBender discarded more 6,151 nuclei compared to Cell Ranger. We reasoned that these might be spermatids, which after meiosis slow transcription, and therefore have low transcript counts^68^. For this reason, to detect spermatids in our snRNA-seq data, we did not apply ambient RNA removal and did not discard clusters with features such as low UMI-count (Data S1). Spermatids were readily identifiable by their low transcript count and their expression of *S-Lap* (*AAEL000108*), *DBF4* (*AAEL008779*)^70^, and *Orco* (*AAEL005776*)^71^ (Figures 2B and Data S1). To avoid potential batch effects (from lack of ambient RNA removal) in the integrated *Aedes aegypti* Mosquito Cell Atlas object (Figures 1C-1F and S2), we used the testes data that were processed with ambient RNA removal and thus lacks spermatids. Testes data processed without CellBender are available on UCSC Cell Browser (http://mosquito.cells.ucsc.edu) and with CellBender at Zenodo Supplemental Data.

We note that the testes sample (without CellBender), as well as others, including those collected from male reproductive glands, and the male and female malpighian tubules, have low fraction reads in cells (Table S1). This may be due to technical artifacts within these particular samples that lead to increased ambient RNA or may indicate the possibility of additional low RNA-count cell populations. Without biological reasoning (such as in the case of spermatids) and specific gene markers to identify low RNA-count cell populations, we did not investigate these other potential cell populations.

We also observed the expression of the taste receptor *Gr39* during the cyst cell developmental trajectory, underscoring the largely uncharacterized presence of chemoreceptors in non-sensory tissues in *Aedes aegypti* and other insects (Data S1)^209–213^.

### Sample merging

In cases where we had multiple samples for a tissue, we merged data, including male and female samples. In general, for a more robust annotation of cell types informed by a greater number of cells, and to enable comparison across sexes, we merged male and female samples. Our preliminary annotations (see Methods: Annotations and gene selection) showed in most cases a noticeable similarity in general cell types in male and female samples. We used *AnnData.concatenate* function^193^ and repeated the processing as described above. Genes expressed in fewer than 18 cells were removed unless they were present in a more comprehensive list of genes of interest (20,587 genes, see Zenodo Supplemental Data). We then renormalized the data, re-computed highly variable genes, principal components (PCs), nearest neighbors, and re-clustered as described above, unless indicated otherwise. See Table S1 for list of final objects. All objects available through either UCSC Cell Browser (https://mosquito.cells.ucsc.edu) or Zenodo Supplemental Data.

### Batch correction

In the case of the two merged ovary samples, we saw a noticeable batch effect of unknown origin (Data S1). We batch-corrected all genes using *batchelor.fastmnn*^194^. Quality control and filtering was done iteratively and informed by annotations on individual and merged samples. It was not necessary to batch correct any other samples for our other analyses.

### Annotations and gene selection

In non-model organisms, lack of knowledge of expected cell types, absence of extensive gene characterization, and few established cell markers, makes cell type annotation challenging. Prior to analysis and annotation, we contacted an international group of mosquito experts to solicit hypotheses about putative cell types, as well as potential cell markers or genes of interest. *Aedes aegypti* genes came from sources including previous mosquito literature and previous bioinformatics analyses assessing putative function or gene families from the AaegL5 genome.

#### Gene orthology to Drosophila melanogaster

In addition to information collected from the *Aedes aegypti* Mosquito Cell Atlas Consortium, we used information from homologues in *Drosophila melanogaster* that have been better-characterized. Orthologous genes were assessed using Ensembl Metazoa BioMart database (Ensembl Genomes release 56^56^, BLAST (nucleotide or protein)^57^, or VectorBase^58^. We also used curated and computed cell marker genes from the Fly Cell Atlas^18^. It is important to note that *Aedes aegypti* and *Drosophila melanogaster* are separated by 260 million years since their last common ancestor^59,61^, with distinct behaviors, life cycles and physiology, so relying on *Drosophila melanogaster* homology to interpret *Aedes aegypti* genes can be problematic.

For instance, in a comparative genomic study of *Drosophila melanogaster* and several mosquito species of developmental genes, while many were well-conserved, key developmental genes in *Drosophila melanogaster* (as well as other insects) were not identified in mosquito genomes^59^. The fibroblast growth factor (FGF) signaling pathway involved in many biological processes including cell differentiation and migration, is conserved between flies and vertebrates, but was not identified in mosquito species.

Additionally, cases were also observed of increased copy numbers of developmental genes in mosquitoes. How these individual copies differ from their homolog in *Drosophila melanogaster* is not known. While some genes and pathways are conserved, divergence in gene function and expression patterns is also expected, which can easily lead to misinterpretation and errors in analysis if one relies too heavily on *Drosophila melanogaster* to benchmark discoveries in *Aedes aegypti*.

#### Gene marker selection

Genes were selected based on *sc.tl.rank_genes_groups* and MAST on Louvain or Leiden algorithm clustering^54,55^. Top computed marker genes for each cluster were each assessed visually (UMAP) and by comparing average gene expression across all clusters in the data object. Genes were manually selected based on their ability to confer information of cell type, orthology to known *Drosophila melanogaster* marker genes, and their distinctiveness as a marker gene across cell types in all datasets. For instance, the transcription factor *Sox100B* (*AAEL009473*) was used as a marker and commonly observed in sensory tissues. Recent work identifying these cells in *Drosophila melanogaster* tarsi suggests that *Sox100B-*expressing cells may be important for neural lamella formation^137^.

Computed gene markers for Louvain or Leiden algorithms used for annotations using *rank_genes_groups* (integrated data object of all sugar-fed cells) and MAST (all other tissues) are available on Zenodo Supplemental Data. Computed gene markers on manual annotations for all tissues using *rank_genes_groups* are available on UCSC Cell Browser and Zenodo Supplemental Data.

#### Annotation using gene markers

Annotations were performed using a combination of semi-automated and manual methods. Principal annotations were performed on each tissue (Figures 2, 3, 6, and Data S2) and on the integrated data object of all sugar-fed cells (Figure 1). Preliminary annotations were performed on each sample individually before merging all samples of each tissue (Table S1). Data were clustered using Louvain or Leiden algorithms and clusters were assigned cell type annotations. Clustering resolution was set based on cellular complexity of tissue and amount of prior information on tissue cell types (Table S1). Clusters were assessed for mean expression of identified gene markers using outputs from MAST, UMAPs, heatmaps, violin plots and bar plots (Zenodo Supplemental Data). Clusters were assigned a cell type annotation based on expression of thresholded gene markers or combinations of gene markers (Table S1, for annotation script see Zenodo Supplemental Data). Gene markers for each cell type were also assigned a threshold through assessment of mean expression levels across clusters (Table S1).

#### Sensory neuron annotations

*nompC-*negative sensory neuron populations in the antenna, maxillary palps, tarsi and proboscis were annotated separately in a similar pattern to tissues (Figures 4, 5, S7 and Data S4). We used the same combination of semi-automated and manual methods as described above, however for these populations, we attributed extra significance to a list of putative sensory genes that might affect the stimulus response profile of a given cell type (Table S1). Clusters were computed with the Leiden algorithm, at high resolution due to sensory neuron complexity (Table S1). Clusters were assigned a cell type annotation. Cell types were named for chemoreceptors uniquely expressed in a cell type.

In the antenna, despite separating clusters at high resolution (Leiden, resolution 10), we found at least 6 examples of chemoreceptor genes co-expressed within a cluster but not within the same cells. 6 examples of co-clustering but putatively distinct olfactory sensory neuron cell types include *Ir41*e and *Ir41b* (Figures 4C-4D), *Or41* and *Or16*, *Or25* and *Or33*, *Or2* and *Or14*, *Ir31a2* and *Ir75l*, *Or103* and *Or108* within *Or91*-expressing cells (all indicated with astrisks in Figure S5B). While this suggests the mutual exclusivity of these genes, we cannot rule out the possibility that it could be due to dropouts in single-cell sequencing, particularly because receptor genes can be expressed at relatively low levels, although this is unlikely given the data’s large number of cells and high sequencing depth. Furthermore, these cells occupy the same phenotype space and are not discernible as distinct clusters computationally, suggesting that these olfactory sensory cell types may be distinct, but transcriptomically similar.

### Sensory neuron analysis

For the antennae, maxillary palps, and proboscis samples, we subsetted and filtered *nompC-*negative sensory neurons for further analysis. We identified the neuronal population based on the expression of *Syt1* (*AAEL000704*), *brp* (*AAEL018153*), *nSyb* (*AAEL024921*) and *CadN* (*AAEL000597*) (Figures 4A, S6A and Data S4). We excluded mechanosensory neurons based on the expression of the *Drosophila melanogaster* orthologue of mechanosensory receptor *nompC* (*AAEL019818*) (Figures S4A-S4B). We removed clusters with a high doublet score (Figures S4C-S4D). Before reclustering, we additionally removed individual nuclei with a doublet score above 0.15 (Figure S4H).

This ensured a conservative filtering of potential doublets given our interest in possible co-expression of receptor genes. For the antenna, we also filtered on neuronal gene fraction to ensure we were only looking at high quality neuronal nuclei (Figure S4E), although we note that this step removed *Gr20* cells from our analysis (cluster 83, Figures S4A, S4C, S4G and S4R and Data S2). For wing and abdominal tip neuron subsetting, all neurons were included for assessment of putative sensory gene expression (Zenodo Supplemental Data). As with other tissues, we removed individual nuclei with a doublet score above 0.15 (Figure S4H).

### Cell type comparison across conditions/sexes

#### Cell abundance comparison

For the sexual cell type abundance difference, the frequencies of each cell type in each tissue for both sexes were determined by calculating the proportion of each cell type relative to the total number of cells in the tissue. The sexual abundance difference index for each cell type in each tissue was calculated using the following equation (Data S2, scripts in Zenodo Supplemental Data):

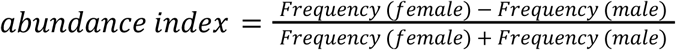

#### Sexual abundance difference index

Where cell type was categorized based on abundance difference across sexes, cells were considered “Female biased” if abundance index >0.3; “Neutral” if abundance index was −0.3 to 0.3, inclusive; “Male biased” if abundance index <-0.3. In bar plots, if there are biological replicates, the value for each replicate was shown as dots, and the standard error was calculated.

#### Differentially expressed genes

MAST was used to calculate the differentially expressed genes for cell type annotation, across sexes, and blood-feeding conditions for each cell type^55^. Log fold change is represented by MAST coefficients *(coef)*. For counting significantly differentially expressed genes in Figures 6, 7 and Data S4, MAST output files were thresholded for absolute value of *coef* above 1, and a false discovery rate of 0.05. *coef* was calculated from normalized expression (natural log). We only analyzed cell types with at least 10 cells in all conditions. In some cases, MAST *coef* could not be calculated for some genes due to their normalized-log expression being zero or close to zero in at least one of the conditions (NaN genes). NaN genes were included in differentially expressed gene counts (bar plots) if they were expressed in greater than 10% of genes in at least one condition and had normalized expression value greater than 1 (Table S2, and Zenodo Supplemental Data). Most NaN genes did not meet this criteria and were discarded. No NaN genes met this criteria for generation of volcano plots (Figures 5, 6 and S8). NaN genes were left grey for log fold change heatmaps (Figures 7 and S10).

For male versus female differential gene expression analysis across annotated cell types in Data S2, genes were discarded prior to analysis if they were not expressed in at least 10% of cells in each sex within each cell type. Differentially expressed gene counts were determined by genes that were |*coef*/log(2)| >1 and a false discovery rate <0.05.

Notably, gut enterocytes and fat tissue in the abdominal pelt showed appreciable sexual dimorphism. For gut enterocytes, both *nhe3* and *NAAT1* cell populations exhibited sex-specific expression (for *nhe3* enterocytes, 236 genes significantly upregulated in females, and 103 in males; for *NAAT1*, 65 and 80 for females and males, respectively) (Figures 1C, Data S1-2 and Table S2). Similarly, female and male abdominal pelt fat tissue also demonstrated large transcriptional differences (77 genes significantly upregulated in females and 84 in males) (Figures 1C, Data S1-2 and Table S2).

#### Volcano plots, log fold change heatmaps

Volcano plots and log fold change heatmaps on differentially expressed genes were made using MAST differentially expressed genes. Log fold change is represented by MAST coefficients *(coef)*. Volcano plots were made with *seaborn.scatterplot* on -log_10_(false discovery rate)^192^. Log fold change heatmaps using *seaborn.heatmap* on individual genes were made by identifying all clusters where the gene had a calculated false discovery rate <0.05 in at least one timepoint. Heatmaps were sorted by sum of *coef* values. Only clusters that had more than 10 cells in each timepoint were included.

### Data visualization

#### UMAPs, Gene fraction visualization

UMAP coordinates were created using *scanpy tl.umap* function on the constructed nearest neighbors graph (described above). The *min_dist* parameter used are described in Table S1. We visualized UMAPs using *sc.pl.umap* function. We quantify gene signature expression by computing gene fraction defined as: *np.asarray(np.sum(adata.X[:, index_mechano_genes], axis=1)/np.sum(adata.X, axis=1)).squeeze()*100* and visualized on UMAP. This is the mRNA content represented by the genes in the list for a given cell as a fraction of total mRNA of the cell.

#### Dot plots and heatmaps

Clusters for dot plots and heatmaps were organized using *sc.tl.dendrogram, sc.tl.heatmap* functions followed by *sc.pl.dendrogram* or *sc.pl.heatmap* functions on selected genes. Heatmaps were visualized using seaborn *sn.heatmap*^192^. Full heatmaps of all putative sensory genes expressed in selected sensory neurons are available in Zenodo Supplemental Data, in addition to wing and abdominal tip datasets. For neuropeptide-related gene heatmaps, genes were identified from literature and *Drosophila melanogaster* orthology, as previously described^56,214,215^. Genes were considered expressed by a cell type if they had a normalized expression value of at least 1 and were expressed by at least 20 percent of all cells in that cluster.

#### Violin plots, stacked bar plots, box plots

Violin plots were made using *sc.pl.violin* or *sc.pl.stacked_violin*. Proportion stacked bar plots were made using *matplotlib ax.bar*. Boxplots made with *seaborn.boxplot* and *seaborn.stripplot*.

### Diffusion component & cluster distance analysis

To quantify the transcriptomic difference between male *ppk317* and other antenna cell types, we applied diffusion components analysis (Figures S3E-S3F) using *sc.tl.diffmap*, with 80 diffusion components using the nearest neighbors graph (described above). Diffusion components have been widely used in single-cell data analysis to approximate phenotypic distances between subpopulations of cells^216–220^. Because the top diffusion components explain the most variance in the data^218,219^, we calculated the top correlating gene for the diffusion components 1 and 2 (Zenodo Supplemental Data). *ppk317* (*AAEL000873*) was the highest scoring gene of diffusion component 1 (|correlation score| >0.89) (Figure S3E, first panel). Neuronal markers including *Syt1 (AAEL000704)* and *nSyb* (*AAEL024921*) ranked highly for diffusion component 2 (for both a |correlation score| >0.72) (Figure S3E, second panel). We then selected top components based on eigengap as has been done previously^218,219^. We observed that the first major gap in eigenvalues occurred between 18th and 19th eigenvalues, as such we chose top 18 eigenvectors for further analysis: Eigenvalues 1 through 18.

Partition-based graph abstraction (*sc.tl.paga*) was then made through recalculating nearest neighbors using thus computed diffusion components (Figure S3F). For box plot in Figure S3J, pairwise Euclidean distances were computed to approximate phenotypic distance based on diffusion embeddings using *sklearn.metrics.pairwise_distances* ^221^ and plotted with *matplotlib.boxplot*.

#### Correlation matrix heatmap

To evaluate pairwise correlation of gene expression between clusters in Figure S3G, we computed the Pearson correlation coefficient matrix (*numpy.corrcoef*) between normalized gene expression matrices for every pair of clusters. We computed correlation between every pair of cells for every pair of clusters and reported the mean correlation value as a heatmap. Diagonal values (cluster to itself) represent intra-cluster correlation values, which vary based on features such as cell number and gene heterogeneity.

#### Raw counts scatterplot

To generate the scatter plot on antenna olfactory sensory neurons in Figure S5C, raw transcript counts (unique molecular identifiers) for a list of putative sensory genes (Table S1) were counted and plotted for each sample using *matplotlib.scatter*.

### Cross-species comparison

For comparison of the *Aedes aegypti* Mosquito Cell Atlas brain to the fly cell atlas (FCA) head data, we used SAMap (v1.0.15^148^). SAMap was used according to documentation. All versus all NCBI BLAST (v2.9.0^57^) was run using the SAMap script map_genes.sh on the annotated proteins from the VectorBase-58 version of LVP_AGWG genome and the “all translation” file from the FB2023_02 version of the FlyBase genome. Analyses were performed on the FCA head dataset^18^ and the *Aedes aegypti* Mosquito Cell Atlas all brain dataset. These datasets were subsetted into neurons and glia and abundant cell clusters were subsampled using scanpy. The FCA head dataset was subsetted using the FCA cell type annotation clusters. Clusters with mean expression of the gene *Dm_repo* >0.4 were considered glia and mean expression of *Dm_nSyb* >1.2 were considered neurons. Then cell clusters of neurons (Leiden algorithm, resolution = 4) with >1000 cells were subsampled down to 1000 using *scanpy.subsample*. The *Aedes aegypti* Mosquito Cell Atlas brain dataset was subsetted using the Leiden algorithm (resolution = 5) clusters. Clusters with mean expression of the gene *repo (AAEL027131)* >2.0 were considered glia and mean expression of *nSyb (AAEL024921)* >0.7 were considered neurons. Neurons clusters with >1000 cells were subsampled down to 1000 using the subsample function in Scanpy. Subsampled neurons and all glia were then run in SAMap using default parameters. FCA and *Aedes aegypti* Mosquito Cell Atlas neurons were run together, FCA and *Aedes aegypti* Mosquito Cell Atlas glia were run together, and as a control FCA glia and *Aedes aegypti* Mosquito Cell Atlas neurons were run together (Figures S9E-S9K). Mapping scores were determined between FCA cell type annotations and *Aedes aegypti* Mosquito Cell Atlas (Leiden, resolution = 5) clusters. Kenyon cells (KCs) were identified in the *Aedes aegypti* Mosquito Cell Atlas dataset by high mapping scores with the FCA KCs and expression of the known markers including *Hr51 (AAEL020847)* and *sNPF (AAEL019691)* (Figures S9L and S9M). We also queried markers for potential Kenyon cell subtypes in the *Aedes aegypti* Mosquito Cell Atlas using *Pde8* (*AAEL019528*) (alpha/beta KCs), *mamo* (*AAEL019481*) (alpha’/beta’ KCs) and *Imp1* (*AAEL006876*) (gamma KCs) (Figures S9N-S9P)^18,150^.

## Supplemental figures and legends

**Figure S1.**
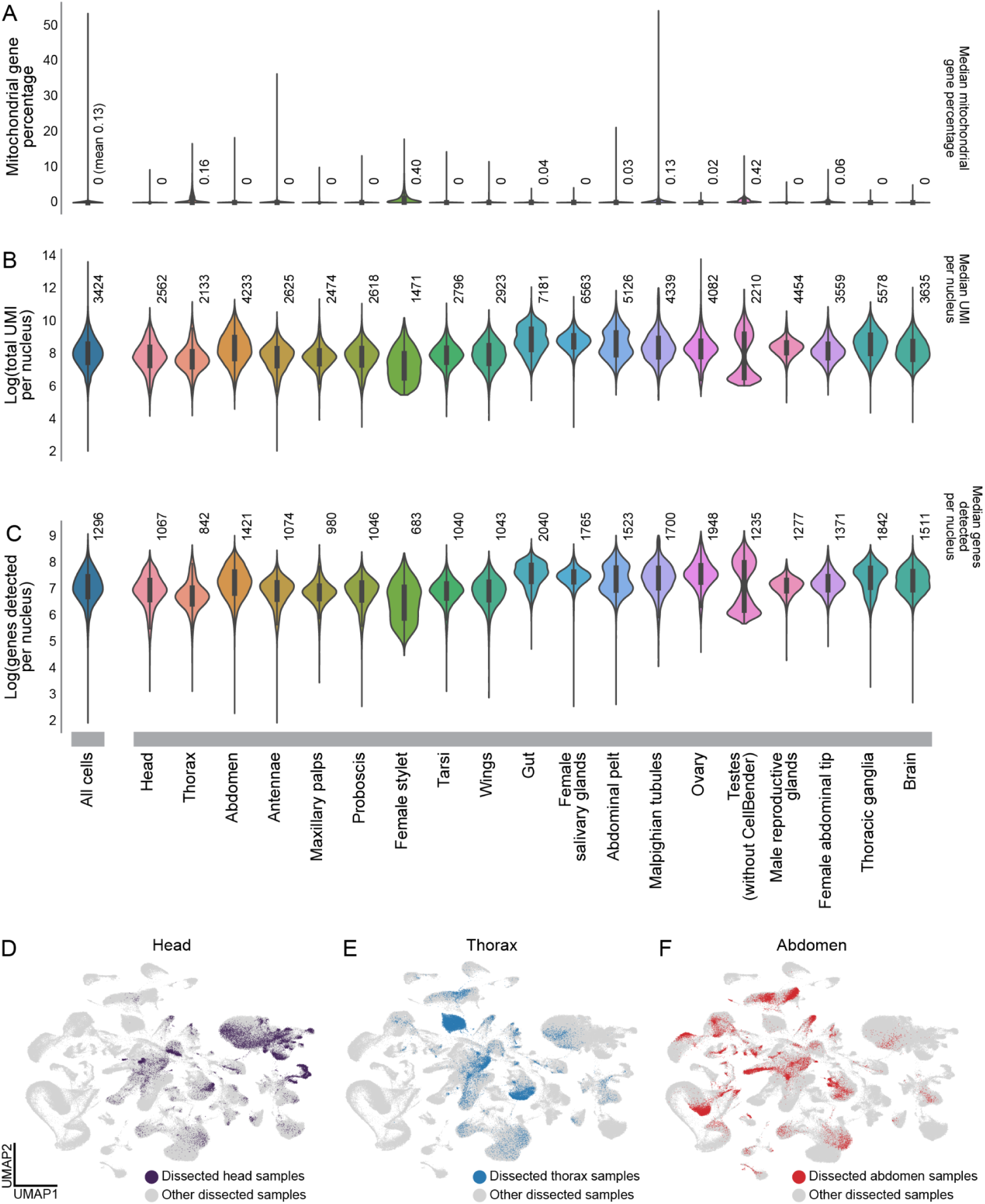
Final data quality control post filtering, related to Figure 1. **(A-C)** Violin box plots depicting data (post-quality control filtering) mitochondrial gene percentage (A), total unique molecular identifiers (UMI) per nuclei (B), and genes detected per cell (C) for all cells (left) and for each tissue (right). Across all nuclei from all tissues, median mitochondrial gene percentage was 0.00% (mean 0.13%), median total UMI per nucleus was 3,424, and median genes per nucleus was 1,296. Annotations above each violin represent median values unless otherwise indicated. Pre-filtering gene per nucleus and UMI per nucleus metrics are available in Table S1, as well as filtering parameters. Note clusters were filtered by quality control metrics on clusters, not filtered on individual cell quality control metrics (Data S1, Table S1). Multiple samples (10x Genomics libraries) from males and females are included for each tissue. Inner boxes represent first quartile to third quartile. Box plot lines represent 1.5x interquartile range. Length of violin indicates complete range of the data, with thickness of the violin representing number of cells at each value. **(D-F)** UMAPs of integrated *Aedes aegypti* Mosquito Cell Atlas data, colored by samples representing major body parts: female and male head (D), thorax (E), and abdomen (F).

**Figure S2.**
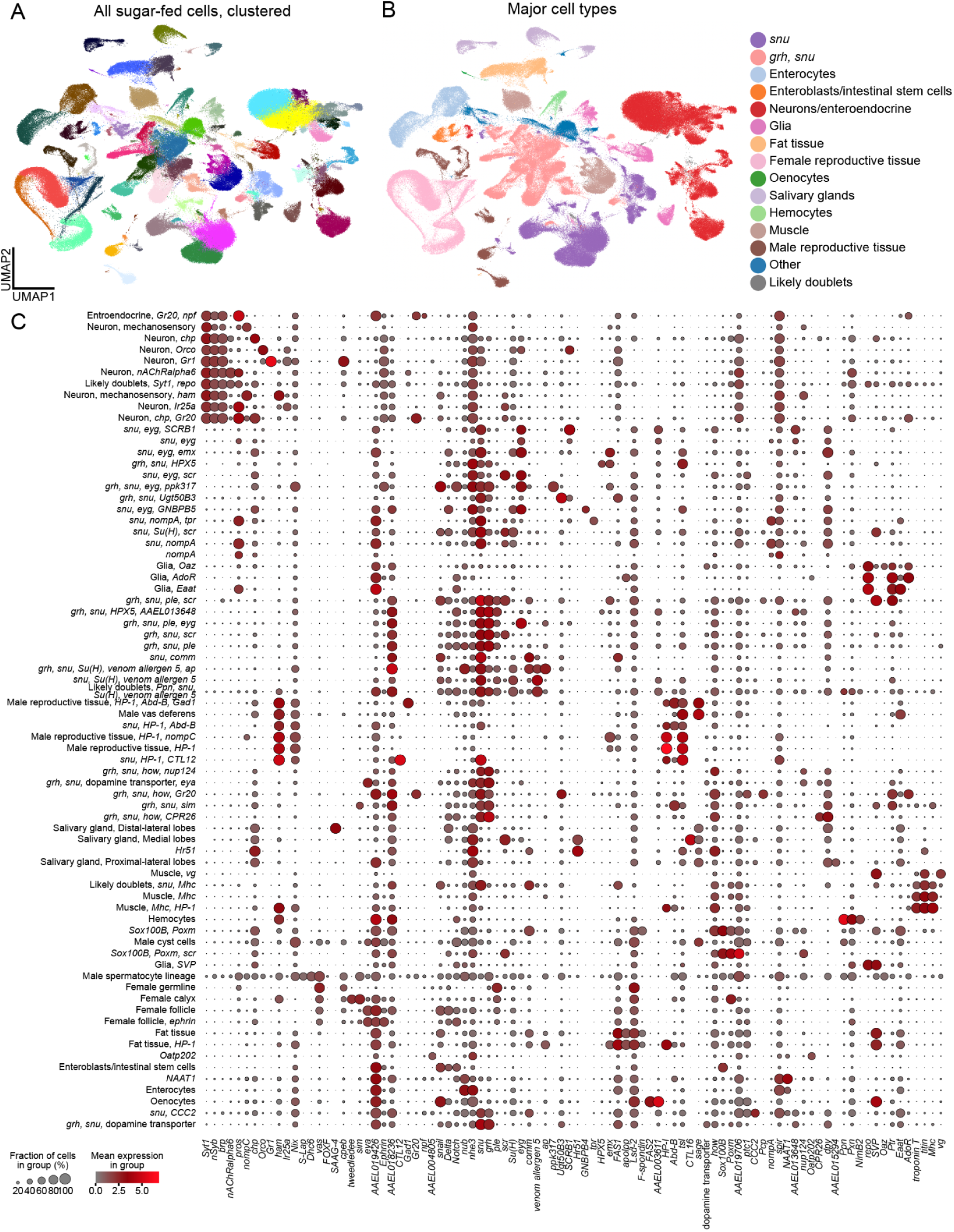
Integrated *Aedes aegypti* Mosquito Cell Atlas tissues and data, related to Figure 1. **(A-B)** UMAPs of integrated *Aedes aegypti* Mosquito Cell Atlas data colored by clustering (Louvain algorithm, resolution = 1) (A) and major cell types based on manual annotation in Figure 1F (B). **(C)** Dot plot of integrated *Aedes aegypti* Mosquito Cell Atlas annotations and gene markers. Color scale indicates mean normalized expression of gene within cell type, size of dot indicates percent of cells expressing gene within the group. See Table S1 for gene IDs and annotation thresholds.

**Figure S3.**
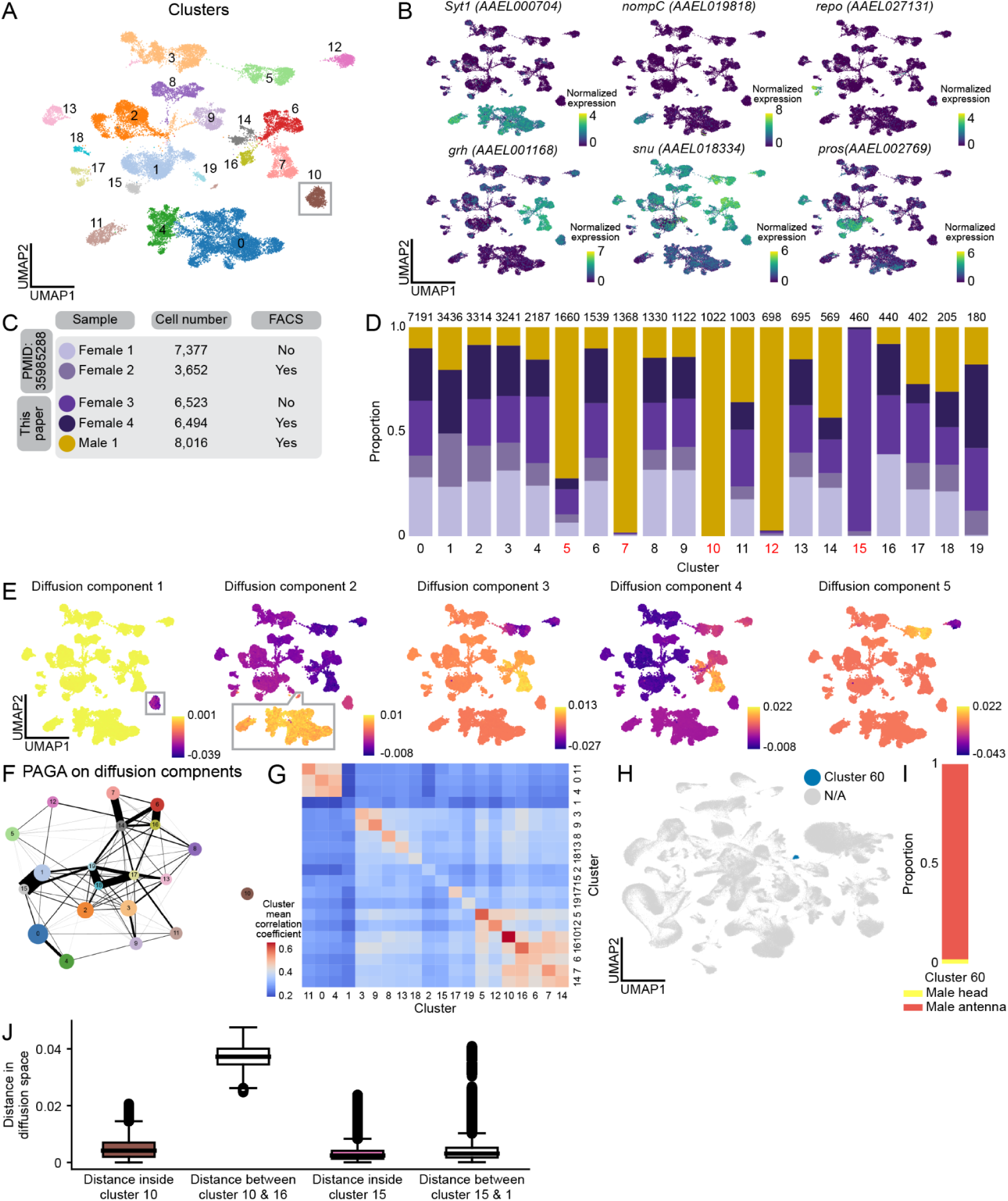
Identification of male-specific *ppk317* cell type in *Aedes aegypti* antenna, related to Figure 3. **(A)** UMAP of antenna nuclei clustered and numbered using the Leiden algorithm (resolution = 0.1). Cluster 10 (male-specific, *ppk317*-expressing cells) highlighted in grey. **(B)** Normalized expression of *Syt1, nompC, repo (AAEL027131), grh (AAEL001168), snu (AAEL018334),* and *pros (AAEL002769)*. Normalized expression is *ln([(raw count/total cell counts)*median total counts across cells]+1)*. **(C)** Number of cells in each sample (female = 4, male = 1), data source, and if each sample underwent fluorescence-activated cell sorting (FACS). **(D)** Stacked bar plot illustrating proportion of each sample within each cluster. Annotated information: cluster numbers (below bar plot), clusters for which over 70% originate from a single sample (red), number of cells in each cluster (above bar plot). **(E)** UMAPs of diffusion components 1 through 5. Diffusion component 1 (first panel) maps to cluster 10 (*ppk317*-expressing cells, highlighted in grey), suggesting a robust biological feature. Diffusion component 2 (second panel) maps to neurons (highlighted in grey). **(F)** Partition-based graph abstraction (PAGA) calculated on diffusion components in (E). All edges illustrated, no edge threshold set. **(G)** Correlation matrix heatmap, depicting pairwise correlation of gene expression matrices between each cluster (mean Pearson correlation coefficient). Diagonal values (cluster to itself) represents intra-cluster correlation values, which vary based on features such as cell number and transcriptome heterogeneity. **(H)** UMAP of integrated *Aedes aegypti* Mosquito Cell Atlas data with cluster 60 colored in blue (Louvain algorithm, resolution = 0.1). *ppk317*-expressing antennal cells belong to cluster 60, see Figure 3E. **(I)** Stacked bar plot indicating tissue origin of cells from cluster 60. Cluster 60 come from the male antenna and male head sample. **(J)** Pairwise Euclidean distances on diffusion embeddings from (E) to approximate phenotypic distance between and within clusters. Boxes represent first quartile to third quartile, and middle line represents median. Whiskers represent the 1.5x interquartile range of the data, with points outside this range as outliers.

**Figure S4.**
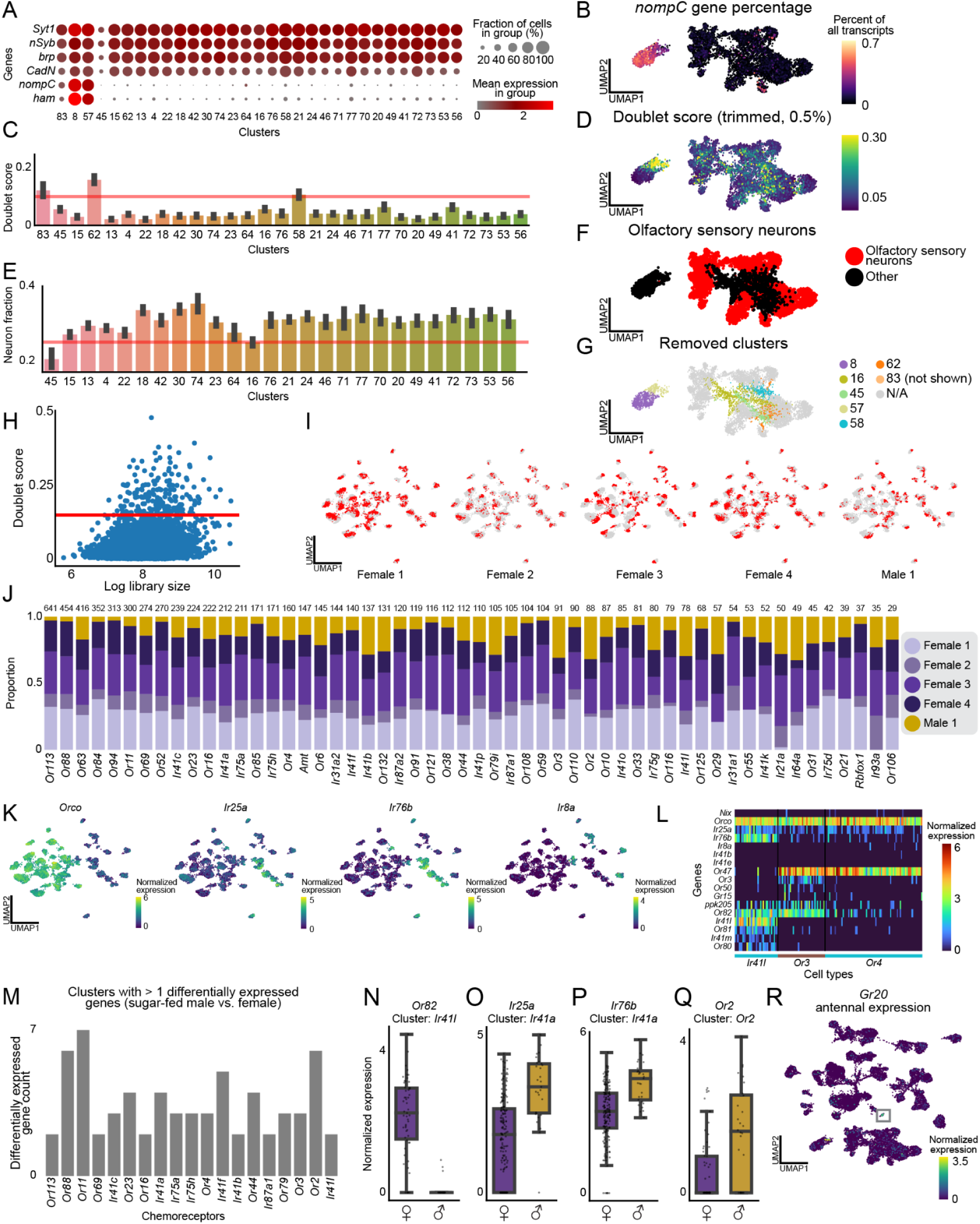
Antenna chemosensory cell type expressing *Orco* and *Ir25a* is sexually dimorphic for *Or82,* related to Figure 4. **(A)** Dot plot illustrating mean normalized expression of neuronal genes set: *Syt1, nSyb, brp (AAEL018153), CadN, nompC,* and *ham (AAEL017229)*. Size of dot indicated the percent of cells in each group, color indicated mean normalized expression. Normalized expression is *ln([(raw count/total cell counts)*median total counts across cells]+1)*. **(B)** Fraction of total transcripts per cell of *nompC* in antennal neuronal nuclei. (B,D,F,G) use the same UMAP coordinates as neurons from Figure 4A. UMAPs cropped for space, cluster 83 not shown. **(C)** Mean doublet score across cells, with error bars indicating 95% confidence interval calculated from bootstrapping. Generated through scrublet^189^. Clusters with an average score above 0.15 were removed from further analysis (red line). **(D)** UMAP depicting doublet score. **(E)** Average percentage of neuronal genes in (A), with error bars indicating 95% confidence interval calculated from bootstrapping. Clusters with an average score below 0.25 were removed from further analysis (red line). **(F)** UMAP demonstrating which clusters (Leiden, resolution = 5) were kept for downstream analysis (red) or removed based on filtering parameters (black). **(G)** UMAP of neurons from antenna samples, demonstrating clusters removed from downstream analysis. **(H)** Log(library size) versus calculated doublet score for neurons filtered in (A-G). Cells with a score above 0.15 were removed from further analysis (red line). **(I)** UMAPs of antenna olfactory sensory neurons (filtered *nompC-*negative sensory neuron population), colored by sample. **(J)** Stacked bar plot illustrating proportion of each sample within each cluster in annotated *nompC*-negative sensory neuron population. Annotated information: cluster numbers (below bar plot), number of cells in each cluster (above bar plot). No clusters has more than 70% of cells from a single sample. **(K)** Normalized gene expression of olfactory co-receptor genes: *Orco, Ir25a, Ir76b,* and *Ir8a*. **(L)** Heatmap of *Ir41l*, *Or3*, and *Or4* cells from female samples 3 and 4 (see Figure S3C). Selected genes are indicated in rows, cells in columns, with cell type annotations below. Heatmap colors represent normalized expression. **(M)** Bar plot showing olfactory sensory neuron cell types with at least 2 differentially expressed genes (DEGs) between male and female cells. Significant genes had |log fold change| >1, false discovery rate <0.05, determined by MAST on normalized expression. For more information on DEGs, see Table S2. **(N-Q)** Distribution of expression for differentially expressed olfactory receptor genes across male and female cells in a particular cluster. *Or82* expression in cluster *Ir41l* (N), *Ir25a* in cluster *Ir41a* (O), *Ir76b* in cluster *Ir41a* (P), *Or2* expression in cluster *Or2* (Q). Boxes represent first quartile to third quartile, and middle line represents median. Whiskers represent the 1.5x interquartile range of the data, with points outside this range as outliers. **(R)** *Gr20* normalized expression. *Gr20* positive antennal cluster highlighted in grey.

**Figure S5.**
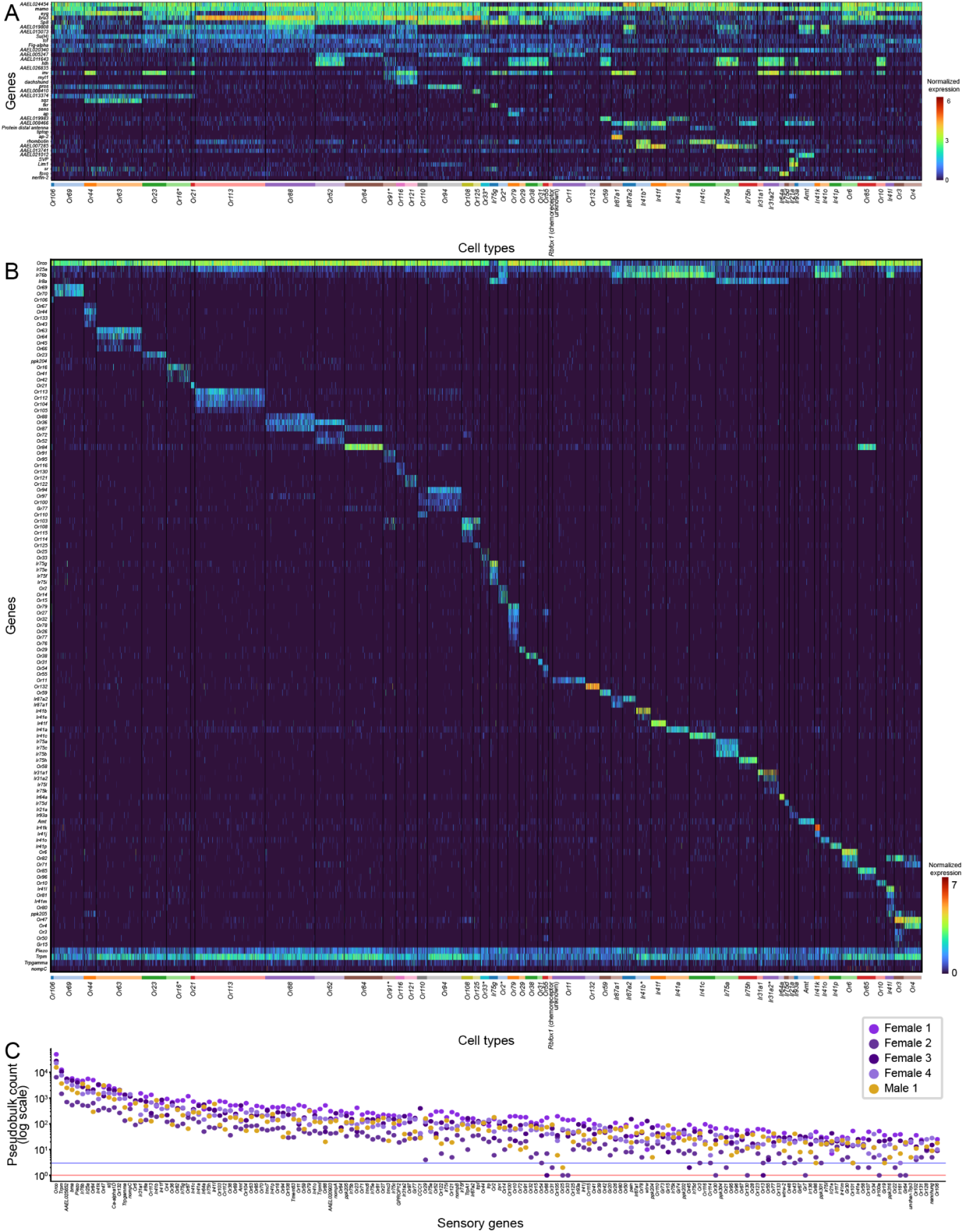
Antenna chemosensory neuron chemoreceptor and putative transcription factor factor expression profiles, related to Figure 4. **(A-B)** Heatmap of all annotated antenna olfactory sensory neuron cell types for selected putative transcription factors (A) and selected sensory genes (B). Selected genes are indicated in rows, cells in columns, with cell type annotations below. Cells are grouped by annotation in Figure 4C and indicated below heatmap. Asterisks (*) on cell type annotation indicates that a group may represent a co-clustering of cells with multiple unique sensory gene expression patterns, as depicted in Figure 4C-4D. Heatmap colors represent normalized expression. Normalized expression is *ln([(raw count/total cell counts)*median total counts across cells]+1)*. See Table S1 for gene IDs and annotation thresholds. **(C)** Scatterplot of summed raw counts of sensory genes within all olfactory sensory neurons within each antenna sample. Red line indicates count of 1, blue line is a count of 5. 158 out of 403 putative sensory genes (Table S1) with the highest counts shown for space (full plot available in Zenodo Supplemental Data).

**Figure S6.**
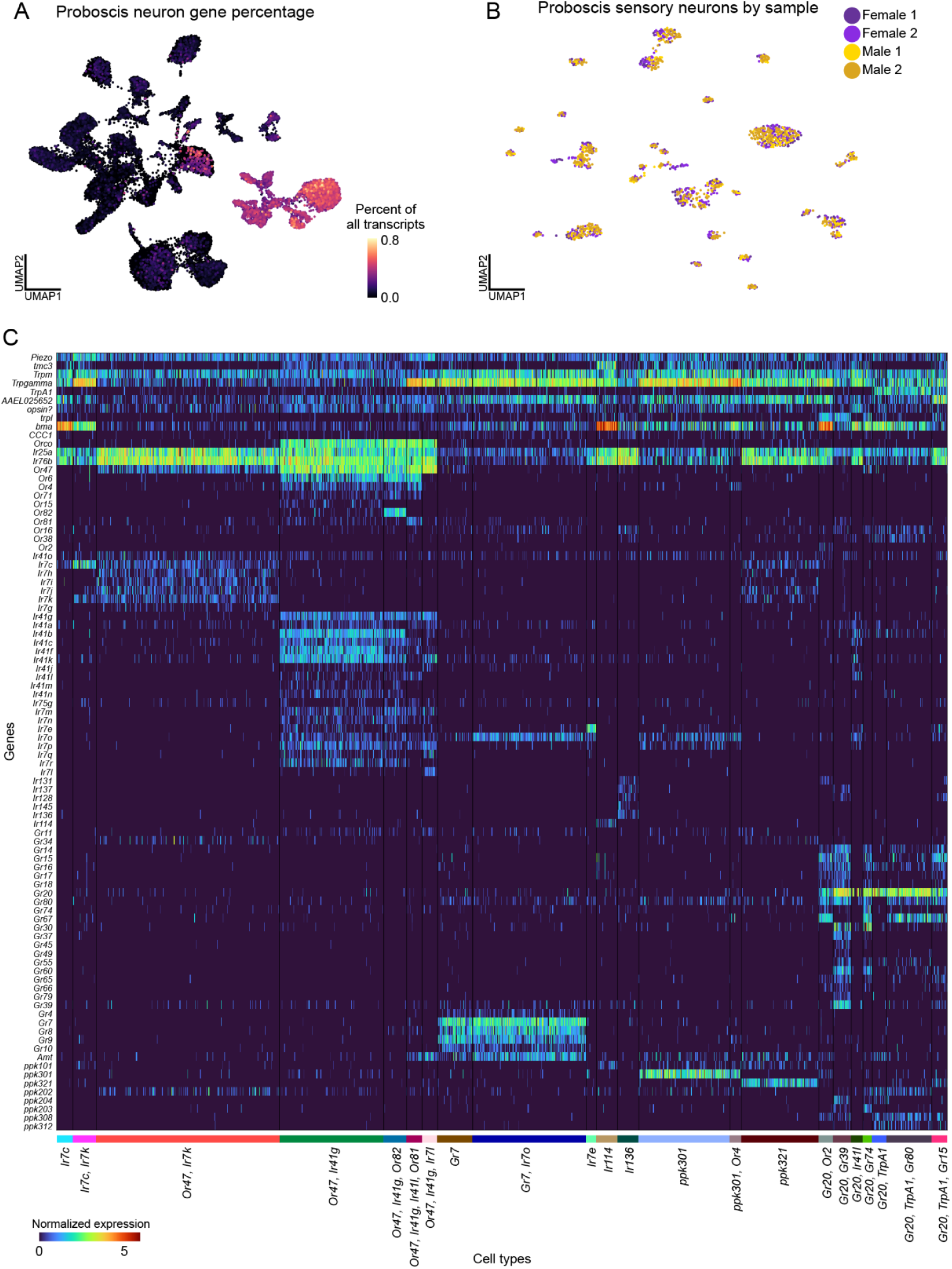
Proboscis sensory gene analysis, related to Figures 4 and 5. **(A)** UMAP of proboscis cells illustrating fraction of total transcripts per cell of neuronal genes set: *Syt1*, *brp*, *nSyb*, *CadN*. *nompC*-negative cells highlighted (grey box). For *nompC* gene percentage, see Data S4. **(B)** UMAP of reclustered proboscis *nompC-*negative sensory cells colored by sample (female = 2, male = 2). **(C)** Heatmap of cells from all annotated clusters. Sensory genes are indicated in rows and cells indicated in columns. Selected genes are indicated in rows, cells in columns, with cell type annotations below. Heatmap colors represent normalized expression. Normalized expression is *ln([(raw count/total cell counts)*median total counts across cells]+1)*.

**Figure S7.**
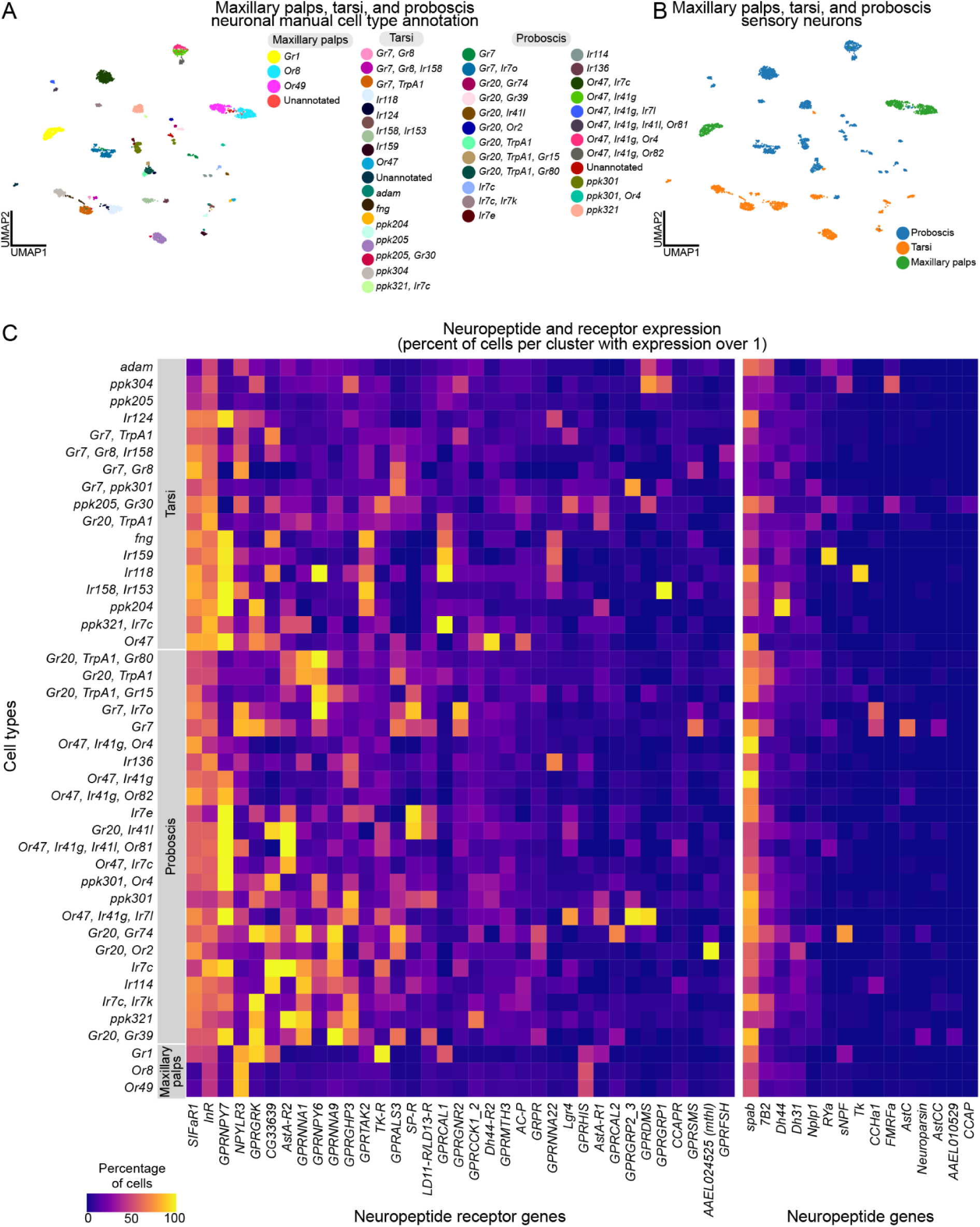
Neuropeptide receptor and synthesis gene analysis, related to Figures 4 and 5. **(A-B)** UMAP of *nompC-*negative sensory neurons from maxillary palps, tarsi and proboscis samples, colored by manual cell type annotation as listed in legend at the right of the figure panel **(A)** and original tissue **(B)**. **(C)** Heatmap of expression of neuropeptide receptor genes (left) and neuropeptide synthesis genes (right) within annotated *nompC-*negative sensory neurons in the maxillary palps, tarsi and proboscis. Color scale indicates percentage of cells expressing a gene above threshold (normalized expression value of 1). Sensory genes are indicated in columns and annotated cell types indicated in rows. Genes were included if they were expressed above threshold in over 20% of cells in at least one cell type. Genes filtered from lists in Table S1. Normalized expression is *ln([(raw count/total cell counts)*median total counts across cells]+1)*.

**Figure S8.**
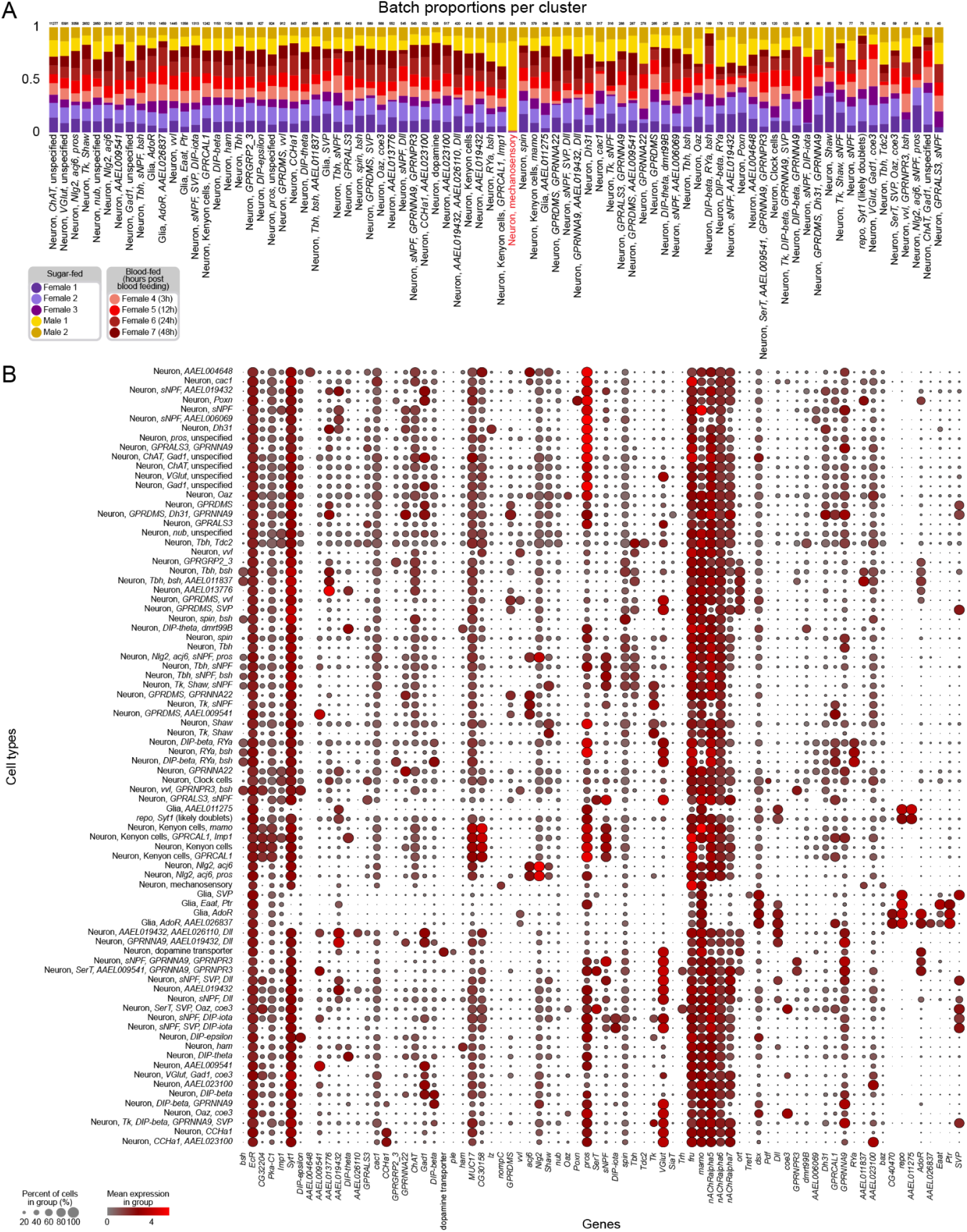
Brain Cell types, related to Figures 6 and 7. **(A)** Stacked bar plot illustrating proportion of each sample within each cluster for brain nuclei cell types. Annotated information: cluster numbers (below bar plot), clusters for which over 70% originate from a single sample (red), number of cells in each cluster (above bar plot). **(B)** Dot plot illustrating mean normalized expression of gene markers. Color scale indicates mean normalized expression of gene within cell type, size of dot indicates percent of cells expressing gene within the group. See Table S1 for gene IDs and annotation thresholds.

**Figure S9.**
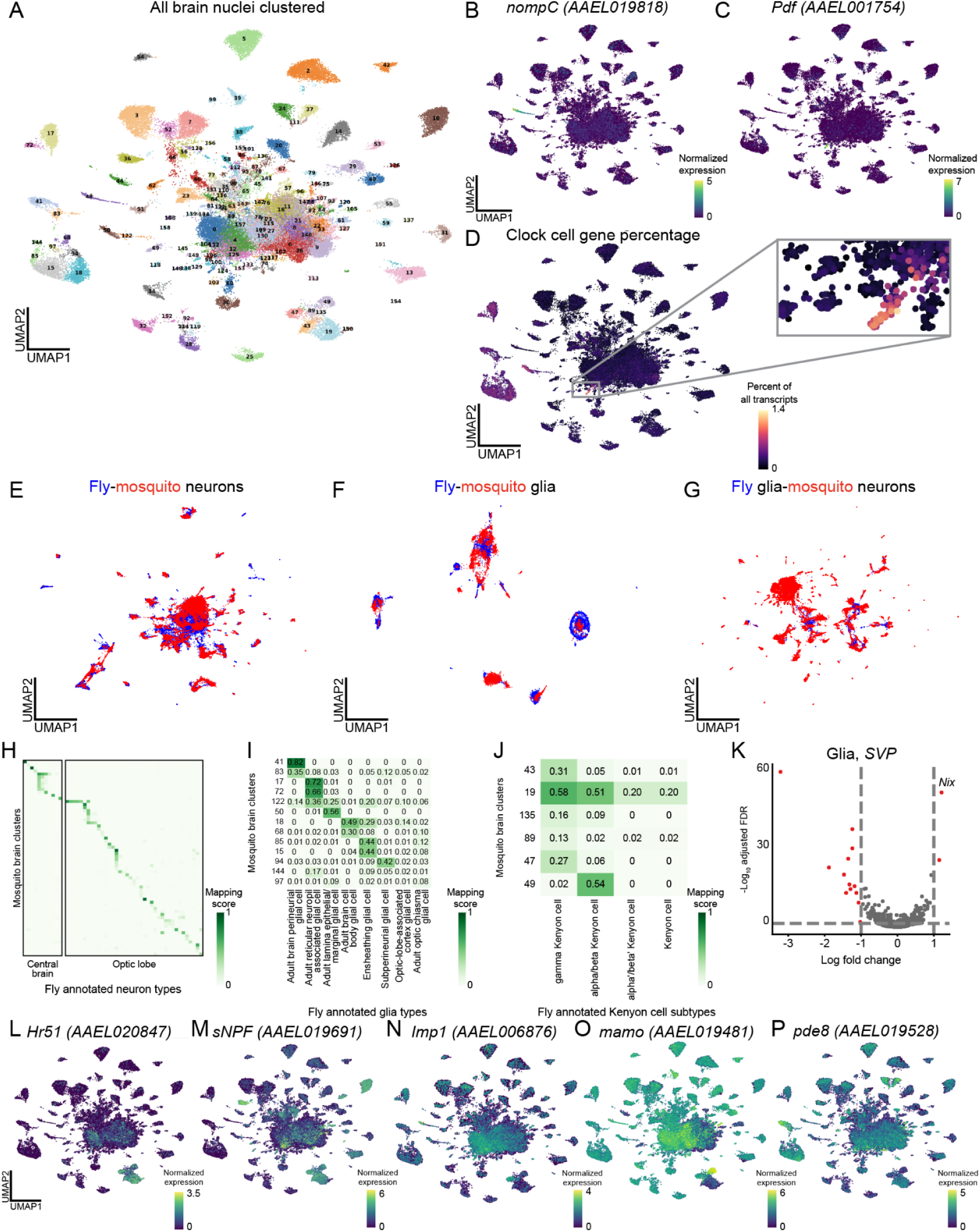
SAMap analysis and identification of clock cells and kenyon cells, related to Figure 6. **(A)** UMAP of brain nuclei clustered using the Leiden algorithm (resolution = 5). **(B-C)** Normalized expression of *nompC* (B) and *Pdf (AAEL001754)* (C). Normalized expression is *ln([(raw count/total cell counts)*median total counts across cells]+1)*. **(D)** Fraction of total transcripts per cell of 10 putative clock cell gene markers (Table S1). Cluster with high expression highlighted in grey box, enlarged in inset. **(E-G)** UMAP of manifold integration of snRNA-seq data from *Aedes aegypti* mosquito brain and published *Drosophila melanogaster* fly head^18^. Plots show integration of fly neurons with mosquito neurons (E), fly glia with mosquito glia (F), and, as a control, fly glia with mosquito neuron (G). Alignment scores are 0.64, 0.64 and 0.47, respectively. **(H-J)** Correlation matrices of mapping scores between *Drosophila melanogaster* head annotations and *Aedes aegypti* of clusters (Leiden, resolution = 5) for neuronal cell types (H), glial cell types (I), and Kenyon cell subtypes (J). For all numerical values, see Table S3. **(K)** Volcano plot of differentially expressed genes between male and female cells in the *SVP* glia *(AAEL002765*) cluster. Significant genes (red) had |log fold change| >1, false discovery rate <0.05, determined by MAST on normalized expression (Table S2). Male biased genes on right, indicated by *Nix* (*AAEL022912*). **(L-P)** Normalized gene expression of *Hr51 (AAEL020847)* (L), *sNPF (AAEL019691)* (M), *Imp1 (AAEL006876)* (N), *mamo (AAEL019481)* (O)*, Pde8 (AAEL019528)* (P).

**Figure S10.**
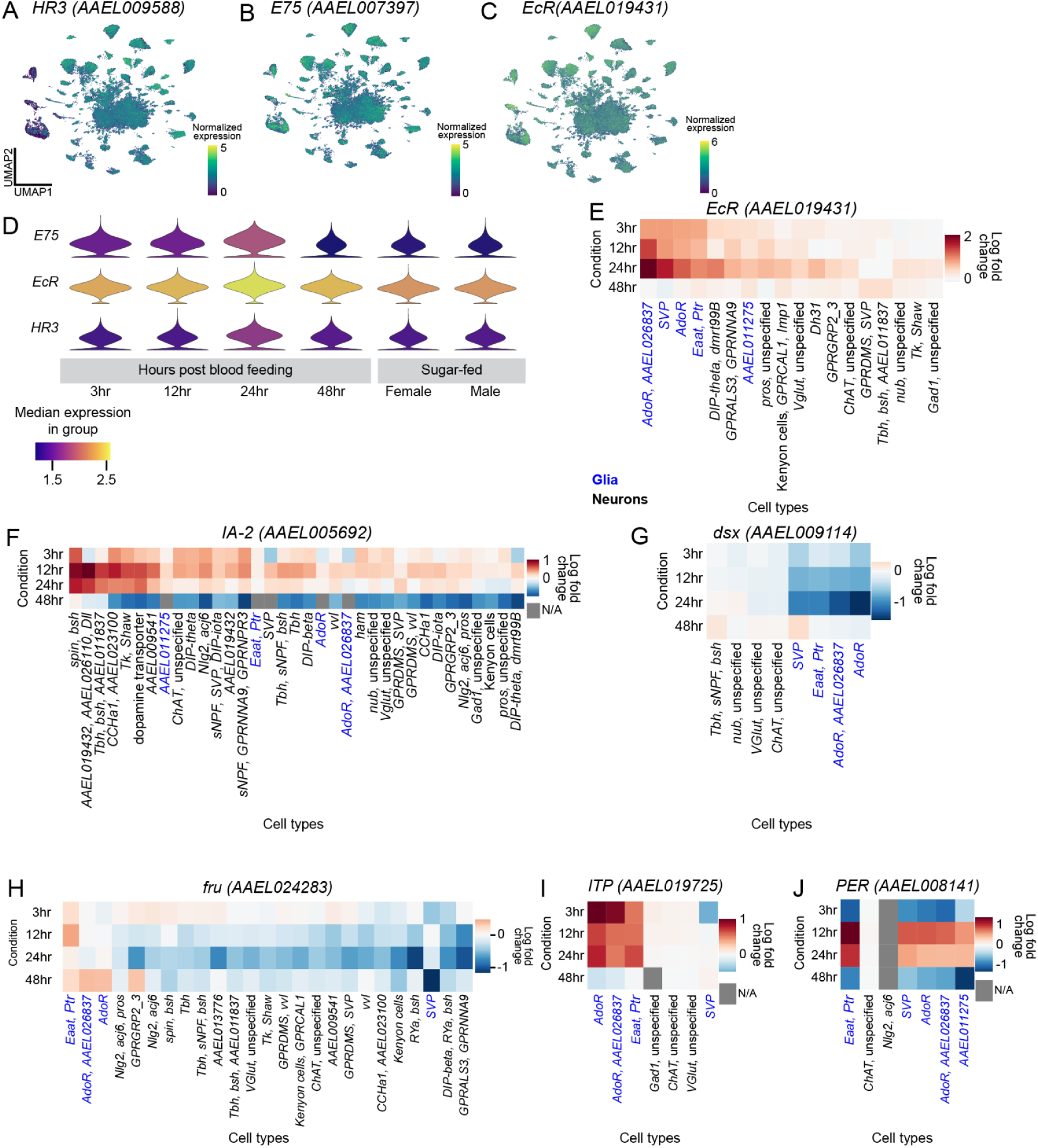
Blood feeding changes in brain, related to Figure 7. **(A-C)** Normalized gene expression UMAP of *E75* (A), *EcR* (*AAEL019431*) (B) and *HR3* (C) in all brain nuclei. Normalized expression is *ln([(raw count/total cell counts)*median total counts across cells]+1)*. **(D)** Violin plot of gene expression of *E75*, *EcR* and *HR3* across all brain nuclei in each timepoint. Length of violin indicates complete range of the data, with thickness of the violin representing number of cells at each value. **(E-J)** Heatmaps of log fold change of gene expression, grouped by annotated cell type, between corresponding cells collected in each blood-feeding timepoint compared to sugar-fed female brain. Genes shown: *EcR* (E), *IA-2 (AAEL005692)* (F), *dsx* (*AAEL009114*), (G), *fru* (*AAEL024283*) (H), *ITP* (*AAEL019725*) (I), and *PER* (*AAEL008141*) (J). Log fold change is determined by MAST on normalized expression. Cell types are sorted by the total log fold change across all timepoints and colored as glia (blue) or neurons (black). Cell types included have over 10 cells in each timepoint, and at least one timepoint where change from sugar-fed condition had a false discovery rate <0.05. Grey boxes indicate log fold change data is not available, due to zero expression within cell type at the specified timepoint (or the sugar-fed condition for *PER* in *Nlg2, acj6* cells).

## Supplemental Data and table titles

**Data S1.** Example sample quality-control filtering, and gene expression patterns in the testes and antimicrobial peptide-expressing tissues, related to Figures 1 and 2.

**Data S2**. Individual tissue annotations of mated, sugar-fed tissues, related to Figures 1-6.

**Data S3.** RNA *in situ* hybridization controls and probe information, related to Figures 2-4.

**Data S4.** Filtering and characterization of *nompC* (*AAEL019818*)-negative sensory neurons, related to Figure 5.

**Data S5.** *Aedes aegypti* Mosquito Cell Atlas Consortium Members and Funding

**Table S1.** Sample, processing, annotation, and gene information for the *Aedes aegypti* Mosquito Cell Atlas (related to Figure 1).

**Table S2.** Differential gene expression for sensory appendage, brain, fat tissue and gut cell types, across sex or bloodfeeding (related to Figures 4-7).

**Table S3.** Cross-species manifold integration mapping of *Aedes aegypti* brain cells to *Drosophila melanogaster* head cell types (related to Figure 6).

