## Supplemental Data S1 for "A single-nucleus transcriptomic atlas of the adult *Aedes aegypti* mosquito"

Example sample quality-control filtering, and gene expression patterns in the testes and antimicrobial peptide-expressing tissues, related to Figures 1 and 2.

Index:

**(Data S1.1)** Illustration of individual sample processing and cluster quality-control filtering, shown on female salivary gland, related to Figures 1 and 2.

**(Data S1.2)** Localization of spermatid RNA transcripts in mated, sugar-fed testes data, related to Figure 2.

**(Data S1.3)** Antimicrobial genes in fat tissue, related to Figure 2.

**(Data S1.4)** Sexually differential expressed genes in gut enterocytes and abdominal pelt fat tissue, related to Figures 1 and 2.

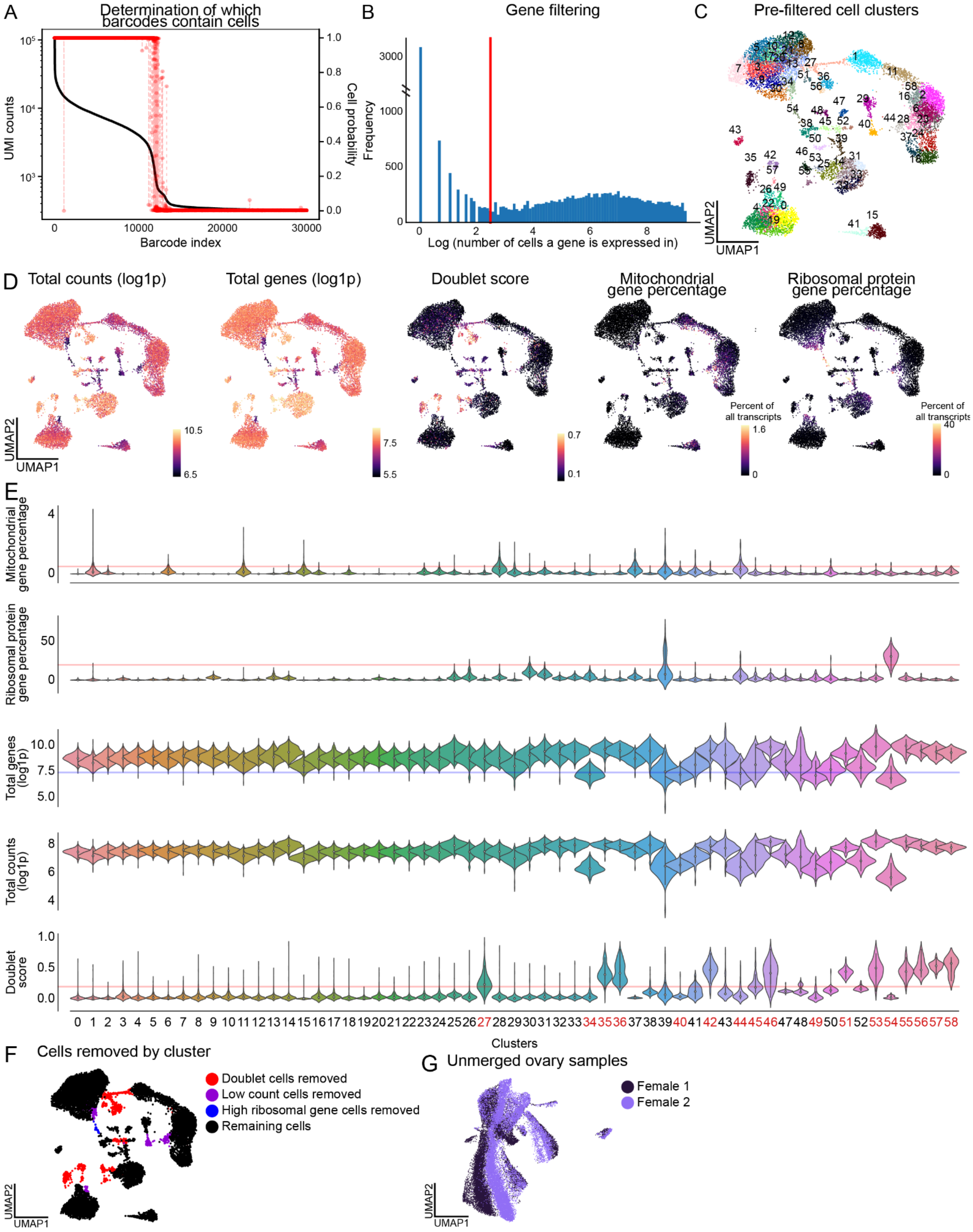
A
Determination of which barcodes contain cells
UMI counts
Cell probability
Barcode index
B
Gene filtering
Frequency
Log (number of cells a gene is expressed in)
C
Pre-filtered cell clusters
UMAP2
UMAP1
D
Total counts (log1p)
Total genes (log1p)
Doublet score
Mitochondrial gene percentage
Ribosomal protein gene percentage
Percent of all transcripts
E
Mitochondrial gene percentage
Ribosomal protein gene percentage
Total genes (log1p)
Total counts (log1p)
Doublet score
Clusters
F
Cells removed by cluster
Doublet cells removed
Low count cells removed
High ribosomal gene cells removed
Remaining cells
G
Unmerged ovary samples
Female 1
Female 2

**Data S1.1. Illustration of individual sample processing and cluster quality-control filtering, shown on female salivary glands, related to Figures 1 and 2.**
**(A)** Determination of individual cell barcodes that contain cells by CellBender[222]. Unique molecular identifier (UMI) counts (black, left axis), probability that a barcode index is a cell (red, right axis). Low probability cells were removed by the CellBender.
**(B)** Histogram of number of cells that an individual gene is expressed in (log scale). Genes expressed in fewer than 13 cells were removed from a given sample (red line).
**(C)** UMAP of mated, sugar-fed salivary gland cells clustered using the Leiden algorithm (resolution = 5) for quality control and cell filtering.
**(D-E)** UMAPs **(D)** and violin plots **(E)** depicting total counts, total genes, doublet score (generated through scrublet[197], mitochondrial gene percentage, ribosomal gene percentages (log1p, indicates a pseudocount of 1 was used for log-scaling) for each cell and across each cluster. Lines in (E) indicate thresholds by which clusters with an average value above (red) or below (blue) were removed from further analysis. Removed clusters indicated in red on the bottom of (E).
**(F)** UMAP colored by cluster that were kept for further analysis (black) or removed due to quality control metrics depicted in (D-E).
**(G)** UMAP of mated, sugar-fed ovary samples female 1 (dark purple) and female 2 (light purple) prior to merging.

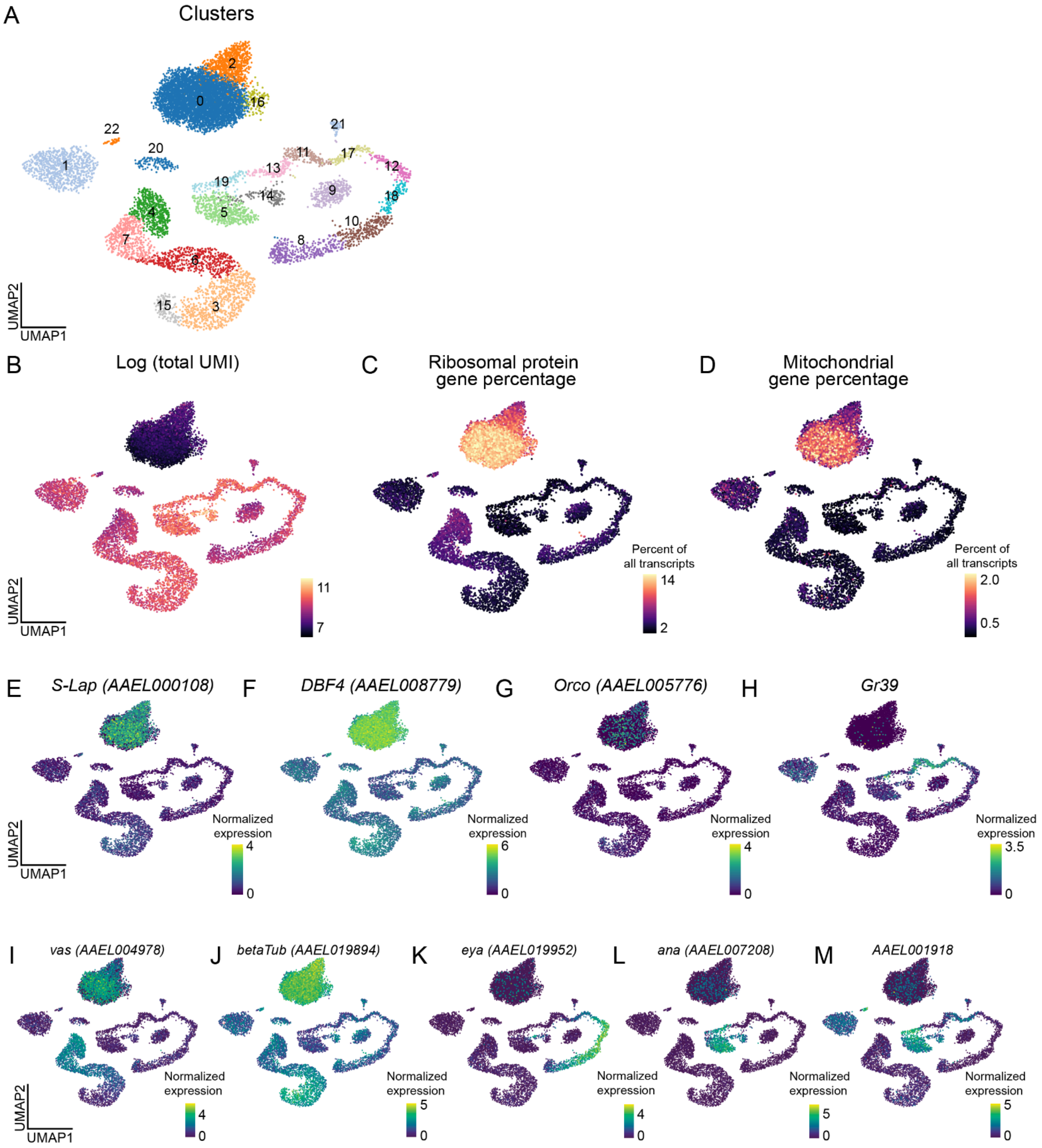
A
Clusters
UMAP2
UMAP1
B
Log (total UMI)
11
7
C
Ribosomal protein gene percentage
Percent of all transcripts
14
2
D
Mitochondrial gene percentage
Percent of all transcripts
2.0
0.5
E
S-Lap (AAEL000108)
Normalized expression
4
0
F
DBF4 (AAEL008779)
Normalized expression
6
0
G
Orco (AAEL005776)
Normalized expression
4
0
H
Gr39
Normalized expression
3.5
0
I
vas (AAEL004978)
Normalized expression
4
0
J
betaTub (AAEL019894)
Normalized expression
5
0
K
eya (AAEL019952)
Normalized expression
4
0
L
ana (AAEL007208)
Normalized expression
5
0
M
AAEL001918
Normalized expression
5
0

**Data S1.2. Localization of spermatid RNA transcripts in mated, sugar-fed testes data, related to Figure 2.**
**(A)** UMAP of all testes cells after quality control and filtering (Louvain algorithm, resolution = 1).
**(B)** UMAP of testes cells colored by log total UMI for each cell.
**(C-D)** UMAP of testes sample colored by percent of all transcripts in each cell for expression of ribosomal protein genes (C), expression of mitochondrial genes (D). Each end of color bars trimmed 0.5% for visibility.
**(E-H)** UMAPs of testes normalized expression of *S-Lap* (*AAEL000108*) (E), *DBF4* (*AAEL008779*) (F), *Orco* (*AAEL005776*) (G), and *Gr39* (H). Normalized expression is ln([(raw count/total cell counts)x median total counts across cells]+1).
**(I-M)** UMAP of normalized gene expression of a subset of genes used to annotate testes data that were used in Figures 2C-2G. Genes include include *vas* (*AAEL004978*) (I), *betaTub* (*AAEL019894*) (J), *eya (AAEL019952*) (K), *ana (AAEL007208)* (L), and *AAEL001918* (M). Note normalized signal of genes expressed in spermatids appears high relative to other cell types in normalized expression due to their overall low transcript count (Data S1).

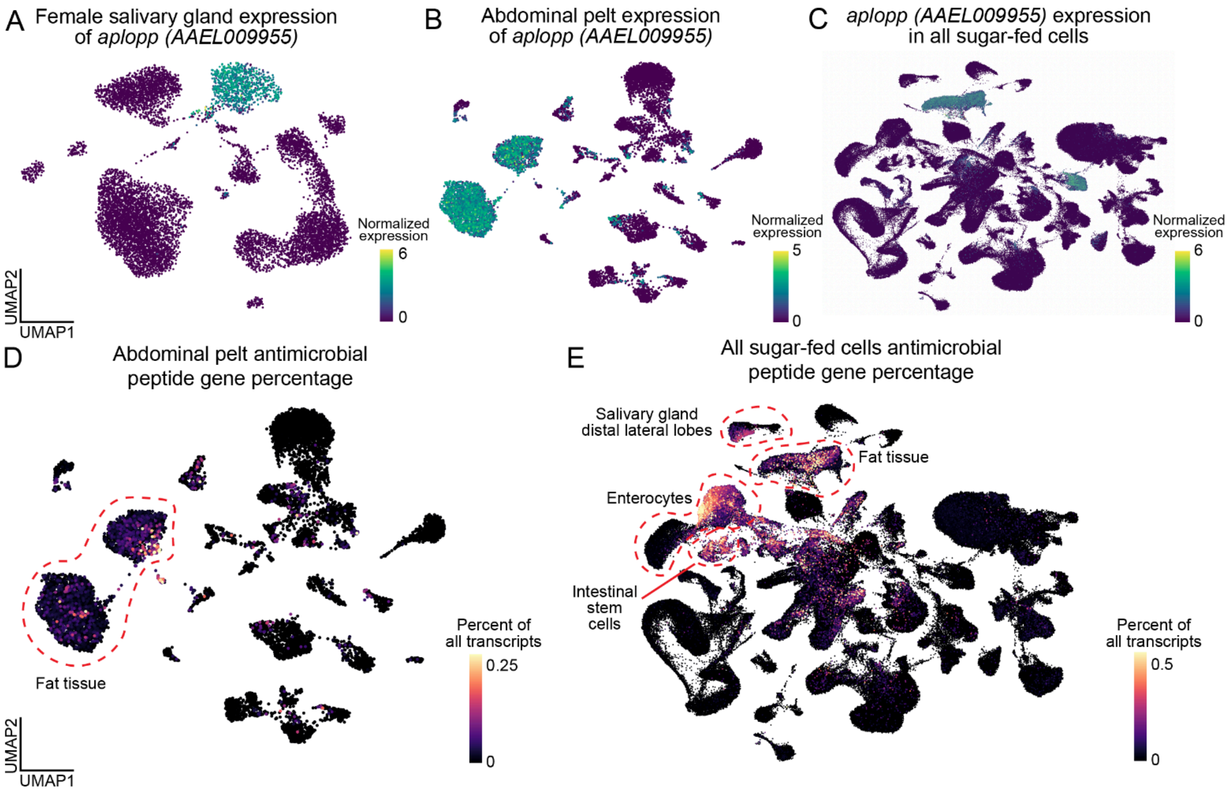

**Data S1.3. Antimicrobial genes in fat tissue, related to Figure 2.**
**(A-C)** UMAPs of normalized expression of marker gene for fat tissue *aplopp* (*AAEL009955*) in the salivary gland (A), abdominal pelt (B), and all mated, sugar-fed nuclei (C). Normalized expression is ln([(raw count/total cell counts)x median total counts across cells]+1).
**(D-E)** UMAP, colored by expression of antimicrobial peptides gene set (Table S1) for abdominal pelt (D) and all sugar-fed nuclei (E). Color represents fraction of total transcripts in each cell (each end of color bar trimmed 0.1% for visibility). Relevant cell types labeled.

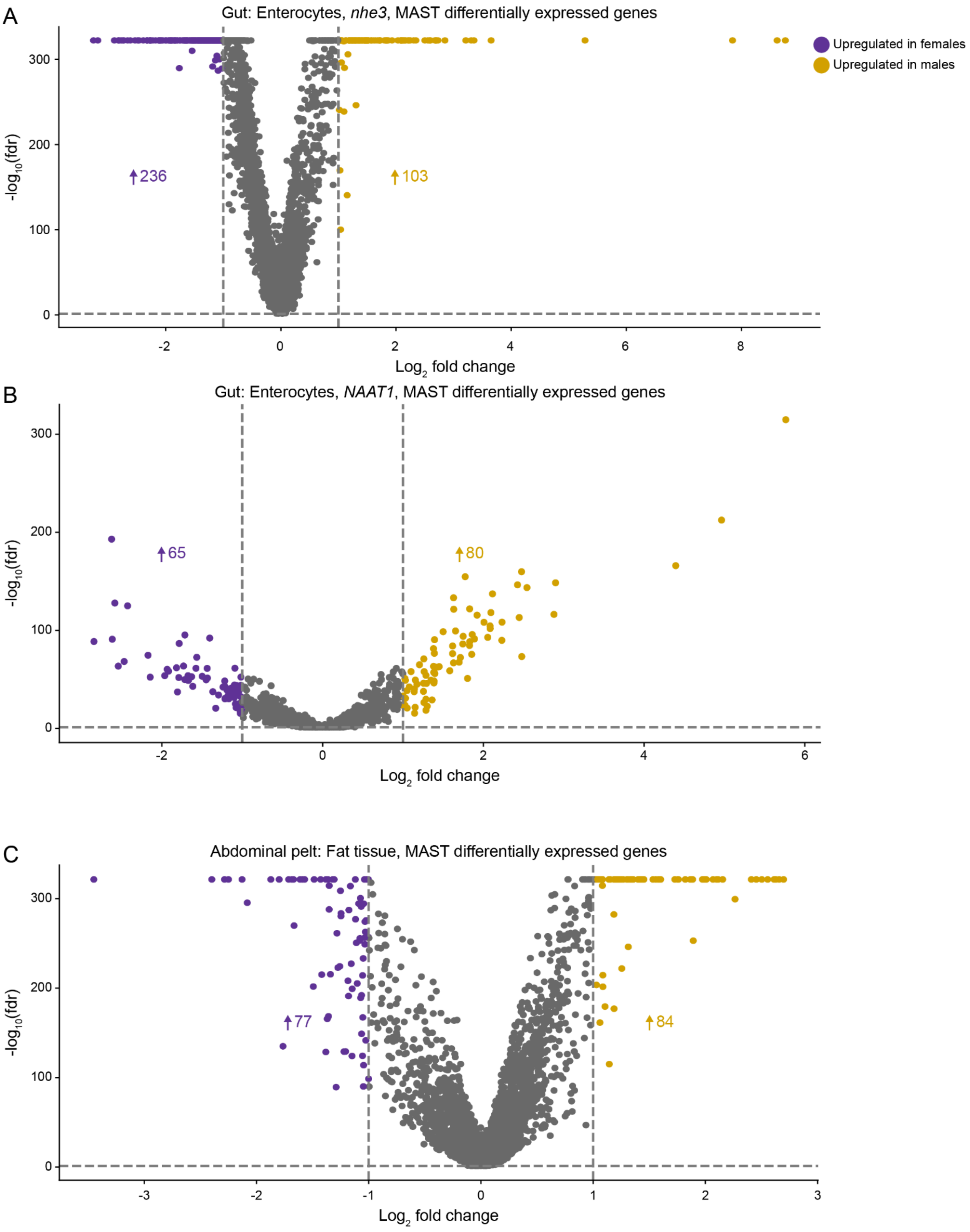
A
Gut: Enterocytes, nhe3, MAST differentially expressed genes
Upregulated in females
Upregulated in males
236
103
-log10(fdr)
0
100
200
300
-2
0
2
4
6
8
Log2 fold change
B
Gut: Enterocytes, NAAT1, MAST differentially expressed genes
65
80
-log10(fdr)
0
100
200
300
-2
0
2
4
6
Log2 fold change
C
Abdominal pelt: Fat tissue, MAST differentially expressed genes
77
84
-log10(fdr)
0
100
200
300
-3
-2
-1
0
1
2
3
Log2 fold change

**Data S1.4. Sexually differential expressed genes in gut enterocytes and abdominal pelt fat tissue, related to Figures 1 and 2.**

**(A-C)** Volcano plots of sexually differential expressed genes (DEGs) between males and females in the gut enterocytes, "*nhe3*" cell type (A) gut enterocytes, "*NAAT1*" cell type (B), and abdominal pelt fat tissue (C). All significantly upregulated genes in females (indicated in purple) and all significantly upregulated genes in males (indicated in yellow) are |log fold change| > 1 and false discovery rate < 0.05, determined by MAST on normalized expression. Number of upregulated genes in females (purple) and upregulated genes in males (yellow) are indicated by the up arrows. For more information on DEGs, see Table S2.
