## Supplemental Data S2 for "A single-nucleus transcriptomic atlas of the adult *Aedes aegypti* mosquito"

Individual tissue annotations of mated, sugar-fed tissues, related to Figures 1-6.

Index:

**(Data S2.1)** Manual cell type annotation of **female and male head** data. Includes differentially expressed genes (DEGs) between male and female samples, and differences in cell type abundance, related to Figure 1.
**(Data S2.2)** Manual cell type annotation of **female and male thorax** data. Includes differentially expressed genes (DEGs) between male and female samples, and differences in cell type abundance, related to Figure 1.
**(Data S2.3)** Manual cell type annotation of **female and male abdomen** data. Includes differentially expressed genes (DEGs) between male and female samples, and differences in cell type abundance, related to Figure 1.
**(Data S2.4)** Manual cell type annotation of **female and male maxillary palps** data. Includes differentially expressed genes (DEGs) between male and female samples, and differences in cell type abundance, related to Figure 1.
**(Data S2.5)** Manual cell type annotation of **female and male thoracic ganglia** data. Includes differentially expressed genes (DEGs) between male and female samples, and differences in cell type abundance, related to Figure 1.
**(Data S2.6)** Manual cell type annotation of **female and male gut** data. Includes differentially expressed genes (DEGs) between male and female samples, and differences in cell type abundance, related to Figure 1.
**(Data S2.7)** Manual cell type annotation of **female and male abdominal pelt** data. Includes differentially expressed genes (DEGs) between male and female samples, and differences in cell type abundance, related to Figure 1.
**(Data S2.8)** Manual cell type annotation of **female and male malpighian tubules** data. Includes differentially expressed genes (DEGs) between male and female samples, and differences in cell type abundance, related to Figure 1.
**(Data S2.9)** Manual cell type annotation of **female and male wings** data. Includes differentially expressed genes (DEGs) between male and female samples, and differences in cell type abundance, related to Figure 1.
**(Data S2.10)** Manual cell type annotation of **female stylet, female abdominal tip, female ovary, and male reproductive glands**, related to Figure 1.
**(Data S2.11)** Manual cell type annotation of **female and male antennae** data. Includes differentially expressed genes (DEGs) between male and female samples, and differences in cell type abundance, related to Figures 1 and 3.
**(Data S2.12)** Manual cell type annotation of **female and male tarsi** data. Includes differentially expressed genes (DEGs) between male and female samples, and differences in cell type abundance, related to Figures 1, 4, and 5.

**(Data S2.13)** Manual cell type annotation of **female and male proboscis** data. Includes differentially expressed genes (DEGs) between male and female samples, and differences in cell type abundance, related to Figures 1, 4, and 5.

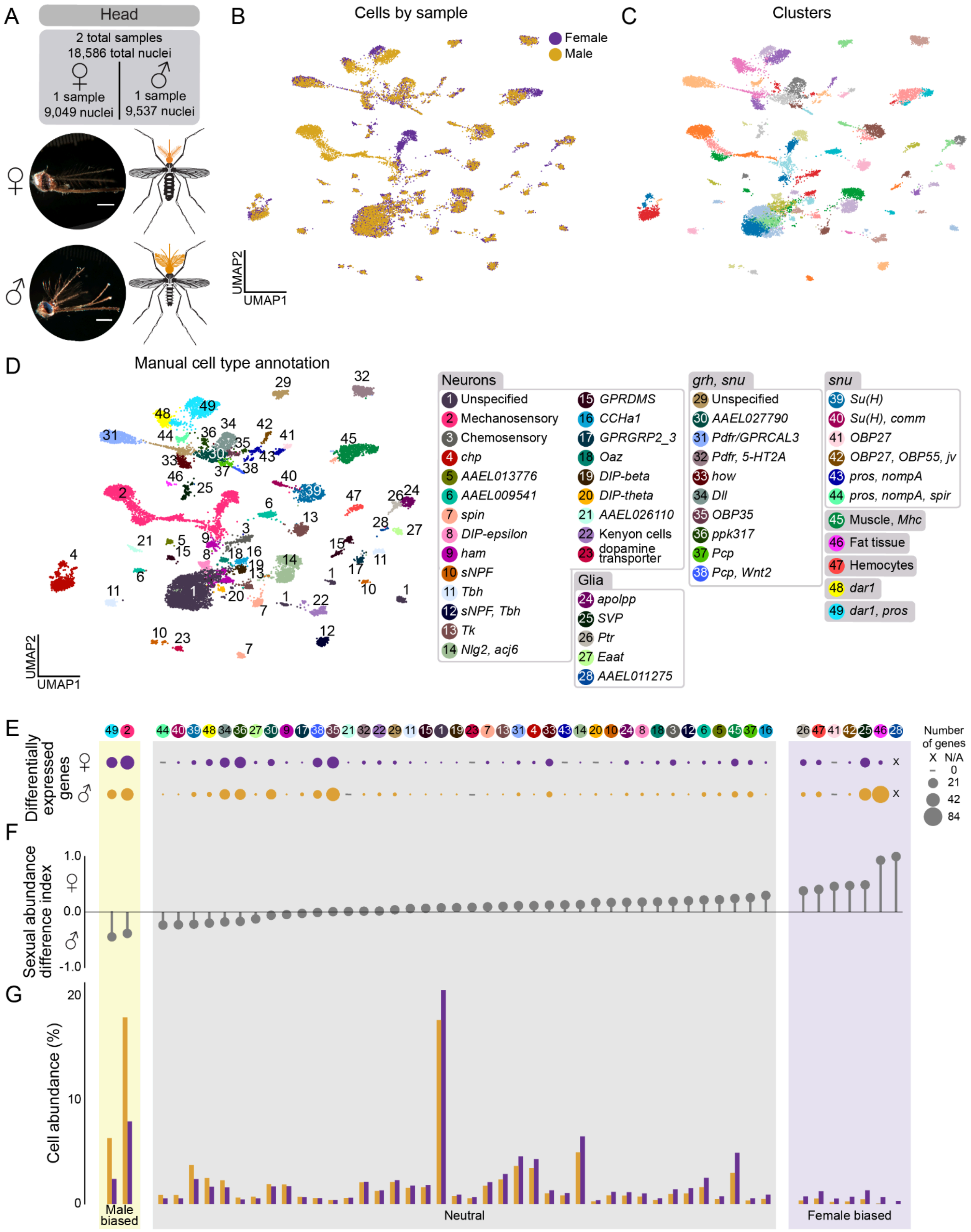
A
Head
2 total samples
18,586 total nuclei
1 sample
9,049 nuclei
1 sample
9,537 nuclei
B
Cells by sample
Female
Male
UMAP2
UMAP1
C
Clusters
D
Manual cell type annotation
UMAP2
UMAP1
Neurons
1 Unspecified
2 Mechanosensory
3 Chemosensory
4 chp
5 AAEL013776
6 AAEL009541
7 spin
8 DIP-epsilon
9 ham
10 sNPF
11 Tbh
12 sNPF, Tbh
13 Tk
14 Nlg2, acj6
15 GPRDMS
16 CCHa1
17 GPRGRP2_3
18 Oaz
19 DIP-beta
20 DIP-theta
21 AAEL026110
22 Kenyon cells
23 dopamine transporter
Glia
24 apolpp
25 SVP
26 Ptr
27 Eaat
28 AAEL011275
grh, snu
29 Unspecified
30 AAEL027790
31 Pdfr/GPRCAL3
32 Pdfr, 5-HT2A
33 how
34 Dll
35 OBP35
36 ppk317
37 Pcp
38 Pcp, Wnt2
snu
39 Su(H)
40 Su(H), comm
41 OBP27
42 OBP27, OBP55, jv
43 pros, nompA
44 pros, nompA, spir
45 Muscle, Mhc
46 Fat tissue
47 Hemocytes
48 dar1
49 dar1, pros
E
Differentially expressed genes
Number of genes
X N/A
0
21
42
84
F
Sexual abundance difference index
1.0
0.0
-1.0
G
Cell abundance (%)
20
10
0
Male biased
Neutral
Female biased

**Data S2.1. Manual cell type annotation of female and male head data. Includes differentially expressed genes (DEGs) between male and female samples, and differences in cell type abundance, related to Figure 1.**
**(A)** Representative photo of mated, sugar-fed female (top) and male (bottom) head dissection with anatomical diagram (in orange). Two samples (10x Genomics libraries) yielded 17,937 nuclei from 60 animals, post-quality control filtering. Scale bar: 500 µm.
**(B)** UMAP of head nuclei, colored by sample (female samples = 1, male samples = 1).
**(C)** UMAP of head cells clustered using the Leiden algorithm (resolution = 5) for annotation.
**(D)** UMAP of nuclei from both head samples, colored and numbered by manual annotation using selected marker genes as listed in legend at the right of the figure panel. Categorization of cell type annotations represented by shaded gray headers. See Table S1 for gene IDs and thresholds.
**(E-G)** Quantification of differences between male and female samples. Numbered and colored circles above (E) correspond to annotations in (D).
**(E)** Dot plot of number of differentially expressed genes between females and males in each cell type. Dots represent relative number of differentially expressed genes upregulated in male or female with a |log fold change| > 1 and false discovery rate < 0.05, determined by MAST on normalized expression and converted to $\log_2$ fold change.
**(F)** Sexual abundance difference index from data in (G). Each cell type was categorized based on the following index: Female biased (purple) if 0.3 < abundance index; Neutral (grey) if -0.3 ≤ abundance index ≤ 0.3; Male biased (yellow) if abundance index < -0.3.
**(G)** Bar plot of relative cell abundance, represented as the percent of each cell type of all nuclei collected from each sex for tissue. Separated by female (purple) and male (yellow).

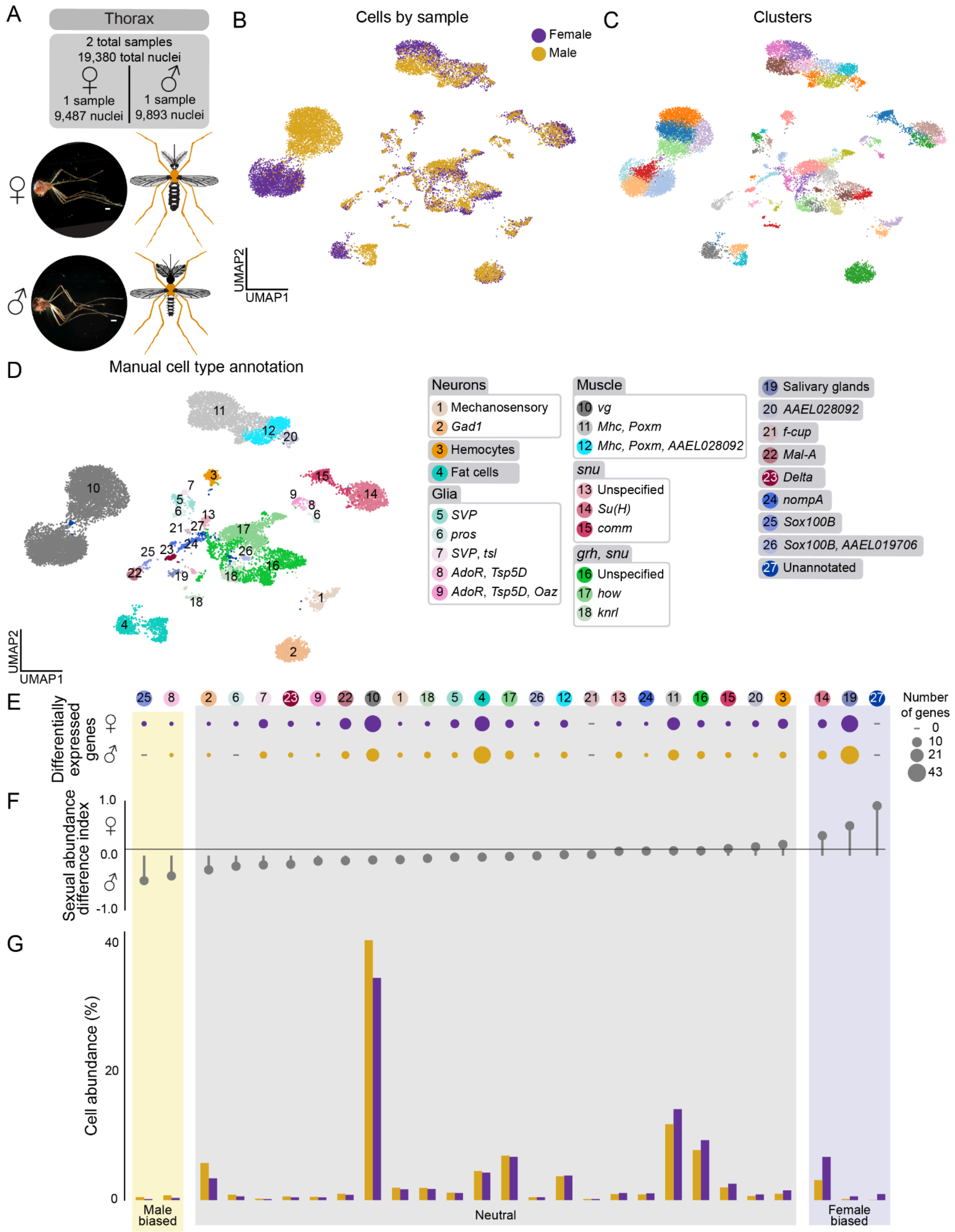
A
Thorax
2 total samples
19,380 total nuclei
1 sample
9,487 nuclei
1 sample
9,893 nuclei
B
Cells by sample
Female
Male
UMAP2
UMAP1
C
Clusters
D
Manual cell type annotation
Neurons
1 Mechanosensory
2 Gad1
3 Hemocytes
4 Fat cells
Glia
5 SVP
6 pros
7 SVP, tsl
8 AdoR, Tsp5D
9 AdoR, Tsp5D, Oaz
Muscle
10 vg
11 Mhc, Poxm
12 Mhc, Poxm, AAEL028092
snu
13 Unspecified
14 Su(H)
15 comm
grh, snu
16 Unspecified
17 how
18 knrl
19 Salivary glands
20 AAEL028092
21 f-cup
22 Mal-A
23 Delta
24 nompA
25 Sox100B
26 Sox100B, AAEL019706
27 Unannotated
E
Differentially expressed genes
Number of genes
0
10
21
43
F
Sexual abundance difference index
1.0
0.0
-1.0
G
Cell abundance (%)
40
20
0
Male biased
Neutral
Female biased

**Data S2.2. Manual cell type annotation of female and male thorax data. Includes differentially expressed genes (DEGs) between male and female samples, and differences in cell type abundance, related to Figure 1.**
**(A)** Representative photo of mated, sugar-fed female (top) and male (bottom) thorax dissection with anatomical diagram (in orange). Two samples (10x Genomics libraries) yielded 19,380 nuclei from 180 animals, post-quality control filtering Scale bar: 500 μm.
**(B)** UMAP of thorax nuclei, colored by sample (female samples = 1, male samples = 1)
**(C)** UMAP of thorax nuclei clustered using the Leiden algorithm (resolution = 5) for annotation.
**(D)** UMAP of nuclei from both thorax samples, colored and numbered by manual annotation using selected marker genes as listed in legend at the right of the figure panel. Categorization of cell type annotations represented by shaded gray headers. See Table S1 for gene IDs and thresholds
**(E-G)** Quantification of differences between male and female samples. Numbered and colored circles above (E) correspond to annotations in (D).
**(E)** Dot plot of number of differentially expressed genes between females and males in each cell type. Dots represent relative number of differentially expressed genes upregulated in male or female with a |log fold change| > 1 and false discovery rate < 0.05, determined by MAST on normalized expression and converted to $\log_2$ fold change.
**(F)** Sexual abundance difference index from data in (G). Each cell type was categorized based on the following index: Female biased (purple) if 0.3 < abundance index; Neutral (grey) if -0.3 ≤ abundance index ≤ 0.3; Male biased (yellow) if abundance index < -0.3.
**(G)** Bar plot of relative cell abundance, represented as the percent of each cell type of all nuclei collected from each sex for tissue. Separated by female (purple) and male (yellow).

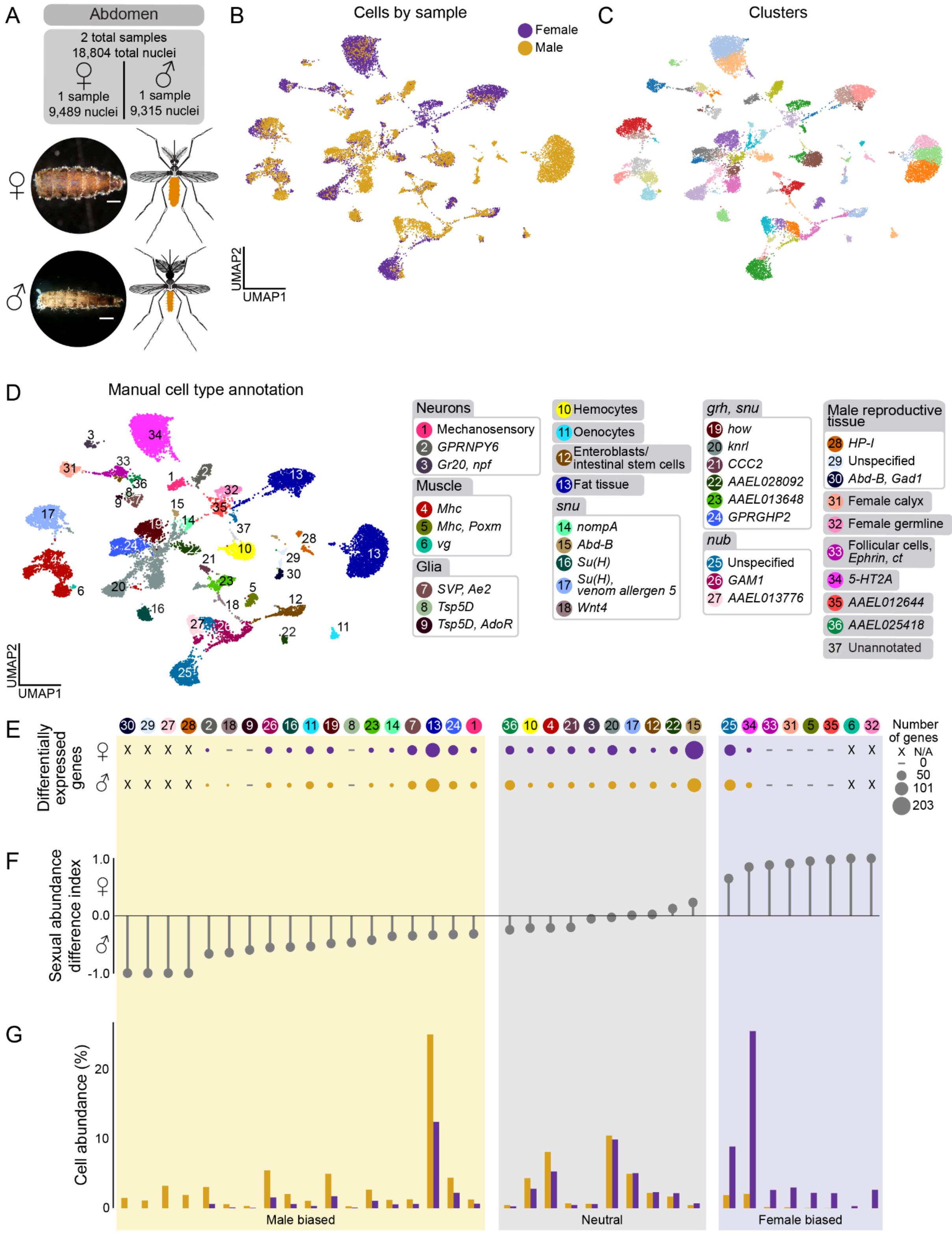

A
Abdomen
2 total samples
18,804 total nuclei
♀ 1 sample 9,489 nuclei
♂ 1 sample 9,315 nuclei
B
Cells by sample
Female
Male
UMAP2
UMAP1
C
Clusters
D
Manual cell type annotation
Neurons
1 Mechanosensory
2 GPRNPY6
3 Gr20, npf
Muscle
4 Mhc
5 Mhc, Poxm
6 vg
Glia
7 SVP, Ae2
8 Tsp5D
9 Tsp5D, AdoR
10 Hemocytes
11 Oenocytes
12 Enteroblasts/ intestinal stem cells
13 Fat tissue
snu
14 nompA
15 Abd-B
16 Su(H)
17 Su(H), venom allergen 5
18 Wnt4
grh, snu
19 how
20 knrl
21 CCC2
22 AAEL028092
23 AAEL013648
24 GPRGHP2
nub
25 Unspecified
26 GAM1
27 AAEL013776
Male reproductive tissue
28 HP-I
29 Unspecified
30 Abd-B, Gad1
31 Female calyx
32 Female germline
33 Follicular cells, Ephrin, ct
34 5-HT2A
35 AAEL012644
36 AAEL025418
37 Unannotated
E
Differentially expressed genes
Number of genes
X N/A
– 0
50
101
203
F
Sexual abundance difference index
1.0
0.0
-1.0
G
Cell abundance (%)
20
10
0
Male biased
Neutral
Female biased

**Data S2.3. Manual cell type annotation of female and male abdomen data. Includes differentially expressed genes (DEGs) between male and female samples, and differences in cell type abundance, related to Figure 1.**
**(A)** Representative photo of mated, sugar-fed female (top) and male (bottom) abdomen dissection with anatomical diagram (in orange). Two samples (10x Genomics libraries) yielded 18,804 nuclei from 99 animals, post-quality control filtering. Scale bar: 500 μm.
**(B)** UMAP of abdomen nuclei, colored by sample (female samples = 1, male samples = 1)
**(C)** UMAP of abdomen nuclei clustered using the Leiden algorithm (resolution = 5) for annotation.
**(D)** UMAP of nuclei from both abdomen samples, colored and numbered by manual annotation using selected marker genes as listed in legend at the right of the figure panel. Categorization of cell type annotations represented by shaded gray headers. See Table S1 for gene IDs and thresholds
**(E-G)** Quantification of differences between male and female samples. Numbered and colored circles above (E) correspond to annotations in (D).
**(E)** Dot plot of number of differentially expressed genes between females and males in each cell type. Dots represent relative number of differentially expressed genes upregulated in male or female with a |log fold change| > 1 and false discovery rate < 0.05, determined by MAST on normalized expression and converted to $\log_2$ fold change.
**(F)** Sexual abundance difference index from data in (G). Each cell type was categorized based on the following index: Female biased (purple) if 0.3 < abundance index; Neutral (grey) if -0.3 ≤ abundance index ≤ 0.3; Male biased (yellow) if abundance index < -0.3.
**(G)** Bar plot of relative cell abundance, represented as the percent of each cell type of all nuclei collected from each sex for tissue. Separated by female (purple) and male (yellow).

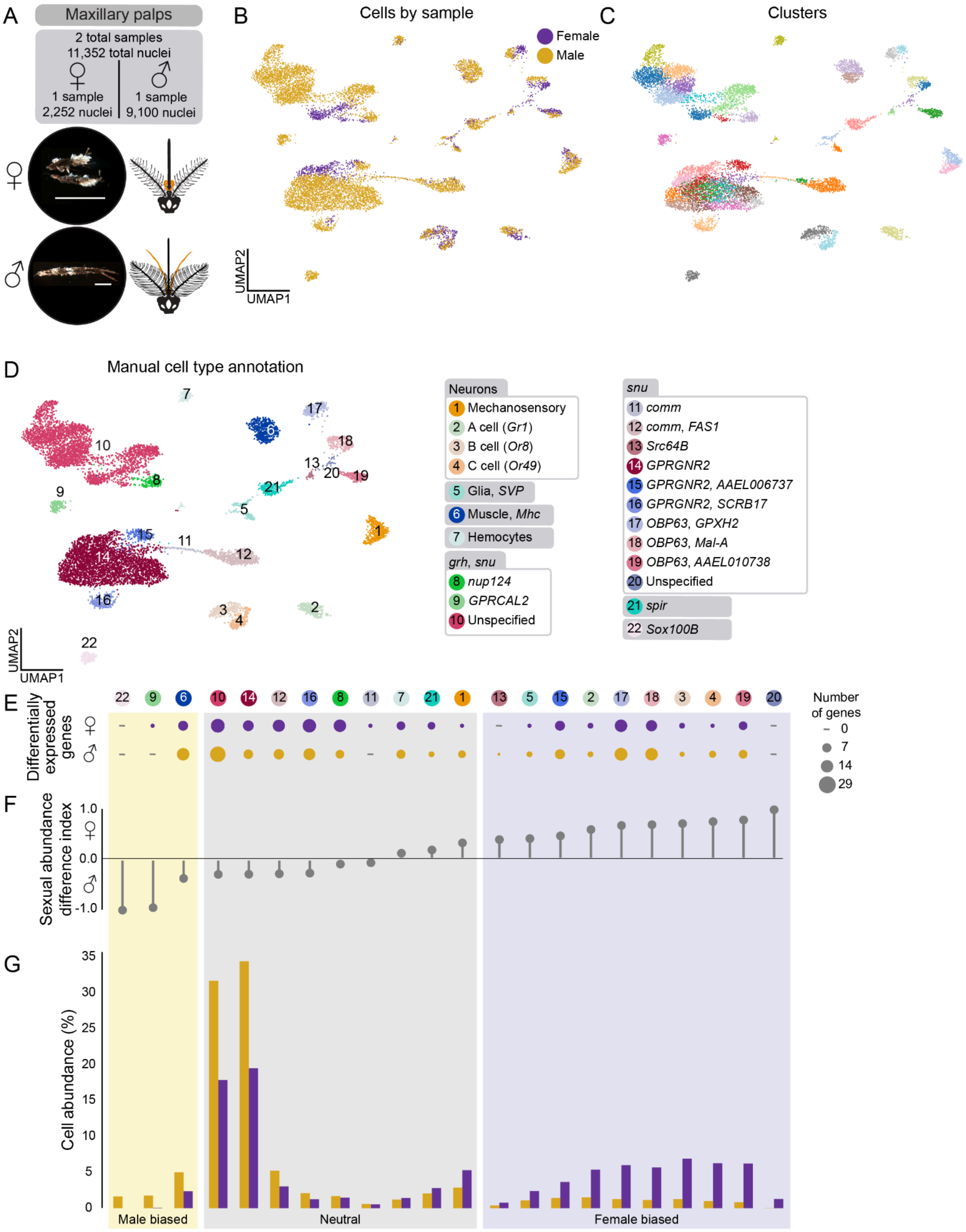
A
Maxillary palps
2 total samples
11,352 total nuclei
1 sample
2,252 nuclei
1 sample
9,100 nuclei
B
Cells by sample
Female
Male
UMAP2
UMAP1
C
Clusters
D
Manual cell type annotation
Neurons
1 Mechanosensory
2 A cell (Gr1)
3 B cell (Or8)
4 C cell (Or49)
5 Glia, SVP
6 Muscle, Mhc
7 Hemocytes
grh, snu
8 nup124
9 GPRCAL2
10 Unspecified
snu
11 comm
12 comm, FAS1
13 Src64B
14 GPRGNR2
15 GPRGNR2, AAEL006737
16 GPRGNR2, SCRB17
17 OBP63, GPXH2
18 OBP63, Mal-A
19 OBP63, AAEL010738
20 Unspecified
21 spir
22 Sox100B
E
Differentially expressed genes
Number of genes
0
7
14
29
F
Sexual abundance difference index
1.0
0.0
-1.0
G
Cell abundance (%)
35
30
25
20
15
10
5
0
Male biased
Neutral
Female biased

**Data S2.4. Manual cell type annotation of female and male maxillary palps data. Includes differentially expressed genes (DEGs) between male and female samples, and differences in cell type abundance, related to Figure 1.**
**(A)** Representative photo of mated, sugar-fed female (top) and male (bottom) maxillary palp dissection with anatomical diagram (in orange). Two samples (10x Genomics libraries) yielded 11,352 nuclei from 1,967 animals, post-quality control filtering. Scale bar: 500 μm.
**(B)** UMAP of maxillary palp nuclei, colored by sample (female samples = 1, male samples = 1)
**(C)** UMAP of maxillary palp nuclei clustered using the Leiden algorithm (resolution = 5) for annotation.
**(D)** UMAP of nuclei from both maxillary palp samples, colored and numbered by manual annotation using selected marker genes as listed in legend at the right of the figure panel. Categorization of cell type annotations represented by shaded gray headers. See Table S1 for gene IDs and thresholds
**(E-G)** Quantification of differences between male and female samples. Numbered and colored circles above (E) correspond to annotations in (D).
**(E)** Dot plot of number of differentially expressed genes between females and males in each cell type. Dots represent relative number of differentially expressed genes upregulated in male or female with a |log fold change| > 1 and false discovery rate < 0.05, determined by MAST on normalized expression and converted to $\log_2$ fold change.
**(F)** Sexual abundance difference index from data in (G). Each cell type was categorized based on the following index: Female biased (purple) if 0.3 < abundance index; Neutral (grey) if -0.3 ≤ abundance index ≤ 0.3; Male biased (yellow) if abundance index < -0.3.
**(G)** Bar plot of relative cell abundance, represented as the percent of each cell type of all nuclei collected from each sex for tissue. Separated by female (purple) and male (yellow).

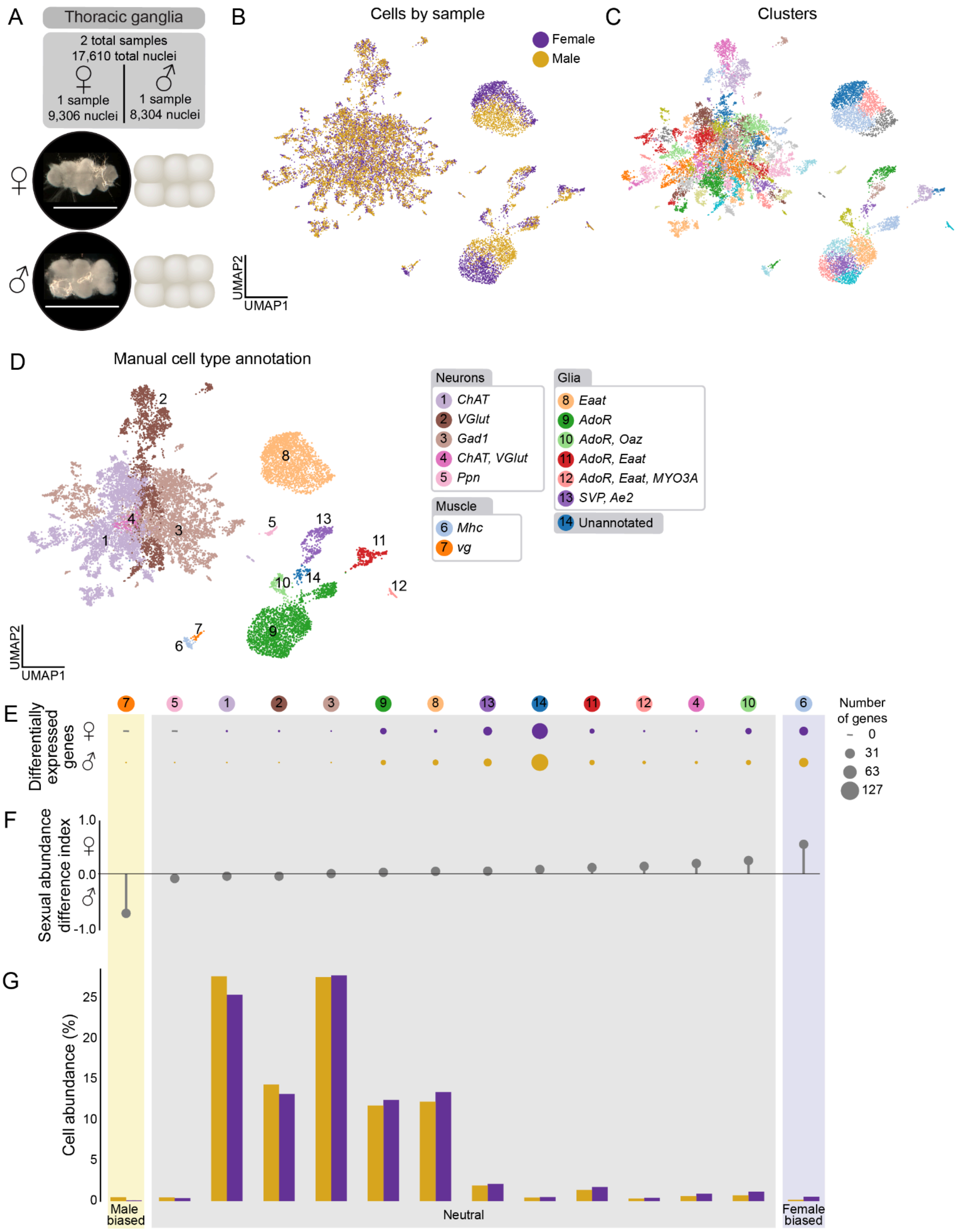

A
Thoracic ganglia
2 total samples
17,610 total nuclei
1 sample
9,306 nuclei
1 sample
8,304 nuclei
B
Cells by sample
Female
Male
UMAP2
UMAP1
C
Clusters
D
Manual cell type annotation
Neurons
1 ChAT
2 VGlut
3 Gad1
4 ChAT, VGlut
5 Ppn
Muscle
6 Mhc
7 vg
Glia
8 Eaat
9 AdoR
10 AdoR, Oaz
11 AdoR, Eaat
12 AdoR, Eaat, MYO3A
13 SVP, Ae2
14 Unannotated
E
Differentially expressed genes
Number of genes
0
31
63
127
F
Sexual abundance difference index
1.0
0.0
-1.0
G
Cell abundance (%)
25
20
15
10
5
0
Male biased
Neutral
Female biased

**Data S2.5. Manual cell type annotation of female and male thoracic ganglia data. Includes differentially expressed genes (DEGs) between male and female samples, and differences in cell type abundance, related to Figure 1.**
**(A)** Representative photo of mated, sugar-fed female (top) and male (bottom) thoracic ganglia dissection with anatomical diagram. Two samples (10x Genomics libraries) yielded 17,610 nuclei from 240 animals, post-quality control filtering. Scale bar: 500 μm.
**(B)** UMAP of thoracic ganglia nuclei, colored by sample (female samples = 1, male samples = 1)
**(C)** UMAP of thoracic ganglia nuclei clustered using the Leiden algorithm (resolution = 5) for annotation.
**(D)** UMAP of nuclei from both thoracic ganglia samples, colored and numbered by manual annotation using selected marker genes as listed in legend at the right of the figure panel. Categorization of cell type annotations represented by shaded gray headers. See Table S1 for gene IDs and thresholds.
**(E-G)** Quantification of differences between male and female samples. Numbered and colored circles above (E) correspond to annotations in (D).
**(E)** Dot plot of number of differentially expressed genes between females and males in each cell type. Dots represent relative number of differentially expressed genes upregulated in male or female with a |log fold change| > 1 and false discovery rate < 0.05, determined by MAST on normalized expression and converted to $\log_2$ fold change.
**(F)** Sexual abundance difference index from data in (G). Each cell type was categorized based on the following index: Female biased (purple) if 0.3 < abundance index; Neutral (grey) if -0.3 ≤ abundance index ≤ 0.3; Male biased (yellow) if abundance index < -0.3.
**(G)** Bar plot of relative cell abundance, represented as the percent of each cell type of all nuclei collected from each sex for tissue. Separated by female (purple) and male (yellow).

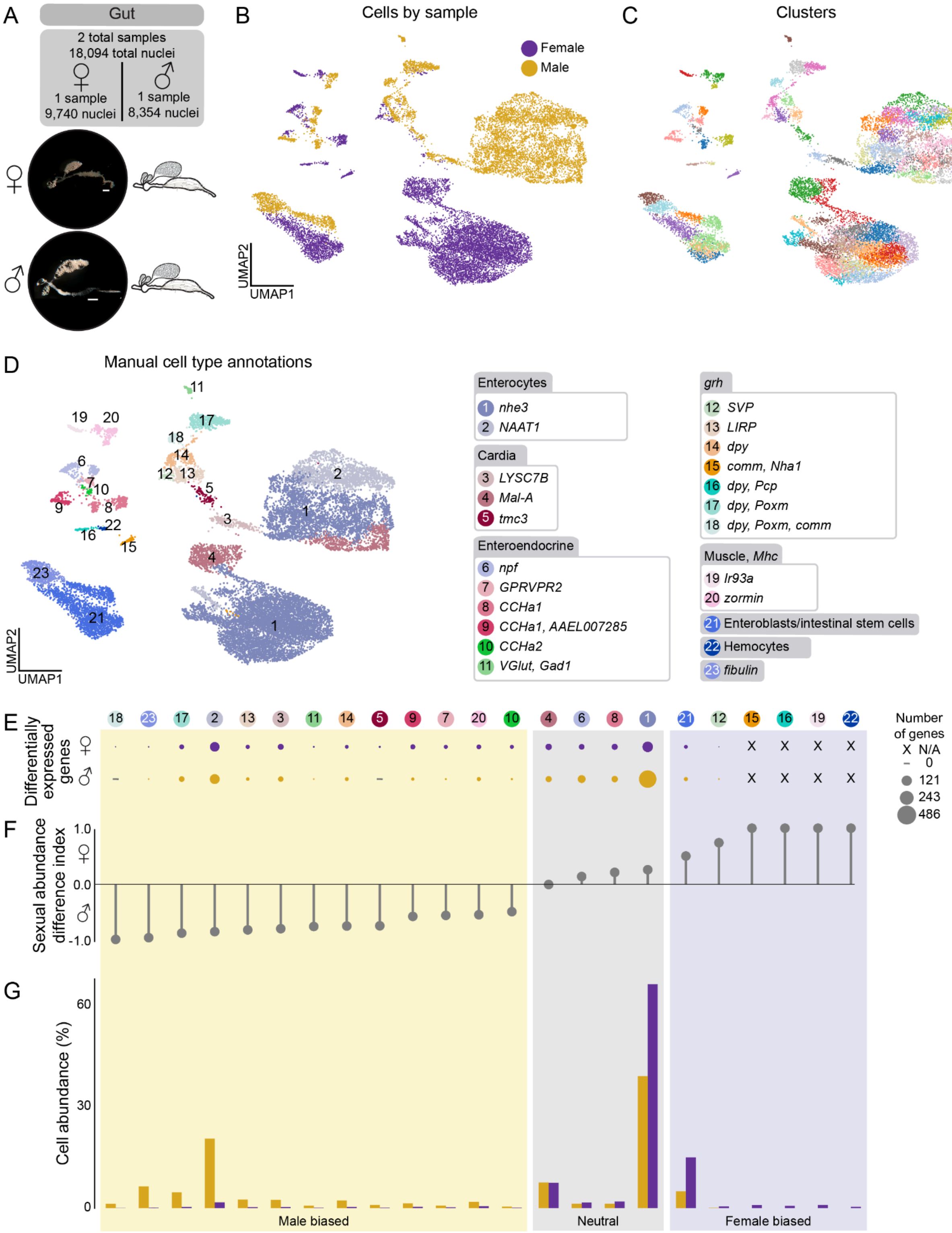
A
Gut
2 total samples
18,094 total nuclei
1 sample
9,740 nuclei
1 sample
8,354 nuclei
B
Cells by sample
Female
Male
UMAP2
UMAP1
C
Clusters
D
Manual cell type annotations
Enterocytes
1 nhe3
2 NAAT1
Cardia
3 LYSC7B
4 Mal-A
5 tmc3
Enteroendocrine
6 npf
7 GPRVPR2
8 CCHa1
9 CCHa1, AAEL007285
10 CCHa2
11 VGlut, Gad1
grh
12 SVP
13 LIRP
14 dpy
15 comm, Nha1
16 dpy, Pcp
17 dpy, Poxm
18 dpy, Poxm, comm
Muscle, Mhc
19 Ir93a
20 zormin
21 Enteroblasts/intestinal stem cells
22 Hemocytes
23 fibulin
E
Differentially expressed genes
Number of genes
X N/A
– 0
121
243
486
F
Sexual abundance difference index
1.0
0.0
-1.0
G
Cell abundance (%)
60
30
0
Male biased
Neutral
Female biased

**Data S2.6. Manual cell type annotation of female and male gut data. Includes differentially expressed genes (DEGs) between male and female samples, and differences in cell type abundance, related to Figure 1.**

**(A)** Representative photo of mated, sugar-fed female (top) and male (bottom) gut dissection with anatomical diagram. Two samples (10x Genomics libraries) yielded 18,094 nuclei from 180 animals, post-quality control filtering. Scale bar: 500 µm.

**(B)** UMAP of gut nuclei, colored by sample (female samples = 1, male samples = 1)

**(C)** UMAP of gut nuclei clustered using the Leiden algorithm (resolution = 5) for annotation.

**(D)** UMAP of nuclei from both gut samples, colored and numbered by manual annotation using selected marker genes as listed in legend at the right of the figure panel. Categorization of cell type annotations represented by shaded gray headers. See Table S1 for gene IDs and thresholds

**(E-G)** Quantification of differences between male and female samples. Numbered and colored circles above (E) correspond to annotations in (D).

**(E)** Dot plot of number of differentially expressed genes between females and males in each cell type. Dots represent relative number of differentially expressed genes upregulated in male or female with a |log fold change| > 1 and false discovery rate < 0.05, determined by MAST on normalized expression and converted to $\log_2$ fold change.

**(F)** Sexual abundance difference index from data in (G). Each cell type was categorized based on the following index: Female biased (purple) if 0.3 < abundance index; Neutral (grey) if -0.3 ≤ abundance index ≤ 0.3; Male biased (yellow) if abundance index < -0.3.

**(G)** Bar plot of relative cell abundance, represented as the percent of each cell type of all nuclei collected from each sex for tissue. Separated by female (purple) and male (yellow).

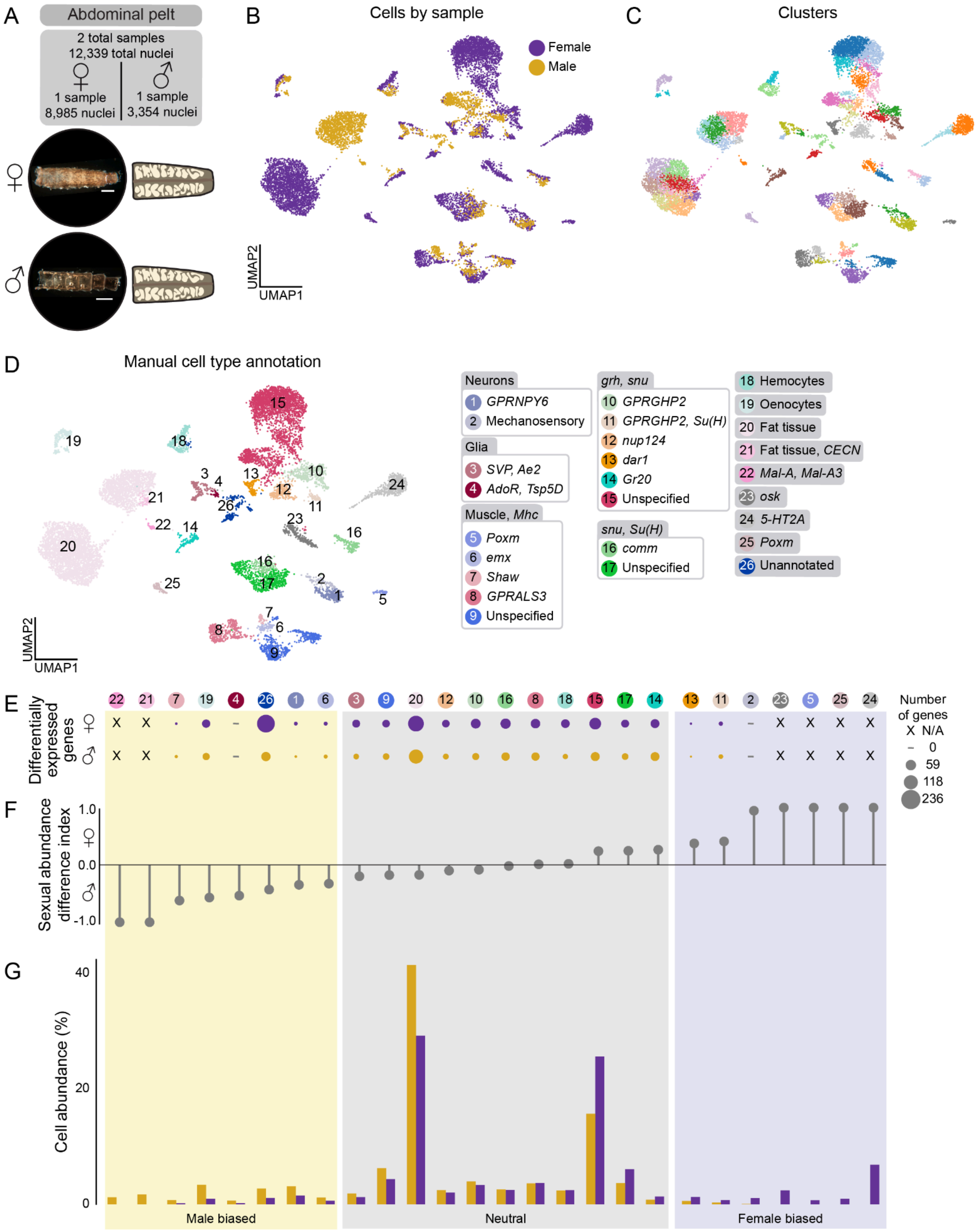
A
Abdominal pelt
2 total samples
12,339 total nuclei
♀
1 sample
8,985 nuclei
♂
1 sample
3,354 nuclei
B
Cells by sample
Female
Male
UMAP2
UMAP1
C
Clusters
D
Manual cell type annotation
Neurons
1 GPRNPY6
2 Mechanosensory
Glia
3 SVP, Ae2
4 AdoR, Tsp5D
Muscle, Mhc
5 Poxm
6 emx
7 Shaw
8 GPRALS3
9 Unspecified
grh, snu
10 GPRGHP2
11 GPRGHP2, Su(H)
12 nup124
13 dar1
14 Gr20
15 Unspecified
snu, Su(H)
16 comm
17 Unspecified
18 Hemocytes
19 Oenocytes
20 Fat tissue
21 Fat tissue, CECN
22 Mal-A, Mal-A3
23 osk
24 5-HT2A
25 Poxm
26 Unannotated
E
Differentially expressed genes
Number of genes
X N/A
0
59
118
236
F
Sexual abundance difference index
1.0
0.0
-1.0
G
Cell abundance (%)
40
20
0
Male biased
Neutral
Female biased

**Data S2.7. Manual cell type annotation of female and male abdominal pelt data. Includes differentially expressed genes (DEGs) between male and female samples, and differences in cell type abundance, related to Figure 1.**

**(A)** Representative photo of mated, sugar-fed female (top) and male (bottom) abdominal pelt dissection with anatomical diagram. Two samples (10x Genomics libraries) yielded 12,339 nuclei from 180 animals, post-quality control filtering. Scale bar: 500 μm.

**(B)** UMAP of abdominal pelt nuclei, colored by sample (female samples = 1, male samples = 1)

**(C)** UMAP of abdominal pelt nuclei clustered using the Leiden algorithm (resolution = 5) for annotation.

**(D)** UMAP of nuclei from both abdominal pelt samples, colored and numbered by manual annotation using selected marker genes as listed in legend at the right of the figure panel. Categorization of cell type annotations represented by shaded gray headers. See Table S1 for gene IDs and thresholds

**(E-G)** Quantification of differences between male and female samples. Numbered and colored circles above (E) correspond to annotations in (D).

**(E)** Dot plot of number of differentially expressed genes between females and males in each cell type. Dots represent relative number of differentially expressed genes upregulated in male or female with a |log fold change| > 1 and false discovery rate < 0.05, determined by MAST on normalized expression and converted to $\log_2$ fold change.

**(F)** Sexual abundance difference index from data in (G). Each cell type was categorized based on the following index: Female biased (purple) if 0.3 < abundance index; Neutral (grey) if -0.3 ≤ abundance index ≤ 0.3; Male biased (yellow) if abundance index < -0.3.

**(G)** Bar plot of relative cell abundance, represented as the percent of each cell type of all nuclei collected from each sex for tissue. Separated by female (purple) and male (yellow).

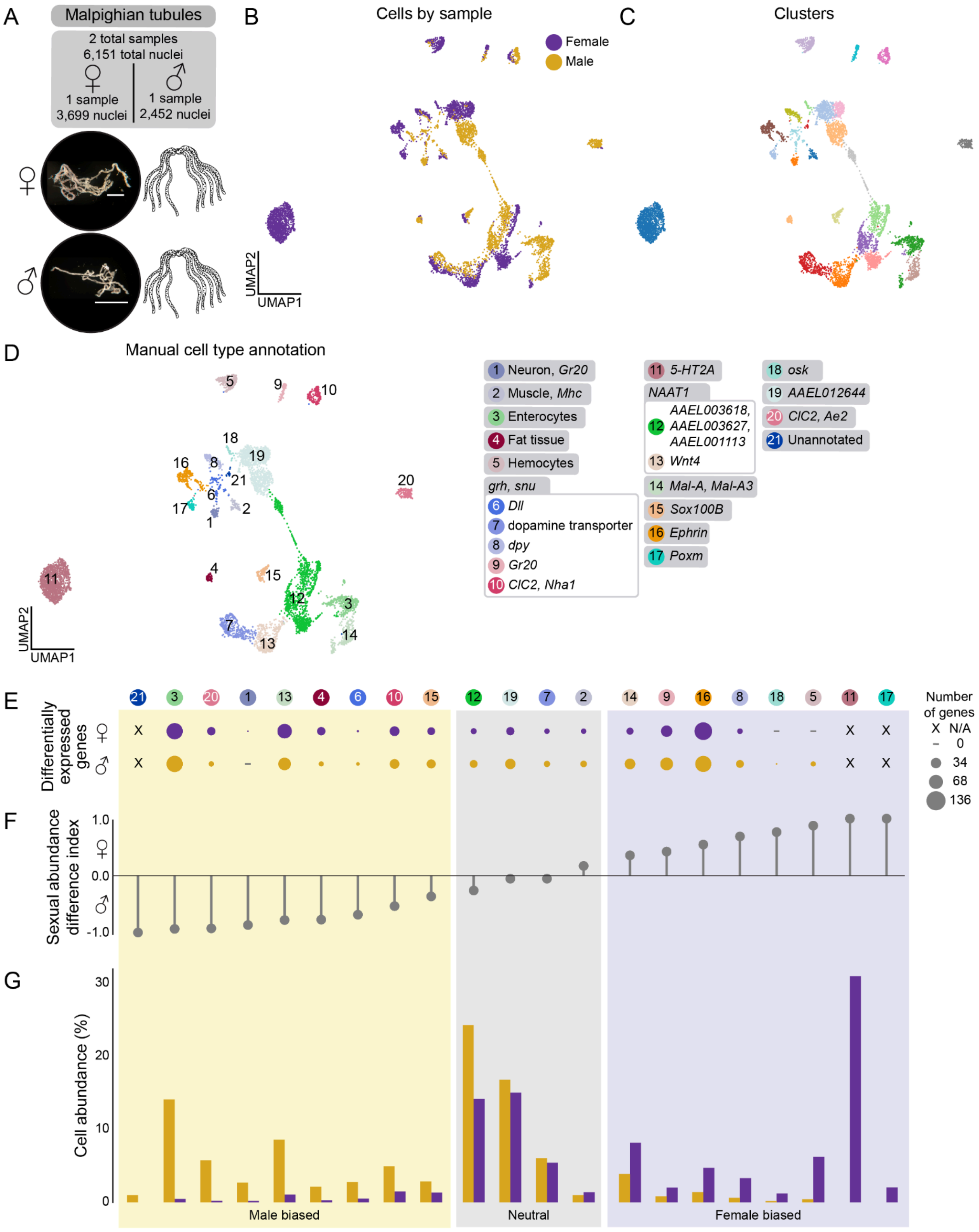

A
Malpighian tubules
2 total samples
6,151 total nuclei
1 sample
3,699 nuclei
1 sample
2,452 nuclei
B
Cells by sample
Female
Male
UMAP2
UMAP1
C
Clusters
D
Manual cell type annotation
1 Neuron, Gr20
2 Muscle, Mhc
3 Enterocytes
4 Fat tissue
5 Hemocytes
grh, snu
6 Dll
7 dopamine transporter
8 dpy
9 Gr20
10 ClC2, Nha1
11 5-HT2A
NAAT1
12 AAEL003618, AAEL003627, AAEL001113
13 Wnt4
14 Mal-A, Mal-A3
15 Sox100B
16 Ephrin
17 Poxm
18 osk
19 AAEL012644
20 ClC2, Ae2
21 Unannotated
E
Differentially expressed genes
Number of genes
X N/A
0
34
68
136
F
Sexual abundance difference index
1.0
0.0
-1.0
G
Cell abundance (%)
30
20
10
0
Male biased
Neutral
Female biased

**Data S2.8. Manual cell type annotation of female and male malpighian tubules data. Includes differentially expressed genes (DEGs) between male and female samples, and differences in cell type abundance, related to Figure 1.**

**(A)** Representative photo of mated, sugar-fed female (top) and male (bottom) malpighian tubule dissection with anatomical diagram. Two samples (10x Genomics libraries) yielded 6,151 nuclei from 309 animals, post-quality control filtering. Scale bar: 500 μm.

**(B)** UMAP of malpighian tubules nuclei, colored by sample (female samples = 1, male samples = 1)

**(C)** UMAP of malpighian tubules nuclei clustered using the Leiden algorithm (resolution = 5) for annotation.

**(D)** UMAP of nuclei from both malpighian tubules samples, colored and numbered by manual annotation using selected marker genes as listed in legend at the right of the figure panel. Categorization of cell type annotations represented by shaded gray headers. See Table S1 for gene IDs and thresholds

**(E-G)** Quantification of differences between male and female samples. Numbered and colored circles above (E) correspond to annotations in (D).

**(E)** Dot plot of number of differentially expressed genes between females and males in each cell type. Dots represent relative number of differentially expressed genes upregulated in male or female with a |log fold change| > 1 and false discovery rate < 0.05, determined by MAST on normalized expression and converted to $\log_2$ fold change.

**(F)** Sexual abundance difference index from data in (G). Each cell type was categorized based on the following index: Female biased (purple) if 0.3 < abundance index; Neutral (grey) if -0.3 ≤ abundance index ≤ 0.3; Male biased (yellow) if abundance index < -0.3.

**(G)** Bar plot of relative cell abundance, represented as the percent of each cell type of all nuclei collected from each sex for tissue. Separated by female (purple) and male (yellow).

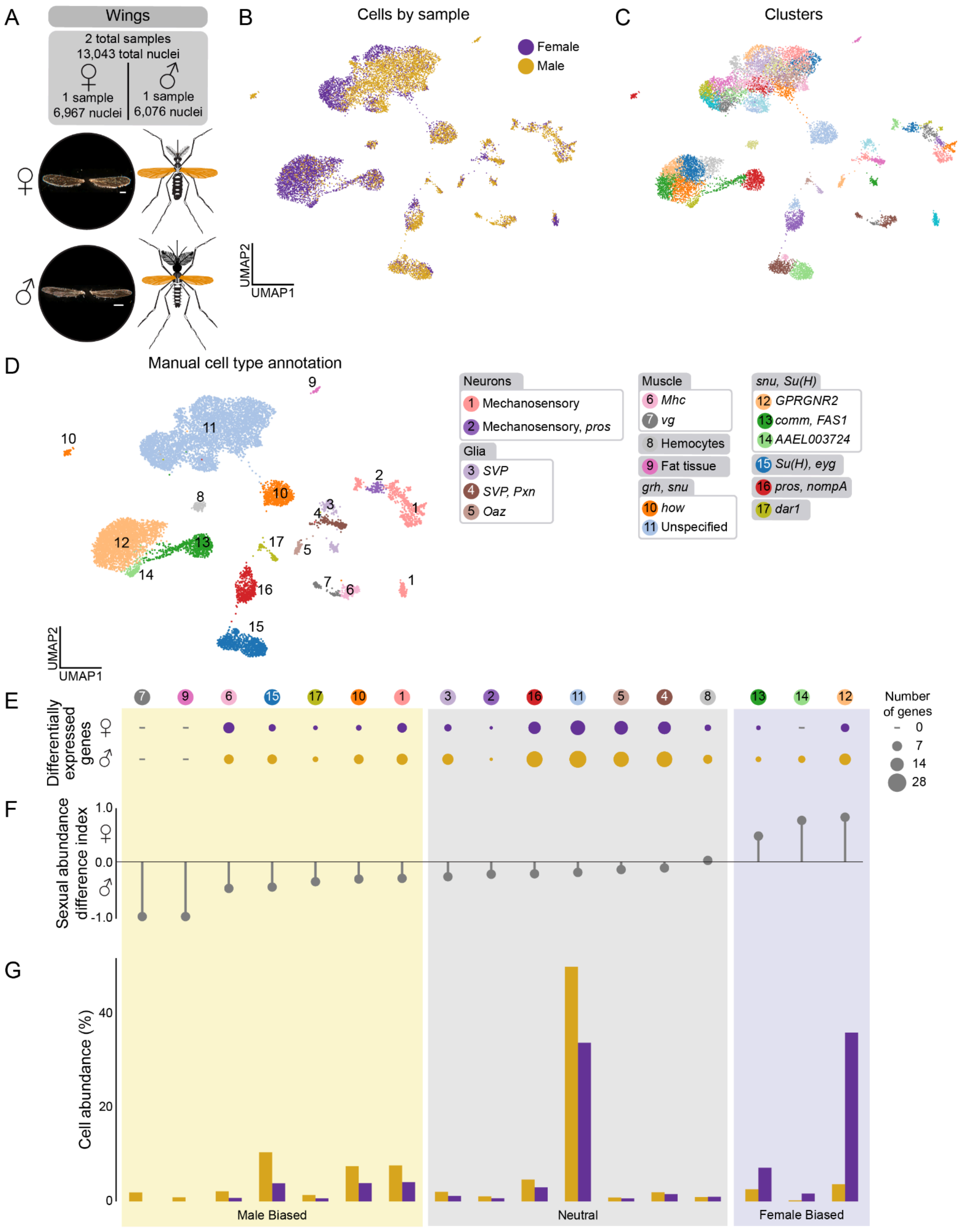

A
Wings
2 total samples
13,043 total nuclei
♀
1 sample
6,967 nuclei
♂
1 sample
6,076 nuclei
B
Cells by sample
Female
Male
UMAP2
UMAP1
C
Clusters
D
Manual cell type annotation
Neurons
1 Mechanosensory
2 Mechanosensory, pros
Glia
3 SVP
4 SVP, Pxn
5 Oaz
Muscle
6 Mhc
7 vg
8 Hemocytes
9 Fat tissue
grh, snu
10 how
11 Unspecified
snu, Su(H)
12 GPRGNR2
13 comm, FAS1
14 AAEL003724
15 Su(H), eyg
16 pros, nompA
17 dar1
E
Differentially expressed genes
Number of genes
0
7
14
28
F
Sexual abundance difference index
1.0
0.0
-1.0
G
Cell abundance (%)
40
20
0
Male Biased
Neutral
Female Biased

**Data S2.9. Manual cell type annotation of female and male wings data. Includes differentially expressed genes (DEGs) between male and female samples, and differences in cell type abundance, related to Figure 1.**

**(A)** Representative photo of mated, sugar-fed female (top) and male (bottom) wing dissection with anatomical diagram (in orange). Two samples (10x Genomics libraries) yielded 13,043 nuclei from 528 animals, post-quality control filtering. Scale bar: 500 μm.

**(B)** UMAP of wing nuclei, colored by sample (female samples = 1, male samples = 1)

**(C)** UMAP of wing nuclei clustered using the Leiden algorithm (resolution = 5) for annotation.

**(D)** UMAP of nuclei from both wing samples, colored and numbered by manual annotation using selected marker genes as listed in legend at the right of the figure panel. Categorization of cell type annotations represented by shaded gray headers. See Table S1 for gene IDs and thresholds

**(E-G)** Quantification of differences between male and female samples. Numbered and colored circles above (E) correspond to annotations in (D).

**(E)** Dot plot of number of differentially expressed genes between females and males in each cell type. Dots represent relative number of differentially expressed genes upregulated in male or female with a |log fold change| > 1 and false discovery rate < 0.05, determined by MAST on normalized expression and converted to $\log_2$ fold change.

**(F)** Sexual abundance difference index from data in (G). Each cell type was categorized based on the following index: Female biased (purple) if 0.3 < abundance index; Neutral (grey) if -0.3 ≤ abundance index ≤ 0.3; Male biased (yellow) if abundance index < -0.3.

**(G)** Bar plot of relative cell abundance, represented as the percent of each cell type of all nuclei collected from each sex for tissue. Separated by female (purple) and male (yellow).

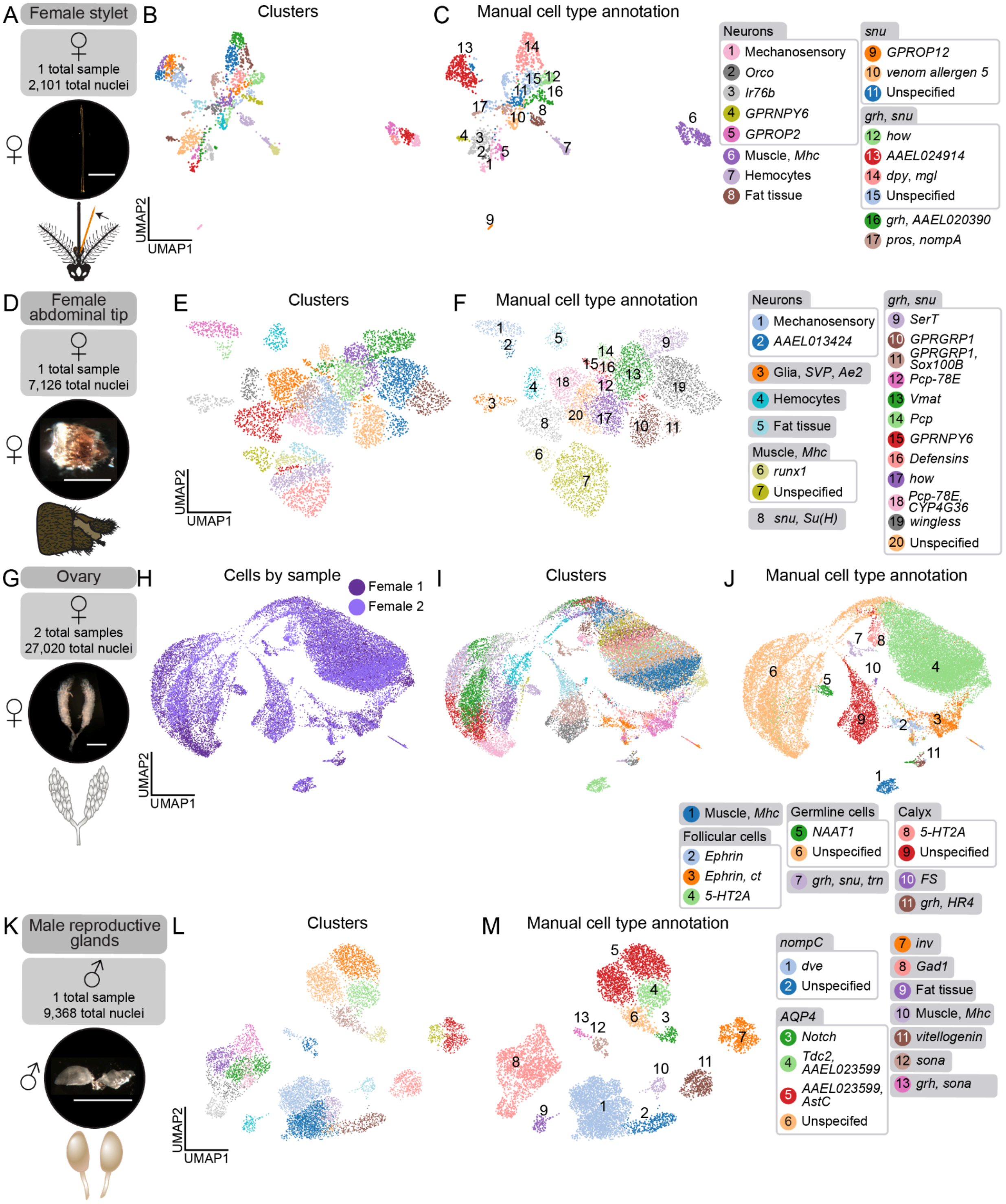

A Female stylet
1 total sample
2,101 total nuclei
B Clusters
UMAP2
UMAP1
C Manual cell type annotation
Neurons
1 Mechanosensory
2 Orco
3 Ir76b
4 GPRNPY6
5 GPROP2
6 Muscle, Mhc
7 Hemocytes
8 Fat tissue
snu
9 GPROP12
10 venom allergen 5
11 Unspecified
grh, snu
12 how
13 AAEL024914
14 dpy, mgl
15 Unspecified
16 grh, AAEL020390
17 pros, nompA
D Female abdominal tip
1 total sample
7,126 total nuclei
E Clusters
UMAP2
UMAP1
F Manual cell type annotation
Neurons
1 Mechanosensory
2 AAEL013424
3 Glia, SVP, Ae2
4 Hemocytes
5 Fat tissue
Muscle, Mhc
6 runx1
7 Unspecified
8 snu, Su(H)
grh, snu
9 SerT
10 GPRGRP1
11 GPRGRP1, Sox100B
12 Pcp-78E
13 Vmat
14 Pcp
15 GPRNPY6
16 Defensins
17 how
18 Pcp-78E, CYP4G36
19 wingless
20 Unspecified
G Ovary
2 total samples
27,020 total nuclei
H Cells by sample
Female 1
Female 2
UMAP2
UMAP1
I Clusters
J Manual cell type annotation
1 Muscle, Mhc
Follicular cells
2 Ephrin
3 Ephrin, ct
4 5-HT2A
Germline cells
5 NAAT1
6 Unspecified
7 grh, snu, trn
Calyx
8 5-HT2A
9 Unspecified
10 FS
11 grh, HR4
K Male reproductive glands
1 total sample
9,368 total nuclei
L Clusters
UMAP2
UMAP1
M Manual cell type annotation
nompC
1 dve
2 Unspecified
AQP4
3 Notch
4 Tdc2, AAEL023599
5 AAEL023599, AstC
6 Unspecifed
7 inv
8 Gad1
9 Fat tissue
10 Muscle, Mhc
11 vitellogenin
12 sona
13 grh, sona

**Data S2.10. Manual cell type annotation of female stylet, female abdominal tip, female ovary, and male reproductive glands, related to Figure 1.**

**(A)** Representative photo of mated, sugar-fed female stylet dissection with anatomical diagram (in orange). One sample (10x Genomics library) yielded 2,101 nuclei from 1,747 animals, post-quality control filtering. Scale bar: 500 µm.

**(B)** UMAP of female stylet nuclei clustered using the Leiden algorithm (resolution = 5) for annotation.

**(C)** UMAP of nuclei from female stylet sample, colored and numbered by manual annotation using selected marker genes as listed in legend at the right of the figure panel. Categorization of cell type annotations represented by shaded gray headers. See Table S1 for gene IDs and thresholds

**(D)** Representative photo of mated, sugar-fed female abdominal tip dissection with anatomical diagram. One sample (10x Genomics library) yielded 7,126 nuclei from 89 animals, post-quality control filtering. Scale bar: 500 µm.

**(E)** UMAP of female abdominal tip nuclei clustered using the Leiden algorithm (resolution = 5) for annotation.

**(F)** UMAP of nuclei from female abdominal tip sample, colored and numbered by manual annotation using selected marker genes as listed in legend at the right of the figure panel. Categorization of cell type annotations represented by shaded gray headers. See Table S1 for gene IDs and thresholds

**(G)** Representative photo of mated, sugar-fed female ovary dissection with anatomical diagram. Two samples (10x Genomics libraries) yielded 27,020 nuclei from 60 animals, post-quality control filtering. Scale bar: 500 µm.

**(H)** UMAP of ovary nuclei, colored by sample (female samples = 2)

**(I)** UMAP of ovary nuclei clustered using the Leiden algorithm (resolution = 5) for annotation.

**(J)** UMAP of nuclei from ovary samples, colored and numbered by manual annotation using selected marker genes as listed in legend at the bottom of figure panel. Categorization of cell type annotations represented by shaded gray headers. See Table S1 for gene IDs and thresholds

**(K)** Representative photo of mated, sugar-fed male reproductive gland dissection with anatomical diagram. One sample (10x Genomics library) yielded 9,368 nuclei from 264 animals, post-quality control filtering. Scale bar: 500 µm.

**(L)** UMAP of male reproductive gland nuclei clustered using the Leiden algorithm (resolution = 5) for annotation.

**(M)** UMAP of nuclei from male reproductive gland sample, colored and numbered by manual annotation using selected marker genes as listed in legend at the right of the figure panel. Categorization of cell type annotations represented by shaded gray headers. See Table S1 for gene IDs and thresholds

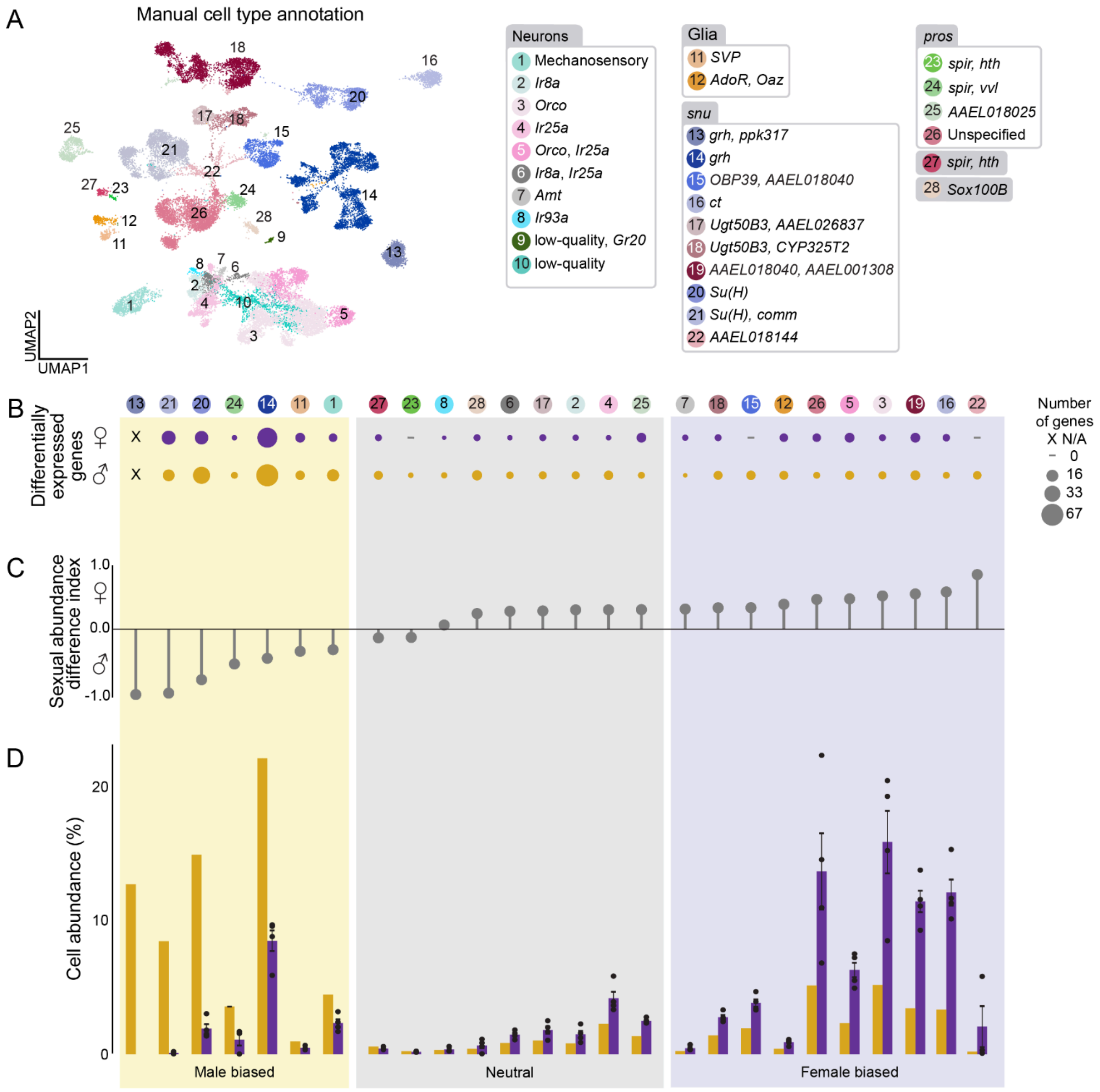
A
Manual cell type annotation
UMAP1
UMAP2
Neurons
1 Mechanosensory
2 Ir8a
3 Orco
4 Ir25a
5 Orco, Ir25a
6 Ir8a, Ir25a
7 Amt
8 Ir93a
9 low-quality, Gr20
10 low-quality
Glia
11 SVP
12 AdoR, Oaz
snu
13 grh, ppk317
14 grh
15 OBP39, AAEL018040
16 ct
17 Ugt50B3, AAEL026837
18 Ugt50B3, CYP325T2
19 AAEL018040, AAEL001308
20 Su(H)
21 Su(H), comm
22 AAEL018144
pros
23 spir, hth
24 spir, vvl
25 AAEL018025
26 Unspecified
27 spir, hth
28 Sox100B
B
Differentially expressed genes
Number of genes
X N/A
0
16
33
67
C
Sexual abundance difference index
1.0
0.0
-1.0
D
Cell abundance (%)
20
10
0
Male biased
Neutral
Female biased

**Data S2.11. Manual cell type annotation of female and male antennae data. Includes differentially expressed genes (DEGs) between male and female samples, and differences in cell type abundance, related to Figures 1 and 3.**
**(A)** UMAP of nuclei from all antennae samples, colored and numbered by manual annotation using selected marker genes as listed in legend at the right of the figure panel. Categorization of cell type annotations represented by shaded gray headers. See Table S1 for gene IDs and thresholds
**(B-D)** Quantification of differences between male and female samples. Numbered and colored circles above (C) correspond to annotations in (B).
**(B)** Dot plot of number of differentially expressed genes between females and males in each cell type. Dots represent relative number of differentially expressed genes upregulated in male or female with a |log fold change| > 1 and false discovery rate < 0.05, determined by MAST on normalized expression and converted to $\log_2$ fold change.
**(C)** Sexual abundance difference index from data in (G). Each cell type was categorized based on the following index: Female biased (purple) if 0.3 < abundance index; Neutral (grey) if -0.3 ≤ abundance index ≤ 0.3; Male biased (yellow) if abundance index < -0.3.
**(D)** Bar plot of relative cell abundance, represented as the percent of each cell type of all nuclei collected from each sex for tissue. Separated by female (purple) and male (yellow). Values for each replicate shown as dots, and standard error indicated by black brackets.

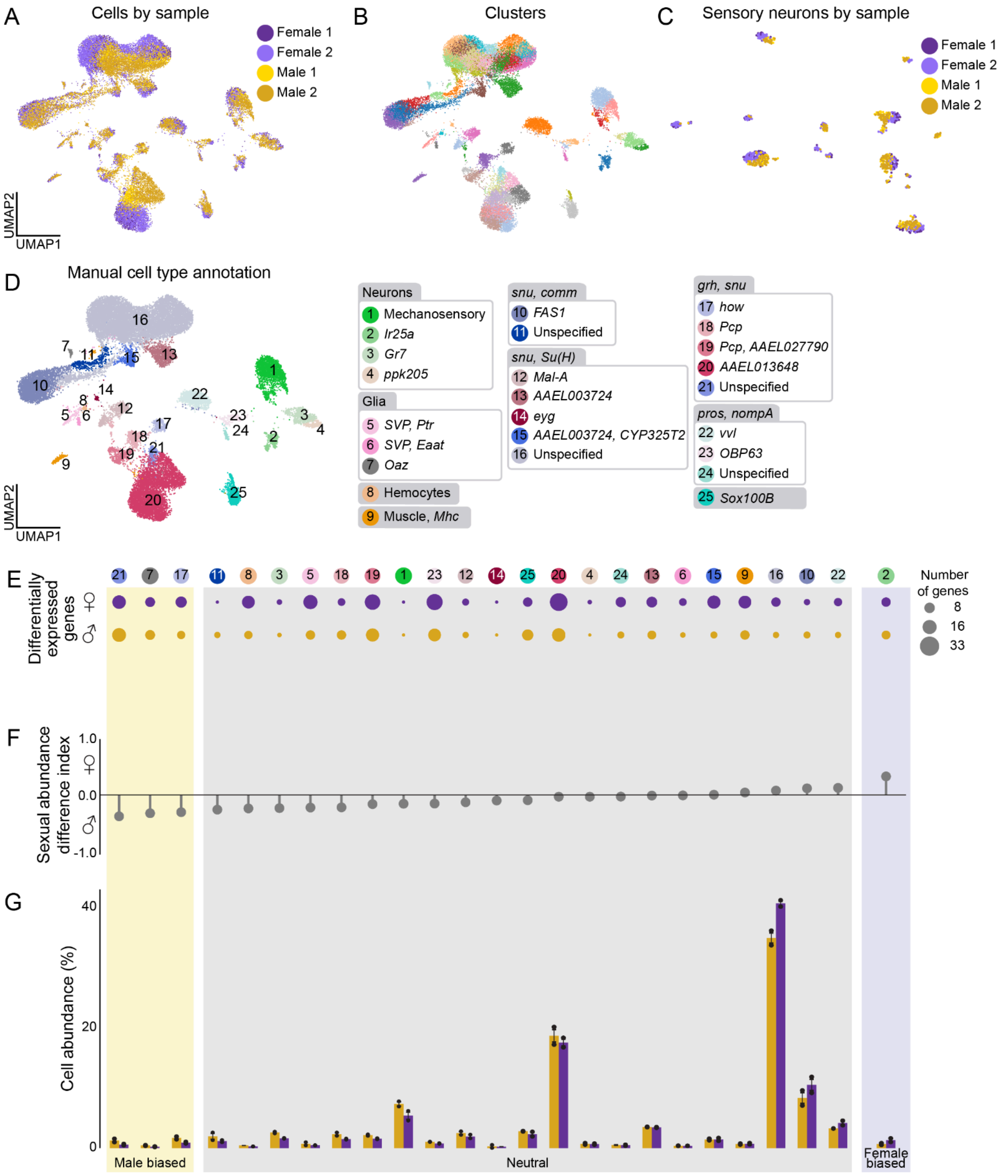
A
Cells by sample
Female 1
Female 2
Male 1
Male 2
UMAP2
UMAP1
B
Clusters
C
Sensory neurons by sample
Female 1
Female 2
Male 1
Male 2
D
Manual cell type annotation
UMAP2
UMAP1
Neurons
1 Mechanosensory
2 Ir25a
3 Gr7
4 ppk205
Glia
5 SVP, Ptr
6 SVP, Eaat
7 Oaz
8 Hemocytes
9 Muscle, Mhc
snu, comm
10 FAS1
11 Unspecified
snu, Su(H)
12 Mal-A
13 AAEL003724
14 eyg
15 AAEL003724, CYP325T2
16 Unspecified
grh, snu
17 how
18 Pcp
19 Pcp, AAEL027790
20 AAEL013648
21 Unspecified
pros, nompA
22 vvl
23 OBP63
24 Unspecified
25 Sox100B
E
Differentially expressed genes
♀
♂
Number of genes
8
16
33
F
Sexual abundance difference index
1.0
0.0
-1.0
G
Cell abundance (%)
40
20
0
Male biased
Neutral
Female biased

**Data S2.12. Manual cell type annotation of female and male tarsi data. Includes differentially expressed genes (DEGs) between male and female samples, and differences in cell type abundance, related to Figures 1, 4, and 5.**

**(A)** UMAP of tarsi nuclei, colored by sample (female samples = 2, male samples = 2)

**(B)** UMAP of tarsi nuclei clustered using the Leiden algorithm (resolution = 5).

**(C)** UMAP of *nompC*-negative tarsi sensory neuron nuclei, colored by sample (female samples = 2, male samples = 2), related to Figure 5C.

**(D)** UMAP of nuclei from all tarsi samples, colored and numbered by manual annotation using selected marker genes as listed in legend at the right of the figure panel. Categorization of cell type annotations represented by shaded gray headers. See Table S1 for gene IDs and thresholds

**(E-G)** Quantification of differences between male and female samples. Numbered and colored circles above (E) correspond to annotations in (D).

**(E)** Dot plot of number of differentially expressed genes between females and males in each cell type. Dots represent relative number of differentially expressed genes upregulated in male or female with a |log fold change| > 1 and false discovery rate < 0.05, determined by MAST on normalized expression and converted to $\log_2$ fold change.

**(F)** Sexual abundance difference index from data in (G). Each cell type was categorized based on the following index: Female biased (purple) if 0.3 < abundance index; Neutral (grey) if -0.3 ≤ abundance index ≤ 0.3; Male biased (yellow) if abundance index < -0.3.

**(G)** Bar plot of relative cell abundance, represented as the percent of each cell type of all nuclei collected from each sex for tissue. Separated by female (purple) and male (yellow). Values for each replicate are shown as dots, and standard error indicated by black brackets.

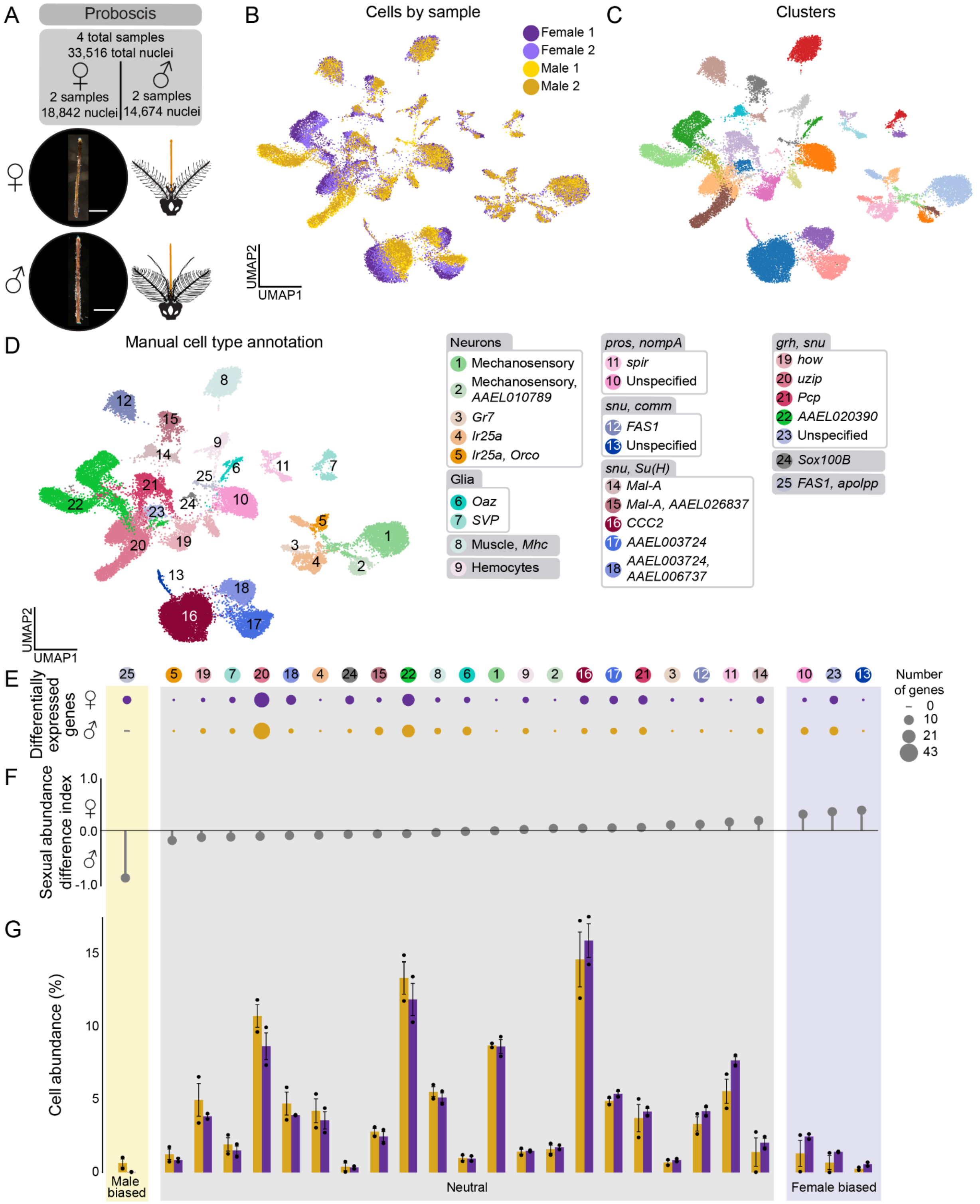
A
Proboscis
4 total samples
33,516 total nuclei
2 samples
18,842 nuclei
2 samples
14,674 nuclei
B
Cells by sample
Female 1
Female 2
Male 1
Male 2
UMAP2
UMAP1
C
Clusters
D
Manual cell type annotation
Neurons
1 Mechanosensory
2 Mechanosensory, AAEL010789
3 Gr7
4 Ir25a
5 Ir25a, Orco
Glia
6 Oaz
7 SVP
8 Muscle, Mhc
9 Hemocytes
pros, nompA
11 spir
10 Unspecified
snu, comm
12 FAS1
13 Unspecified
snu, Su(H)
14 Mal-A
15 Mal-A, AAEL026837
16 CCC2
17 AAEL003724
18 AAEL003724, AAEL006737
grh, snu
19 how
20 uzip
21 Pcp
22 AAEL020390
23 Unspecified
24 Sox100B
25 FAS1, apolpp
E
Differentially expressed genes
Number of genes
0
10
21
43
F
Sexual abundance difference index
1.0
0.0
-1.0
G
Cell abundance (%)
15
10
5
0
Male biased
Neutral
Female biased

**Data S2.13. Manual cell type annotation of female and male proboscis data. Includes differentially expressed genes (DEGs) between male and female samples, and differences in cell type abundance, related to Figures 1, 4, and 5.**
**(A)** Representative photo of mated, sugar-fed female (top) and male (bottom) proboscis dissection with anatomical diagram (in orange). Four samples (10x Genomics libraries) yielded 33,516 nuclei from 800 animals, post-quality control filtering. Scale bar: 500 μm.
**(B)** UMAP of proboscis nuclei, colored by sample (female samples = 2, male samples = 2)
**(C)** UMAP of proboscis nuclei clustered using the Louvain algorithm (resolution = 1).
**(D)** UMAP of nuclei from all proboscis samples, colored and numbered by manual annotation using selected marker genes as listed in legend at the right of the figure panel. Categorization of cell type annotations represented by shaded gray headers. See Table S1 for gene IDs and thresholds
**(E-G)** Quantification of differences between male and female samples. Numbered and colored circles above (E) correspond to annotations in (D).
**(E)** Dot plot of number of differentially expressed genes between females and males in each cell type. Dots represent relative number of differentially expressed genes upregulated in male or female with a |log fold change| > 1 and false discovery rate < 0.05, determined by MAST on normalized expression and converted to $log_2$ fold change.
**(F)** Sexual abundance difference index from data in (G). Each cell type was categorized based on the following index: Female biased (purple) if 0.3 < abundance index; Neutral (grey) if -0.3 ≤ abundance index ≤ 0.3; Male biased (yellow) if abundance index < -0.3.
**(G)** Bar plot of relative cell abundance, represented as the percent of each cell type of all nuclei collected from each sex for tissue. Separated by female (purple) and male (yellow). Values for each replicate shown as dots, and standard error indicated by black brackets.
