## Supplemental Data S3 for "A single-nucleus transcriptomic atlas of the adult *Aedes aegypti* mosquito"

RNA *in situ* hybridization controls and probe information, related to Figures 2-4.

A

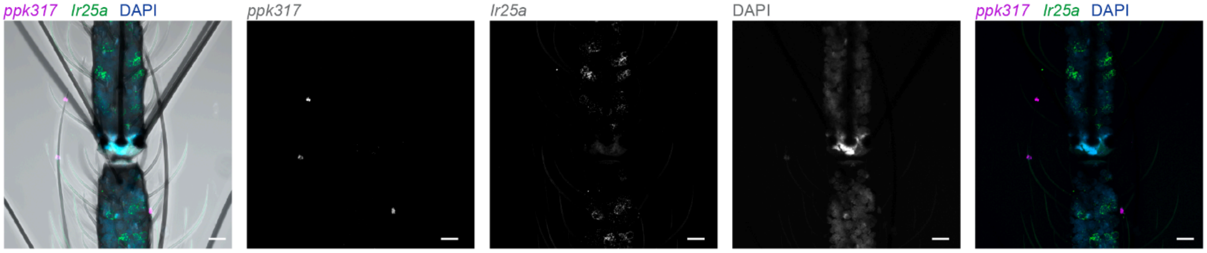


B

| Target Name | Gene | Organism | HCR Amplifier | Molecular Instruments Probe Lot Number |
|---|---|---|---|---|
| *ppk317* | *AAEL000873* | *Aedes aegypti* | B2 | RTR390 |
| *Ir41l* | *Ir41l* | *Aedes aegypti* | B2 | RTR716 |
| *Or82* | *Or82* | *Aedes aegypti* | B4 | RTR717 |
| *Or47* | *Or47* | *Aedes aegypti* | B5 | RTR718 |
| *Or3* | *Or3* | *Aedes aegypti* | B2 | RTR719 |
| *Ir25a* | *Ir25a* | *Aedes aegypti* | B3 | PRB511 |

C

| Target Name | Gene | Organism | HCR Amplifier | Individual probe sequence |
|---|---|---|---|---|
| *eya* | *AAEL019952* | *Aedes aegypti* | B1 | GAGGAGGGCAGCAAACGGAACCTGGGCTACGGATTTCCGATTTGA |
| | | | | TCCCCCTGAGATGGCTGCGTCAAGTTAGAAGAGTCTTCCTTTACG |
| | | | | GAGGAGGGCAGCAAACGGAACACCACTCATTCGTTTCTCTACCTG |
| | | | | CAACTGCGTCGCTTCGTTCTTCTTTTAGAAGAGTCTTCCTTTACG |
| | | | | GAGGAGGGCAGCAAACGGAATTGAAGAAGAAGAACGCATCCGCCA |
| | | | | AACCTGATCGCACTCCTCCAGATCGTAGAAGAGTCTTCCTTTACG |
| | | | | GAGGAGGGCAGCAAACGGAACTACTCGGTGGCAAGCAGCTGTCCG |
| | | | | AGCCGTTGGATAATGCAGCGTTGAGTAGAAGAGTCTTCCTTTACG |
| *AAEL001918* | *AAEL001918* | *Aedes aegypti* | B4 | CCTCAACCTACCTCCAACAATAGTTGTTGAAGCCGATGGTTCCAG |
| | | | | GACATGTCAGTTGATGGCAGATCAGATTCTCACCATATTCGCTTC |
| | | | | CCTCAACCTACCTCCAACAAATTGCTGTTGGCTGTCTGCTGATGT |
| | | | | CTTGCGAAGGGAAACGAATCCTAGAATTCTCACCATATTCGCTTC |
| | | | | CCTCAACCTACCTCCAACAATATGCCAACATCCCACGTCAAATGG |
| | | | | CGGATTTTTTCTTCGCTTTGCCTCCATTCTCACCATATTCGCTTC |
| | | | | CCTCAACCTACCTCCAACAACAGGCTGTTCCACTGGCAAATCTAT |
| | | | | GTTTCTTCCGGAGCAAGCTGTTGAAATTCTCACCATATTCGCTTC |
| *ana* | *AAEL007208* | *Aedes aegypti* | B2 | CCTCGTAAATCCTCATCAAACACCCCGCACAGCGCCATAGTTCCA |
| | | | | ATGGCCACATCTGAGGCGTACGGGGAAATCATCCAGTAAACCGCC |
| | | | | CCTCGTAAATCCTCATCAAAGCGCGTTGCATGTTCTTCTGGGCGG |
| | | | | TTGCTTGATGGCCATCGGAGCCCCGAAATCATCCAGTAAACCGCC |
| | | | | CCTCGTAAATCCTCATCAAATGCCGAATTAACCGAAGTGTAAGGC |
| | | | | CGCCGATCATGTACAACCTCAACATAAATCATCCAGTAAACCGCC |
| | | | | CCTCGTAAATCCTCATCAAAAACACGAATCATCTGCTCGGCGGGG |
| | | | | TCTTGCGCCAGTACACGGAAACGGTAAATCATCCAGTAAACCGCC |
| *beta2-tubulin* | *AAEL019894* | *Aedes aegypti* | B2 | CCTCGTAAATCCTCATCAAACCTCGGTGGTGAAAATCGAAGTGAA |
| | | | | AATGGTTCCAAATCACGTCCGCGTTAAATCATCCAGTAAACCGCC |
| | | | | CCTCGTAAATCCTCATCAAAGGATCCATTCGACGAAGTAGCTACT |
| | | | | CAGACGGCCGTCTTCACATTGTTCGAAATCATCCAGTAAACCGCC |
| | | | | CCTCGTAAATCCTCATCAAATGATCCGCTCCAGCTGCAAATCGCT |
| | | | | CCGGTAGCCTCGTTGTAGTACACGTAAATCATCCAGTAAACCGCC |
| | | | | CCTCGTAAATCCTCATCAAAGGGAATGGAACCATATTGACTGCCA |
| | | | | GCCGGTCATGAAAAAGTGCAGCCGCAAATCATCCAGTAAACCGCC |
| *Vas* | *AAEL004978* | *Aedes aegypti* | B4 | CCTCAACCTACCTCCAACAAGCTTCGTTGAAAATCTGTAGCGCCA |
| | | | | AACGGTTCCAAGCGCAAACTTGCGGATTCTCACCATATTCGCTTC |
| | | | | CCTCAACCTACCTCCAACAAAAATCGCTACGGCCGCCTCCTCGTC |
| | | | | TTCTCCATCATTGTCGCCACCGCCGATTCTCACCATATTCGCTTC |
| | | | | CCTCAACCTACCTCCAACAATTCGTTGTCGCCTCCGCCGAAGTCG |
| | | | | CACCACGACCGAATCCGTTTTCTCCATTCTCACCATATTCGCTTC |
| | | | | CCTCAACCTACCTCCAACAAGGGTTCTCTCCCGTTACGTTTACTT |
| | | | | GCCGAACGACTTGATTGGGCTAGGAATTCTCACCATATTCGCTTC |

**Data S3. RNA *in situ* hybridization controls and probe information, related to Figures 2-4.**
**(A)** Maximum-intensity projection of whole-mount female antennae with RNA *in situ* hybridization of *ppk317* probe (magenta), *Ir25a* (green) and nuclear staining (DAPI). Scale bar: 10 μm, related to Figure 3.
**(B) Antennae RNA *in situ* hybridization probe information.** Table includes targeted gene names, gene IDs, organism, HCR Amplifier, and probe lot numbers for ordering, related to Figures 3 and 4. To order probe sets, contact Molecular Instruments (https://www.molecularinstruments.com/) and provide the probe lot number.
**(C) Testis RNA *in situ* hybridization probe information.** Table includes targeted gene names, gene IDs, organism, HCR Amplifier, and probe sequences, related to Figure 2.
