## Supplemental Data S4 for "A single-nucleus transcriptomic atlas of the adult *Aedes aegypti* mosquito"

Filtering and characterization of *nompC (AAEL019818)*-negative sensory neurons, related to Figure 5.

Index:

**(Data S4.1)** Sensory neuron filtering and analysis, related to Figures 5 and S7.
**(Data S4.2)** Neuropeptide receptor and synthesis gene analysis in antenna, and characterization and sexual dimorphism in *nompC* (*AAEL019818*)-negative sensory neurons, related to Figures 4, 5, S6, and S7.

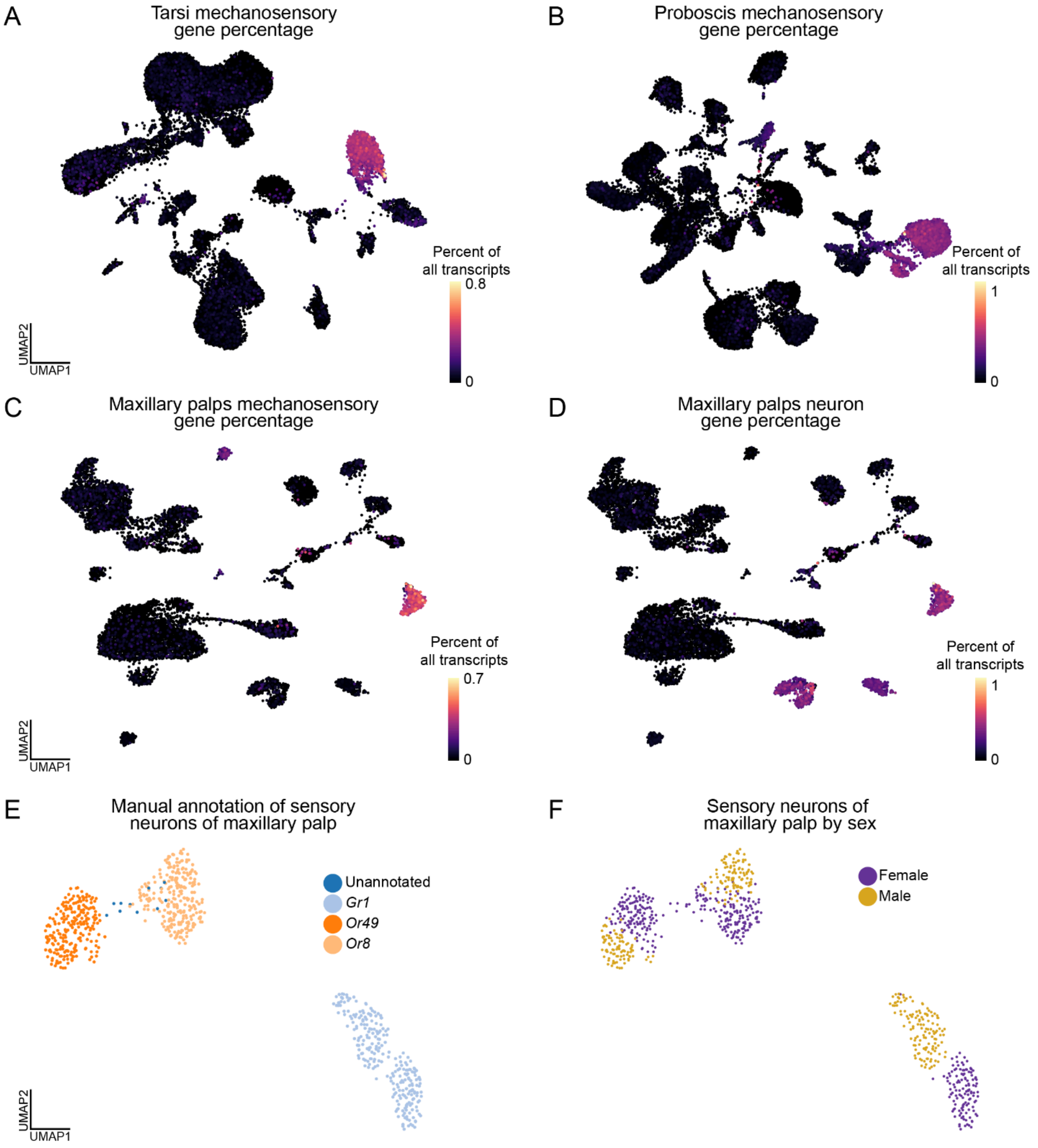


**Data S4.1. Sensory neuron filtering and analysis, related to Figures 4 and 5.**
**(A-C)** UMAP of neurons from tarsi (A), proboscis (B), and maxillary palps (C) data with expression of *nompC* (*AAEL019818*) as percent of all transcripts in each cell.
**(D)** UMAP of all maxillary palp cells with expression of neuron marker genes (*Syt1 (AAEL000704), nSyb (AAEL024921), brp (AAEL018153), CadN (AAEL000597)*) as percent of all transcripts in each cell.
**(E-F)** UMAP of *nompC*-negative sensory neurons from maxillary palps, colored by manual annotation (E) and sample (F) (female = 1, male = 1). Female sample previously published[36].

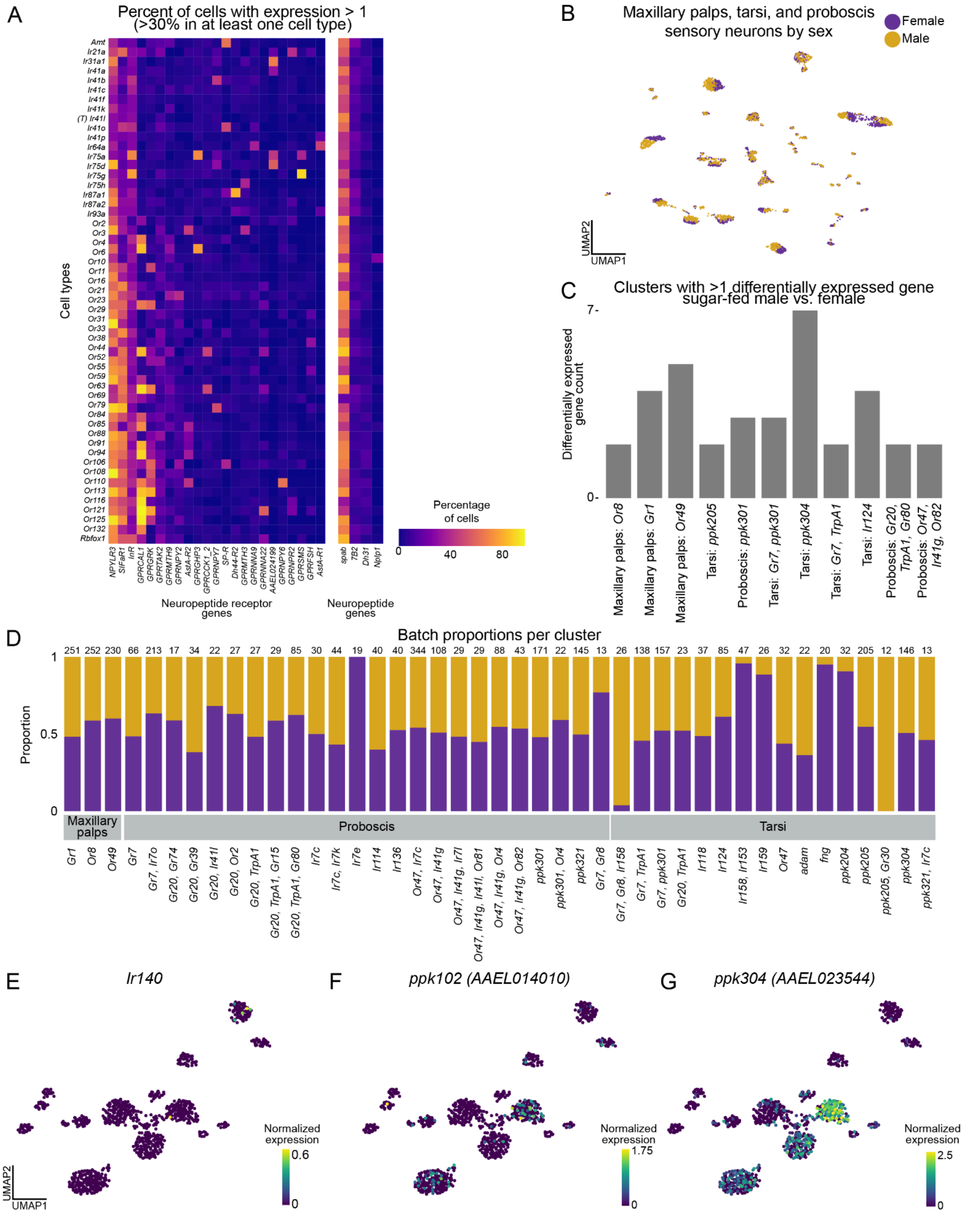
A
Percent of cells with expression > 1 (>30% in at least one cell type)
Cell types
Neuropeptide receptor genes
Neuropeptide genes
Percentage of cells
B
Maxillary palps, tarsi, and proboscis sensory neurons by sex
Female
Male
UMAP1
UMAP2
C
Clusters with >1 differentially expressed gene sugar-fed male vs. female
Differentially expressed gene count
D
Batch proportions per cluster
Proportion
Maxillary palps
Proboscis
Tarsi
E
Ir140
F
ppk102 (AAEL014010)
G
ppk304 (AAEL023544)
Normalized expression

**Data S4.2. Neuropeptide receptor and synthesis gene analysis in antenna, and characterization and sexual dimorphism in nompC-negative sensory neurons, related to Figures 4, 5, S6, and S7.**

**(A)** Heatmap of expression of neuropeptide receptor genes (left) and neuropeptide synthesis genes (right) within manually annotated olfactory sensory neurons (*nompC*-negative) in the antenna. Color scale indicates percentage of cells expressing a gene above threshold (normalized expression value of 1). Sensory genes are indicated in columns and cell types indicated in rows. Genes were included if they were expressed above threshold in over 20% of cells in at least one cell type. Genes filtered from lists in Table S1. Normalized expression is *ln([(raw count/total cell counts)*median total counts across cells]+1)*.

**(B)** UMAP of combined *nompC*-negative sensory neurons from maxillary palps, tarsi and proboscis samples, colored by sex.

**(C)** Bar plot of number of differentially expressed genes (DEGs) between females and males in each olfactory sensory neuron cell type. Clusters shown contain at least 1 DEG with a |log fold change| >1 and false discovery rate 0.05 (determined by MAST on normalized expression). For more information on DEGs, see Table S2.

**(D)** Proportion plot for annotated *nompC*-negative sensory neurons from maxillary palps, tarsi and proboscis. Bar plot of proportion cells from each sex that makes up each cell type. Numbers above each bar indicate the number of cells in that cluster. Cell types that are more than 70% from a single sex indicated in red.

**(E-G)** In tarsi *nompC*-negative sensory neurons, UMAP of normalized gene expression of *Ir140* (E), *ppk102 (AAEL014010)* (F), and *ppk304 (AAEL023544)* (G). Each end of color bar trimmed 0.1% for visibility.
