## Supplemental Data S5 for "A single-nucleus transcriptomic atlas of the adult *Aedes aegypti* mosquito"

*Aedes aegypti* Mosquito Cell Atlas Consortium Members and Funding

***Aedes aegypti* Mosquito Cell Atlas Consortium** (in alphabetical order):
Joshua X. D. Ang[1], Igor Antoshechkin[2], Yu Cai[3,4], Fangying Chen[5], Yen-Chung Chen[6], Julien Devilliers[7], Linhan Dong[8], Roberto Feuda[7], Paolo Gabrieli[9], Artyom Kopp[10], Hyeogsun Kwon[11], Hsing-Han Li[5], Tzu-Chiao Lu[12,13], Thalita Lucio[14,15], João T. Marques[16], Marcus F. Oliveira[14,15], Roenick P. Olmo[16], Umberto Palatini[17,+], Zeaan M. Pithawala[18], Julien Pompon[19], Yan Reis[14,15], João Rodrigues[14,15], Ryan C. Smith[11]

[1]The Department of Biology, University of York, Wentworth Way, York, YO10 5DD, UK
[2]Division of Biology and Biological Engineering (BBE), California Institute of Technology, Pasadena, CA 91125, USA
[3]Temasek Life Sciences Laboratory, 1 Research Link, National University of Singapore, Singapore 117604
[4]Department of Biological Sciences, National University of Singapore, Singapore 117543
[5]School of Biological Sciences, Department of Cell and Developmental Biology, University of California, San Diego, La Jolla, CA 92093, USA
[6]Department of Biology, New York University, New York, NY 10003, USA
[7]Department of Genetics, Genomics and Cancer Sciences, University of Leicester, Leicester, LE1 7RH, UK
[8]Department of Biological Sciences, Columbia University, New York, NY 10027, USA
[9]Department of Biosciences, University of Milan, Milan, 20133, Italy
[10]Department of Evolution and Ecology, University of California Davis, Davis, CA 95616, USA
[11]Department of Plant Pathology, Entomology and Microbiology, Iowa State University, Ames, IA 50011, USA
[12]Huffington Center on Aging, Baylor College of Medicine, Houston, TX 77030, USA
[13]Department of Molecular and Human Genetics, Baylor College of Medicine, Houston, TX 77030, USA
[14]Institute of Medical Biochemistry Leopoldo de Meis, Universidade Federal do Rio de Janeiro, Rio de Janeiro, RJ, CEP 21941-590, Brazil
[15]Instituto Nacional de Ciência e Tecnologia em Entomologia Molecular (INCT-EM), Rio de Janeiro, Brazil
[16]Inserm U1257, CNRS UPR9022, Université de Strasbourg, 67084 Strasbourg, France
[17]Laboratory of Neurogenetics and Behavior, The Rockefeller University, New York, NY 10065, USA
[18]Department of Epidemiology of Microbial Diseases, Yale School of Public Health, New Haven, CT 06510, USA

[19]MIVEGEC, Univ. Montpellier, IRD, CNRS, INRAe, Montpellier, France
[+]Present address: Department of Biology and Biotechnology, University of Pavia, Pavia 27100, Italy

***Aedes aegypti* Mosquito Cell Atlas Consortium members support**: J.X.D.A. by EPSRC (EP/X025535/1). Y.C. by Temasek Life Sciences Laboratory. J.D. by PhD scholarship from the College of Life Science (University of Leicester). T.L., Y.R., J.R., and M.F.O. by CNPq (308629/2021-3) and Coordenação de Aperfeiçoamento de Pessoal de Nível Superior-Brasil (CAPES)-Finance Code 001. Y.-C.C. by New York University MacCracken Fellowship, NYSTEM institutional training grant (Contract #C322560GG), and Scholarship to Study Abroad from the Ministry of Education, Taiwan. U.P. is supported by EMBO Long-Term Fellowship (ALTF 664-2022), Human Frontier Science Program (LT 0012-2023L), and Stavros Niarchos Foundation (SNF) Institute for Global Infectious Disease Research. J.P. by ANR-20-CE15-006. A.K. by NIH R35GM122592. R.F. by Royal Society University Research Fellowship URF\R\221011. R.C.S by NIH R21AI166857.
